# Codon optimality modulates cellular stress and innate immune responses triggered by exogenous RNAs

**DOI:** 10.1101/2024.11.26.625518

**Authors:** Chotiwat Seephetdee, Nada Bejar, Eric Chau, Thy Y. Nguyen, Biana Godin, Daniel L. Kiss

## Abstract

Despite advancements in RNA sequence design, evidence regarding the preferential use of synonymous codons on cellular stress and innate immune responses is lacking. To this end, we developed a new codon optimality formula to re-engineer the coding sequences of three luciferase reporters. We demonstrate that mRNAs enriched in optimal codons elicited dramatic increases in luciferase activities compared to less optimal sequences both *in vitro* and *in vivo*. Notably, transfecting low optimality test RNA suppresses the translation of co-transfected control mRNAs in dual reporter assays. Further, low optimality mRNAs activated innate immune pathways and the phosphorylation of the translation initiation factor eIF2α, a central event of the integrated stress response. eIF2α phosphorylation was suppressed by a GCN2 inhibitor, but not by other eIF2α kinase inhibitors. Using nucleoside-modified or circular RNAs also fully or partially abrogated these responses. Finally, optimal, but not non-optimal, circular RNAs have enhanced RNA lifespan and duration of protein expression. Our results show that RNA sequence, composition, and structure all govern RNA translatability. Further, RNA sequences with poor codon optimality are immunogenic and induce cellular stress. Together, we show that RNA coding sequence design is a key consideration for both mRNA and circular RNA therapeutics.

## Introduction

As demonstrated by the efficacy of messenger RNA (mRNA) vaccines against severe coronavirus disease 2019 (COVID-19), mRNA has emerged as a crucial therapeutic modality in medicine [1, 2]. With applications in cancer immunotherapy, regenerative medicine, and the treatment of genetic disorders, mRNA therapies are expanding to indications beyond infectious diseases [3–5]. Several innovative approaches have been shown to advance mRNA technology by reducing immunogenicity and/or by enhancing mRNA stability and translation efficiency. Substitution of uridine (U) with *N*^1^-methylpseudouridine (m^1^Ψ) minimizes inherent mRNA immunogenicity by dampening recognition by Toll-like receptors (TLRs) of the innate immune system while also markedly increasing protein expression [6–8]. Coding sequence (CDS) and untranslated region (UTR) optimization has also been shown to enhance protein production [9–11]. The development of circular RNA (circRNA) and self-amplifying mRNA, which prolong RNA lifespan and translation durability due to exonuclease resistance and self-replication ability, respectively, are major advances that extend the pharmacokinetic profiles of those RNA platform technologies [12–16]. Further, RNA circularization also decreases the immunogenicity of unmodified mRNAs, in part due to TLR evasion [13, 17]. Recently, through chemo-topological engineering efforts, mRNA lifetime and translation capacity have also been augmented by multimerizing the poly(A) tail and by chemical modifications of the 5’ end’s 7-methylguanosine cap structure [18, 19]. These continuous improvements evince the potential for building better mRNA therapeutics.

Despite the advancements observed in mRNA as a platform technology, the impact of open reading frame (ORF) sequences on regulating gene expression remains under-reported in the published literature, especially in the context of protein-coding RNA therapeutics. For instance, synonymous codon usage is one way to modulate gene expression by regulating translation, mRNA stability, and mRNA localization [20, 21]. The degeneracy in the genetic code means that multiple distinct triplet codons code for most amino acids. During translation elongation, the ribosome recognizes synonymous codons differently, leading to local changes in translation rates within the CDS of an mRNA. The stochastic recognition of the codon at the ribosome A-site by charged tRNAs, which contributes to ribosome elongation dynamics, is also influenced by, among other things, the abundance of the cognate tRNA and wobble (non-Watson-Crick) base pairing. New evidence has also implicated selected arginine tRNAs for co-translational P-site tRNA-mediated mRNA decay pathway mediated by CNOT3 [22]. A sustained series of non-optimal codons is thought to locally slow or pause translation elongation, sometimes leading to ribosome collisions and subsequent mRNA and/or ribosome degradation [21, 23, 24]. A growing body of work has also highlighted the implication of ribosome collisions on integrated stress response and innate immune activation [25, 26]. Therefore, it is reasonable to postulate that codon optimality of exogenous mRNAs is relevant to these cellular events as well.

Codon optimization has become a common practice in heterologous gene expression [27]. As a result, several indices have been employed to optimize codon usage including the codon adaptation index (CAI) [28]. The CAI provides an approximate indication of heterologous gene expression level by scoring the codon usage frequency of a given gene compared to that of a reference organism [29, 30]. Although based on genomic sequences, the CAI score can also be used to *approximate* codon optimality. For example, the COVID-19 mRNA vaccines represent an obvious example of the superiority of optimal codon usage in an RNA therapeutic. The CAI score of the native SARS-CoV-2 Spike protein is 0.67 while CAI scores of mRNA-1273 and BNT162b2 vaccines are 0.98 and 0.95, respectively [31–33]. Since mammals have a bias for G/C at wobble positions, codon optimization for human therapeutics tends to increase GC content, thereby reducing U content [34]. As a result, high GC content and strong RNA secondary structure can influence both translation rate and efficiency [35]. Furthermore, unmodified mRNAs with reduced U content could also help reduce innate immune activation, thereby increasing protein expression [9, 10].

In this work, we hypothesized that the codon optimality of exogenous mRNAs can influence or modulate cellular stress and/or innate immune responses. We re-engineered the coding sequence of three luciferase genes, including Nanoluciferase (NLuc), *Renilla* luciferase (RLuc), and Firefly luciferase (FLuc) using a new codon optimality formula to increase the number of optimal codons. In all cases, the engineered, high optimality reporter mRNAs achieved superior luciferase activity compared to less optimal sequences. Moreover, we observed that the optimality of the exogenous test mRNAs also affected the expression of the transfection control mRNAs. In particular, poorly optimal mRNAs suppressed the expression of the independent control reporter compared to highly optimal mRNAs. We further interrogated the effects of mRNA codon usage on possible mechanisms contributing this observation. Specifically, we show that low optimality mRNAs led to a substantial increase in the phosphorylation of eukaryotic translation initiation factor 2 α (eIF2α), which is a hallmark of integrated stress response that globally suppresses translation initiation. Our results suggest that non-optimal codon usage in mRNAs induced eIF2α phosphorylation primarily through the activity of the eIF2α kinase GCN2. Intriguingly, we demonstrate that substituting non-optimal codons with synonymous, optimal codons significantly reduced eIF2α phosphorylation levels. Further, we show that codon optimality also contributes to the immunogenicity of mRNA. Our data reveal that mRNAs with poor optimality strongly induced the expression of innate immunity genes which is markedly reduced both by using m^1^Ψ or by substituting an mRNA with an increased prevalence of optimal codons. Lastly, by utilizing destabilized luciferase reporters, we demonstrate that only circRNAs with highly optimal coding sequences have an extended lifespan. Taken together, our findings indicate that engineering coding sequences to increase codon optimality provides an additional therapeutic mRNA design approach to modulate or minimize cellular stress and innate immune responses.

## Results

### mRNAs with optimal sequences produce more protein

To increase reporter protein expression from exogenous mRNAs, we developed and used a new codon optimality formula to re-engineer the coding sequences of NLuc, RLuc, and FLuc. For ease of detection, FLuc and NLuc were engineered with 3x FLAG tags (with codon distributions similar to their fused reporter protein) at their C-terminus. Overall, the optimal reporters have increased GC content, reduced U content, and a lower minimum free energy suggesting a more stable tertiary structure (Figure S1A) [35]. We *in vitro* transcribed both non-optimal and optimal linear mRNAs encoding the individual reporters with standard ribo-nucleotides (Figure S1). To compare the expression of non-optimal and optimal reporter mRNAs *in vivo*, we injected BALB/c mice with FLuc reporter mRNAs encapsulated in lipid nanoparticles (LNP). Mice were imaged every 24 hours after LNP-encapsulated mRNA administration using an *In Vivo* Imaging System (IVIS) (Figures 1A and S2). The luminescence from optimal FLuc mRNA was higher than the signal from non-optimal FLuc mRNA at every time point (Figures 1A and S2). Although there is no consensus regarding a unified codon optimality score, we used the human CAI score as an approximate measure of codon optimality [28, 30]. Depending upon the reporter, the resulting codon selections for each luciferase reporter corresponded to a CAI score of 0.98, 0.97, and 0.97 for the highly optimal coding sequences of NLuc (ΔCAI = +0.25), RLuc (ΔCAI = +0.31), and FLuc (ΔCAI = +0.24), respectively (Figure S1A).

**Figure 1.**
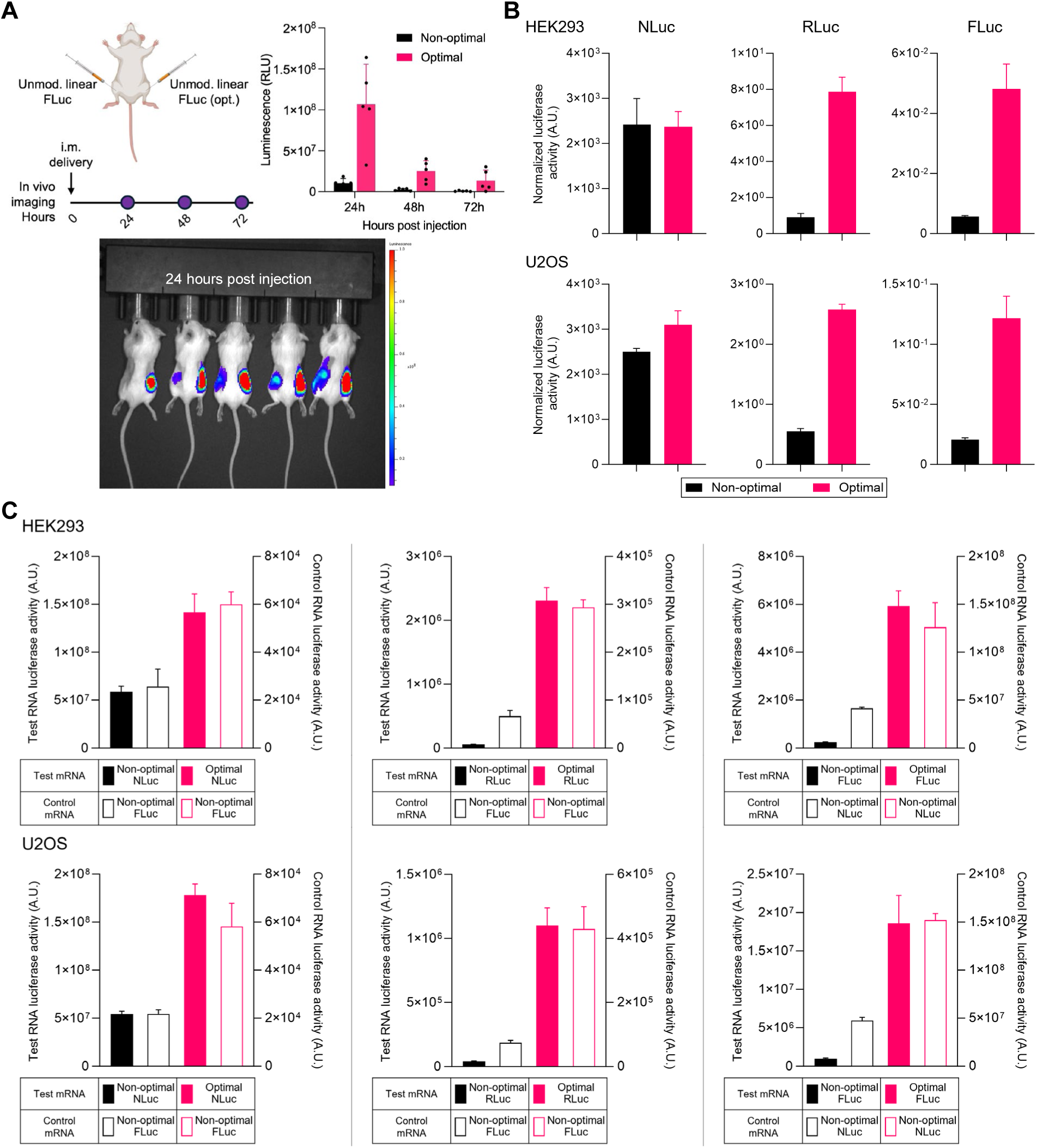
The codon optimality of test mRNAs affects expression of both test and control mRNAs. **(A)** Mice were injected with unmodified non-optimal (left flank) and optimal (right flank) firefly luciferase (FLuc) reporter mRNAs encapsulated in lipid nanoparticles (LNP) and monitored for 3 days. FLuc bioluminescence was captured at 24, 48, and 72 h using IVIS and mean luminescence +Std Dev for each day are shown. IVIS images at 24 h after FLuc mRNA-LNP injection are shown. **(B)** Normalized luciferase activity from HEK293 or U2OS cells lysed 24 h after co-transfection with mRNAs encoding either optimal or non-optimal forms of Nanoluciferase (NLuc), *Renilla* luciferase (RLuc), or firefly luciferase (FLuc) reporters. Protein expression was measured by dual-luciferase assays, and the luminescence of optimal or non-optimal test reporters were normalized to control reporter luminescence. **(C)** Codon optimality of test mRNAs shows differences in protein levels of both test and transfection control mRNAs. Dual-luciferase assays from HEK293 or U2OS cells lysed 24 h after transfection with mRNAs encoding either optimal or non-optimal forms of NLuc, RLuc, or FLuc test reporters (left axis, solid bars) along with non-optimal luciferase transfection control RNA (right axis, empty bars). Each experiment consists of a color-matched solid bar (read on the left Y-axis) and the empty bar (read on the right Y-axis) directly adjacent to it. Data are shown as the mean +Std Dev of four independent experiments.

With regards to the CAI scores of our reporters, it is often assumed that the CAI scores for natural genes range from 0.0 to 1.0. In fact, sequences with such ultra-low or super high CAI scores are exceedingly rare in nature. To demonstrate this point, we calculated the CAI scores of several abundant housekeeping genes, which we presumed would have high CAI scores, several genes whose proteins are known to undergo aberrant folding, which we reasoned would have lower CAI scores to facilitate co-translational protein folding, and several other RNA or disease-linked genes. We also made sure to include short, intermediate-length and long ORFs as well. In humans, most genes ‒even those with known co-translational folding defects‒ have CAI scores ranging between ∼0.7 and 0.9 (Table S1). Notably, the CAI scores of the high and low optimality reporters (which for simplicity, we will call optimal and non-optimal respectively from hereon) exceed those of endogenous human genes, while the CAI score for non-optimal reporters align with low-optimality natural genes (Figure S1A and Table S1).

Exemplifying the disagreement in the field regarding which codons are most optimal, as shown in Table S2, with ∼220 and ∼300 different codon selections merely by count, our codon optimality formula generates starkly different sequences for 550 codon open reading frame of firefly luciferase than the formulas from two other well-regarded teams [36, 37]. Critically, the codon selection differences shown in the table only account for the most favored codon used and do not account for the positional choices for where each codon was employed [36, 37]. To further illustrate the point, the CAI scores of poorly and highly optimal Fluc mRNA in Barrington et al. are 0.62 and 0.75 (compared to ours at 0.73 and 0.97) respectively[36]. Since these sequence designs are so divergent, although they encode the identical protein, it is exceedingly unlikely that the cellular RNA surveillance machinery will survey and act upon our coding sequences as was observed in those earlier studies [36, 37].

### The optimality of test mRNAs affects protein output from both test and control mRNAs *in vitro*

We co-transfected reporter mRNAs with control mRNAs into multiple cell lines for dual luciferase assays. To our surprise, while the optimal RLuc and FLuc reporters had 6-10-fold increases in luciferase activity when we normalized the signals to the co-transfected control RNA, the optimal NLuc reporters showed no difference in luciferase activity in HEK293 cells and only had a marginal increase on luciferase activity in U2OS cells (Figure 1B). However, upon closer examination of the raw, non-normalized data, we noticed that all optimal reporters showed large increases in luciferase activities when compared to non-optimal mRNAs in all four (HEK293, U2OS, NIH/3T3, and BJ) cell types assayed (Figures 1C, left Y-axis, and S3A). Intriguingly, we found that the co-transfected control reporter in non-optimal test reporter-transfected cells showed substantially lower levels of luciferase activity driven from the control mRNA compared to the same mRNA in cells transfected with optimal test reporters (Figure 1C, right Y-axis). In other words, the optimality of the test mRNA was exerting an effect on and changing the signal returned by the control mRNA. This made normalizing the data difficult since the control RNA is assumed to remain constant. Further, the distortions of signals from the control RNA held true regardless of which pair of luciferase transcripts were used. To rule out that this effect was caused by experimental artifacts within dual luciferase assays, we validated this observation by co-transfecting the optimal or non-optimal luciferase reporters with mRNA encoding an EGFP reporter protein (Figure S3B). In agreement with the dual luciferase assays, cells co-transfected with non-optimal luciferase mRNAs showed lower EGFP fluorescence as measured by an Incucyte live cell imaging system than cells transfected with optimal luciferase reporters (Figure S3B). To further exclude the possibility that this observation was caused by the effects of mRNA codon optimality on cell health, viability of the transfected cells was evaluated using CellTiter-Glo assay. Cells transfected with non-optimal or optimal RLuc mRNAs showed similar viability at 24 and 48 hours after transfection (Figure S4). Together, these data show that the codon optimality of experimental mRNAs influences translation from the co-transfected control mRNAs.

### Unmodified linear mRNAs with poor codon optimality induce eIF2α phosphorylation in a predominantly GCN2-dependent manner

In yeast, reporter mRNAs with non-optimal codons can induce the phosphorylation of translation initiation factor eIF2α in a CDS length-dependent manner [38]. The effect that codon usage has on eIF2α phosphorylation resembles the effect of ribosome stalling induced by translation elongation inhibitors [38–40]. Phosphorylated eIF2α leads to global reduction in translation initiation and is an integral part of integrated stress response pathways [25, 41]. We compared levels of eIF2α phosphorylation in HEK293 cells individually transfected with unmodified linear mRNAs encoding non-optimal or optimal reporters by western blot analysis. Interestingly, cells transfected with non-optimal reporters exhibited significant increases in eIF2α phosphorylation over optimal mRNAs (Figure 2A). The increased levels of phosphorylated eIF2α are consistent with globally reduced (but not blocked) translation initiation and could explain the reduced control reporter expression observed in cells transfected with non-optimal mRNA. To elucidate which stress-induced kinase(s) is/are responsible for the elevated levels of phosphorylated eIF2α, we pre-treated HEK293 cells with the inhibitors of protein kinase R (PKR) (C16), PKR-like endoplasmic reticulum kinase (PERK) (AMG-44), and general control nonderepressible 2 (GCN2) (A-92) prior to transfection of the non-optimal or optimal FLuc reporter mRNAs and conducted western blot analysis to assay eIF2α phosphorylation (Figure 2B, top) [42–44]. The blots show that the PKR inhibitor had a minimal effect, while the PERK inhibitor caused no change in eIF2α phosphorylation, meaning that neither the presence of double-stranded RNA (dsRNA) nor the accumulation of misfolded proteins and/or ER stress drove the observed stress response (Figure 2B). In fact, the eIF2α phosphorylation observed in cells transfected with non-optimal mRNAs was only suppressed by A-92. Further, as shown before, optimal mRNAs cause a small increase in eIF2α phosphorylation, but even that is suppressed by A-92 treatment. These data suggest that, in our assays, GCN2 is the major stress-induced kinase triggered by non-optimal mRNA (Figure 2B).

**Figure 2.**
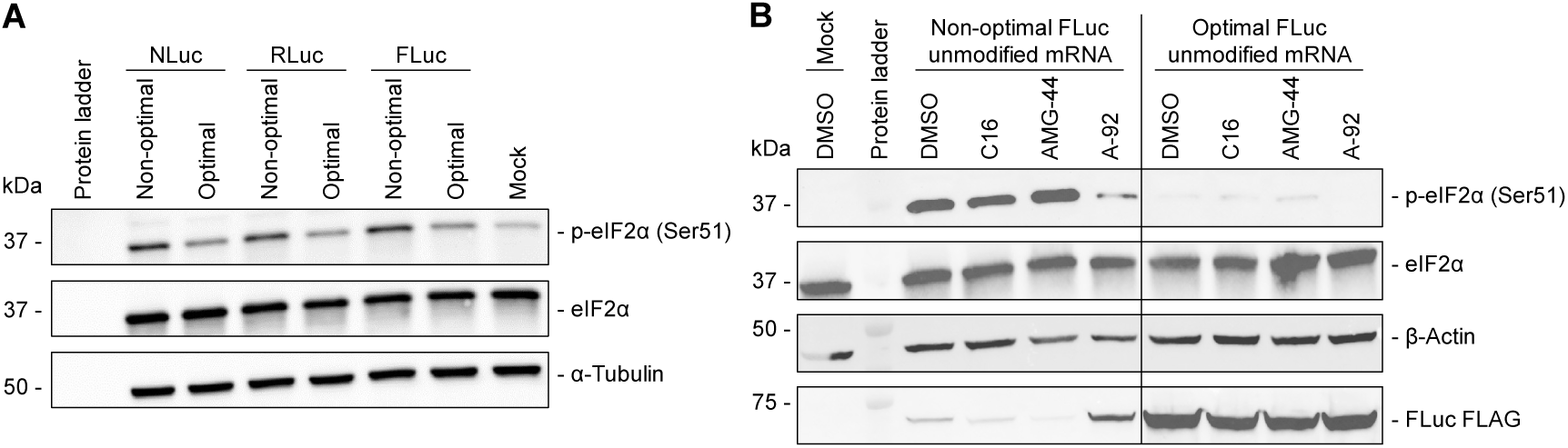
Poorly optimal coding sequences cause eIF2α phosphorylation in a GCN2-dependent manner. **(A)** HEK293 cells were harvested 16 h after transfection with mRNAs encoding either non-optimal or optimal forms of three luciferase reporters. Cell lysates were assayed by immunoblotting with antibodies targeting either phosphorylated eIF2α, total eIF2α, or α-tubulin (as a loading control). **(B)** HEK293 cells were pre-treated with DMSO, C16, AMG-44, or A-92 as detailed in methods, and were transfected with either non-optimal or optimal FLuc mRNA. After 16 h, cells were lysed and lysates were assayed by immunoblotting with antibodies targeting either phosphorylated eIF2α, total eIF2α, β-actin, and the FLAG epitope.

As eIF2α phosphorylation inhibits global protein synthesis, and we saw that A-92 and C16 (to a much smaller extent) reduced eIF2α phosphorylation, we reasoned that the inhibitors could influence protein production from the transfected mRNAs. We used western blotting with anti-FLAG antibodies to assay translation from non-optimal or optimal FLuc RNA in the presence or absence of the three inhibitors. In cells transfected with non-optimal RNA, treatment with A-92 resulted in a noticeable increase in FLuc protein levels (Figure 2B, bottom), indicating that GCN2 inhibition boosted translation from our non-optimal mRNA. This effect was specific to the A-92 and non-optimal mRNA combination, as no increase in protein expression was observed with the other inhibitors, or when the inhibitors were added to cells transfected with optimal mRNAs.

### Unmodified linear mRNAs with poor codon optimality activate innate immunity genes

Since exogenous unmodified linear mRNA is known to be immunogenic, we next asked whether differences in reporter protein expression were partially caused by reduced immunogenicity associated with optimal codon usage. We used RT-qPCR to assess the expression profiles of a panel of genes known to be activated by different innate immune responses. In total, we report results from a dozen innate immunity genes, including interferon-stimulated genes (ISGs), antiviral response genes, inflammatory cytokines, and innate immune regulators in U2OS cells transfected with unmodified linear mRNAs encoding optimal or non-optimal reporters. As expected, transfection of reporter mRNAs led to increased expression of innate immune response genes relative to mock transfection (Figure 3). Notably, optimal mRNAs resulted in a significant suppression of innate immunity gene activation compared to the non-optimal reporters. These data show that poor codon optimality can contribute to the immunogenicity of exogenous mRNAs.

**Figure 3.**
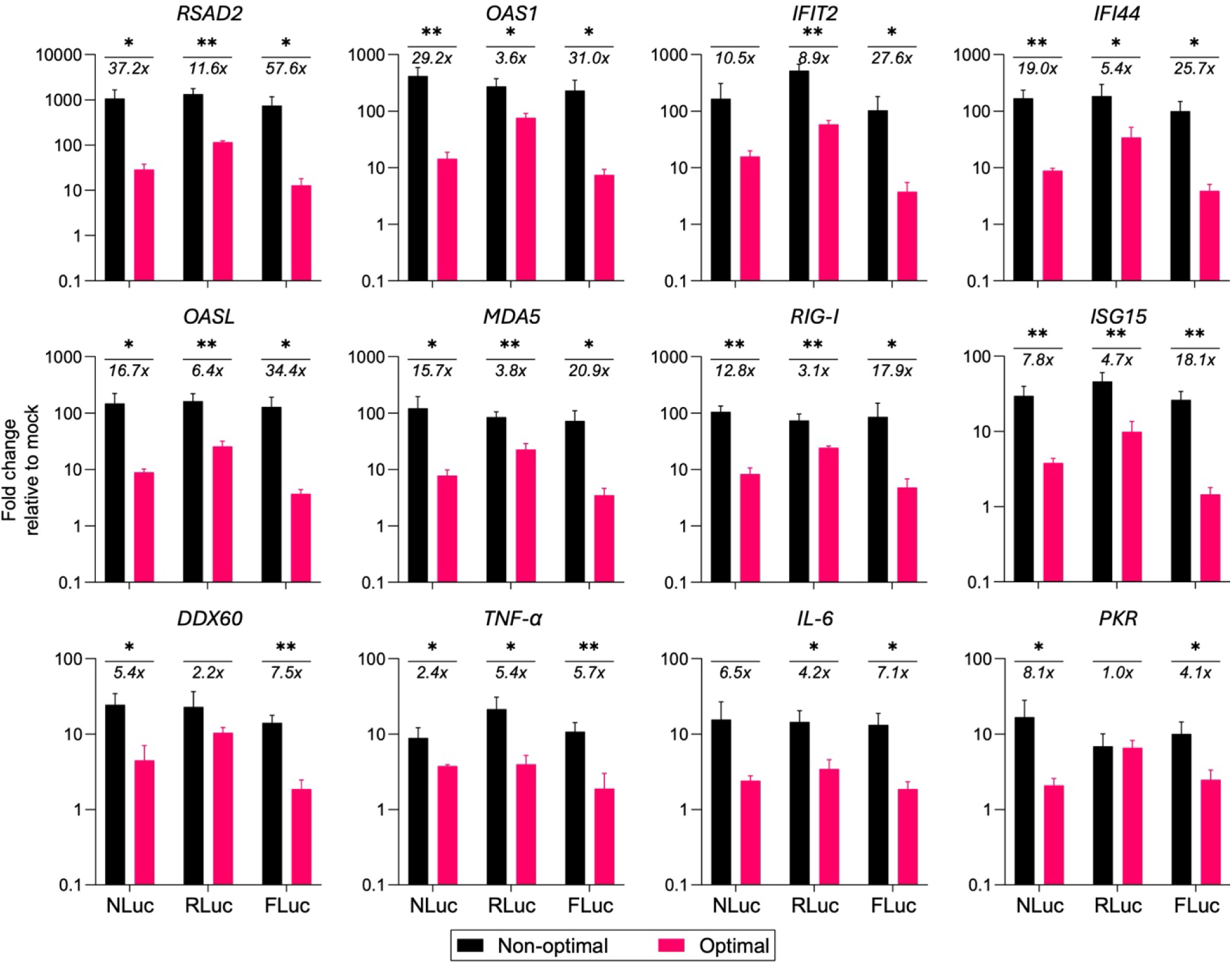
High optimality sequences reduce innate immunity gene activation by unmodified linear mRNA. U2OS cells were transfected with reporter mRNAs encoding either non-optimal or optimal luciferase reporters for 16 h. RNA was extracted from cell lysates and relative induction of innate immunity genes were measured by RT-qPCR, with relative fold changes normalized to expression levels in mock transfected cells. Data are shown as the mean +Std Dev of three independent biological replicates. Student’s t-test was used to evaluate the significance of the results, and \**p* < 0.05, \*\**p* < 0.01.

### Optimal codons increase reporter protein expression from linear and circular RNAs

mRNA engineering approaches, including chemical modification of nucleotides and RNA circularization, have also been shown to reduce immunogenicity while increasing the efficiency and durability of protein expression [6, 12]. We next evaluated the level and duration of protein expression from our optimality reporters delivered using three different RNA therapy platform technologies (unmodified mRNA, modified mRNA, and circRNA). Consistent with the mRNA vaccines for COVID-19, modified linear mRNAs were made by complete substitution of U with m^1^Ψ [32, 33]. circRNAs containing a CVB3 IRES driving the translation of an optimal or non-optimal reporter CDS were generated by adapting a previously published permuted intron-exon (PIE) strategy [12]. Transfection into HEK293 and U2OS cells showed that optimal reporter mRNAs exhibited higher luciferase activities compared to non-optimal counterparts across both RNA types (Figures 4A and 4B). We also observed different expression profiles and durability of protein expression when comparing linear and circular RNAs in which circRNAs predominantly showed lower peak level protein expression but maintained protein expression longer than linear mRNAs. The expression level and durability of expression also varied depending upon the reporter protein, but optimal sequences always outperformed non-optimal sequences for both parameters. Further, non-optimal circRNAs generally had similar expression profiles and RNA half-lives as non-optimal mRNAs (Figures 4A and 4B).

**Figure 4.**
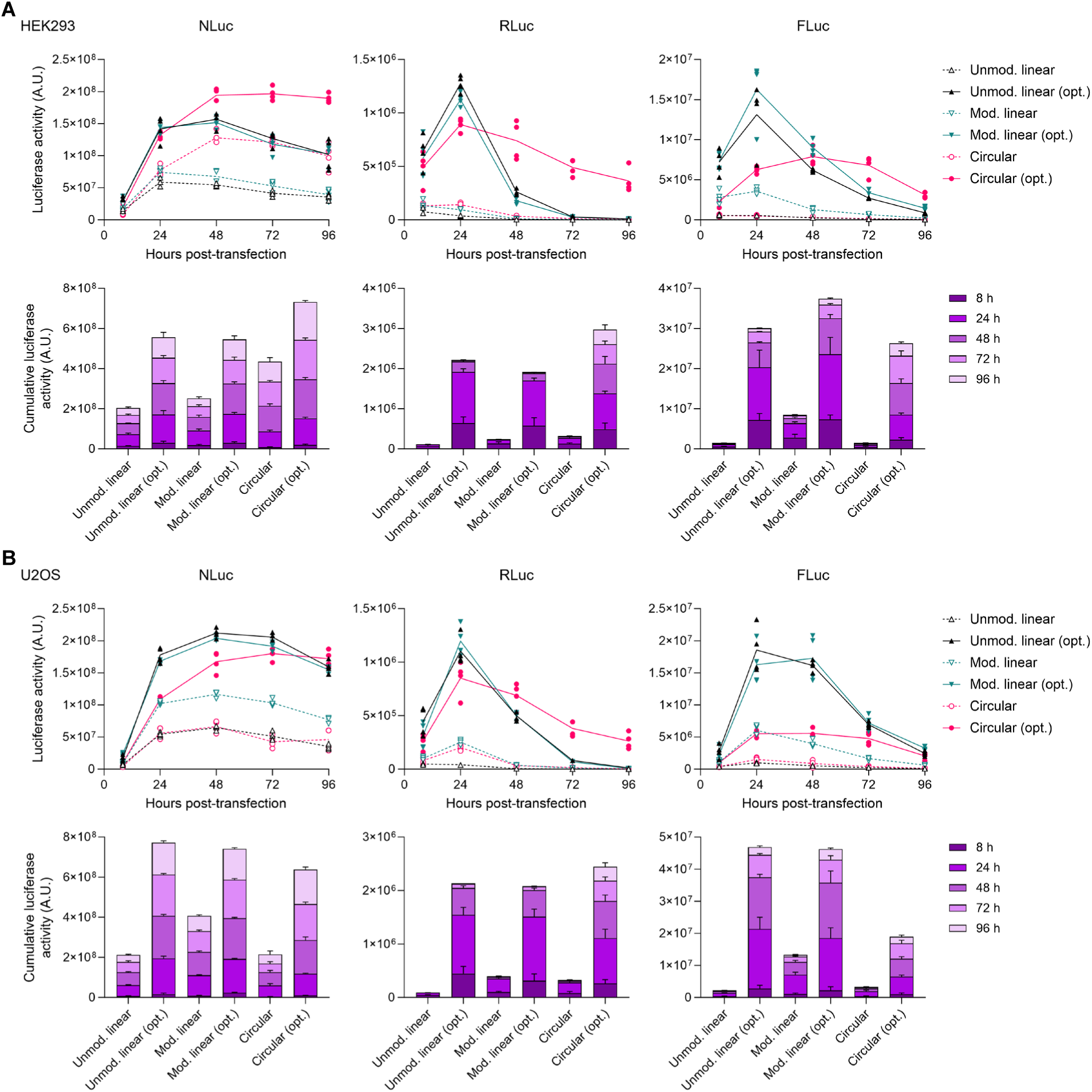
Optimal codons increase reporter protein expression from linear and circular RNAs. **(A)** HEK293 or **(B)** U2OS cells were transfected with unmodified linear, modified linear (100% m^1^Ψ substitution), or CVB3 IRES-driven circular mRNAs encoding either non-optimal or optimal forms of the indicated luciferase reporters. Cells were lysed and reporter protein expression was measured by the corresponding luciferase assay at 8, 24, 48, 72, and 96 h after transfection. The data are presented as a time-course of luciferase activity (top) and as accumulated luciferase activity from 8 h to 96 h (bottom). Error bars represent Std Dev of four independent replicates.

Since incorporating modified nucleotides into mRNAs and circularization have been shown to reduce the immunogenicity of exogenous RNAs, we also tested to see how modified mRNA and circRNAs affected the expression of the transfection control mRNA, eIF2α phosphorylation, and innate immunity genes. The co-transfection of different optimality reporters encoding RLuc with an EGFP control mRNA in HEK293 and U2OS cells revealed that the reduction of EGFP signals observed in non-optimal unmodified linear mRNA transfection was abrogated when modified linear mRNA or circRNA were used (Figure S5A). Next, we analyzed the level of eIF2α phosphorylation in HEK293 cells individually transfected with unmodified linear, modified linear, or circRNAs encoding non-optimal or optimal RLuc reporters. Western blot analysis shows that the induction of eIF2α phosphorylation by non-optimal reporters was diminished when the identical CDS was delivered as either modified linear mRNA or circRNA (Figure S5B). The immunogenicity of different RNA platforms encoding optimality reporters was then compared by RT-qPCR analysis of transfected U2OS cells. The expression of innate immunity genes induced by non-optimal sequences was significantly blunted when delivered by circRNAs or nucleoside-modified mRNAs (Figure S5C). While m^1^Ψ substitutions abolished immunogenicity of linear mRNAs, non-optimal circRNAs remained somewhat immunogenic, possibly linking GCN2 driven responses to RNA immunogenicity. Another possibility is that trace amounts of circRNA precursors or processing intermediates (that are below the detection threshold of FlashGels (Figure S1) drive the immune activation observed in our circRNA samples. Regardless, these data are consistent with each RNA therapy platform conferring unique immunostimulatory, or immune-evasive, properties.

### RNA circularization increases the lifespan of optimal reporter transcripts

circRNAs have been touted as a promising next-generation RNA therapy platform since they are resistant to cellular exonucleases and confer longer *in vivo* lifespan than mRNA [12]. Further, costly modified nucleotides and capping reagents are not required for circRNA production [13]. To de-couple mRNA lifespan from the half-life of their encoded reporter proteins, we fused CL1/PEST degron motifs to the C-terminus of each reporter protein to stimulate targeted degradation by proteasome [45, 46]. We estimated the half-life of all three luciferase reporter proteins in multiple cell lines by measuring luciferase activities of cells transfected with unmodified linear mRNA. Half the cells were treated with the translation inhibitor cycloheximide (CHX), and luciferase signals were measured 0, 1, 2, and 3 hours after CHX addition. Consistent with previous reports, luminescence data showed that the reporters bearing degron motifs had ∼8-fold lower signals than non-degron containing versions, and that the half-life of degron-bearing luciferase reporter proteins was ∼1 hour or less (Figures 5A and S7A) [45]. Notably, at the protein level, both non-optimal and optimal mRNA-encoded reporter proteins showed essentially identical half-lives, validating that the codon optimality of the RNA does not affect the stability of the encoded reporter protein. We then used an extended time-course to compare the luciferase activity of each destabilized reporter from unmodified linear, modified linear, and circRNAs. As predicted, circRNAs encoding optimal degron-tagged reporters have longer half-lives than their linear mRNA counterparts (Figures 5B, S6, and S7B). We also observed the differences in the half-lives of RNAs encoding different reporters that correlated with the length of the CDS, with RNAs encoding shorter ORFs being longer-lived. Notably, we did not assay RNA levels directly, but since the half-life of the degron-containing proteins are under an hour, we know that the RNA encoding them must be both present and actively translated to observe signals as many as four days later.

**Figure 5.**
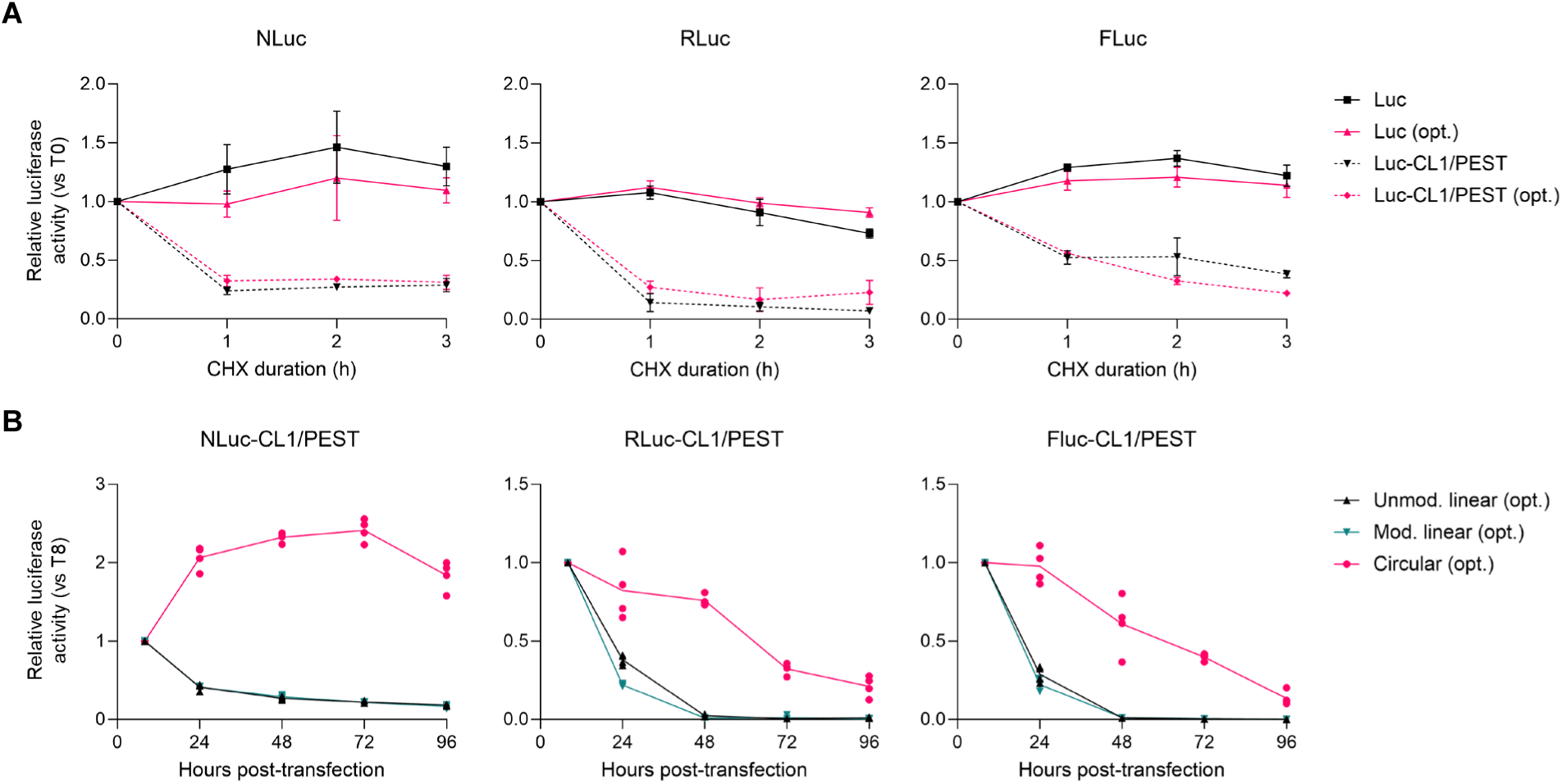
RNA circularization increases the lifespan of optimal reporter transcripts in HEK293 cells. Paired CL1/PEST degron motifs were fused to the indicated luciferase coding sequence to stimulate protein turnover. **(A)** HEK293 cells were co-transfected with linear mRNAs encoding a transfection control and the indicated reporters for 16 h. Luminescence signals were measured and normalized to the co-transfected control at the indicated time points after addition of cycloheximide (CHX, 100 μg/mL). The data shown for each reporter are shown as the mean ±Std Dev of four independent replicates and were compared to the time point when CHX was added (0 h). **(B)** HEK293 cells were co-transfected with a transfection control and either unmodified linear, modified linear, or circular mRNAs encoding optimal CL1/PEST degron-tagged luciferase reporters. Protein expression was measured by the corresponding luciferase assay at 8, 24, 48, 72, and 96 h after transfection. The data (shown as relative luciferase activity (RLU)) for each reporter are the mean of four independent replicates and were compared to the signal observed 8 h after transfection.

### Codon optimality of test mRNAs does not affect proteasome-mediated degradation

We next asked whether the codon optimality of the test RNAs influences global proteasome-mediated protein degradation which could conceivably cause the differences we observed in the expression of the transfection control reporter. We compared the degradation of degron-tagged reporters after treating two cell lines with the proteasome inhibitor MG-132 or DMSO as a control. Regardless of the reporter transfected, we did not observe significant differences in luciferase signals, which we use as a proxy for protein degradation rates, of the control reporter proteins after transfection with either non-optimal or optimal mRNAs (Figure S8).

Taken together, our data indicate that mRNAs enriched in non-optimal codons preferentially trigger cellular innate immune pathways and the phosphorylation of the translation initiation factor eIF2α, resulting in reduced global translation and at least a partial activation of cellular stress response. These responses were fully or partially abrogated by pre-treatment with A-92, a GCN2 inhibitor, or when nucleoside-modified or circular RNAs were used. Finally, we show that circularizing an optimal CDS enhances both RNA lifespan and the durability of protein expression from exogenous RNAs.

## Discussion

Despite the tremendous success of RNA therapeutics in various areas of medicine, their inherent instability and temporally limited translation capacity *in vivo* limit their application to many indications. Many previous reports have delineated the consequences of codon optimality on the translational regulation and stability of mRNAs encoded by different mixes of synonymous codons [20, 21, 36, 47, 48]. Indeed, our *in vivo* and *in vitro* data agree with those earlier observations. As shown by our small animal data, our optimal FLuc mRNA expressed markedly more protein for a longer duration than its non-optimal counterpart. Further, using three different reporters, our *in vitro* data demonstrate that optimal mRNAs also have superior expression parameters (peak luciferase activity and expression duration) for all three luciferase reporter proteins tested. This is in line with previous attempts to enhance protein expression from mRNAs by optimizing the codon optimality and/or global mRNA secondary structure [49, 50]. To further investigate how platform technology affects RNA persistence independent of protein stability, degron-tagged reporters with reduced half-life were used as a proxy for the presence of translatable RNA [45, 46] . While we acknowledge that measuring reporter protein activity is an imperfect and indirect measurement of RNA stability, our data show that circRNAs encoding optimal reporters exhibit superior persistence compared to their linear mRNA counterparts in an ORF-dependent manner. Differences in expression kinetics between linear and circular RNAs were observed, which is possibly due to distinct translation initiation mechanisms (cap-dependent vs. IRES) and the distinct topology of circRNAs. With our new optimization method, we provide further support to previous data demonstrating the importance of codon optimality for efficient translation. However, little has been published about the significance of codon optimality on cellular stress and innate immune responses with regards to exogenous RNAs. Elucidating any link(s) would be particularly timely since the use cases for RNA medicines will continue to expand, and such knowledge can drive the design of future therapies.

Here, we report that codon optimality influences both mRNA immunogenicity and cellular stress response in mammalian cells. We observed that non-optimal reporters markedly suppressed the activity of co-transfected control mRNA. In other words, the codon optimality of test mRNAs changed the protein output from the transfection control mRNAs. This is a key observation as the yields from the dual reporter assay’s transfection control must (and are assumed to) remain unchanged to accurately capture the changes observed in these reporter experiments. Importantly, since the readings derived from control RNAs do not remain constant in these experiments, our findings show that increased caution (and specific attention on both the normalized and raw signals from both the test and control RNAs) is needed when interpreting dual reporter experiments like those in this paper where the optimality of the test constructs is substantially different.

We show that non-optimal mRNAs induce eIF2α phosphorylation, a key mediator of the integrated stress response that globally suppresses translation initiation. In line with our observations in mammalian cells, experiments in yeast also show that consistent use of sub-optimal codons induces eIF2α phosphorylation in a CDS length-dependent manner [38]. We identified GCN2 kinase as the primary mediator of the integrated stress response, with minimal to no contribution from PKR or PERK. This implicates a translation-dependent trigger rather than dsRNA sensing or ER stress. While reporters enriched with rare codons might exhaust the charged tRNA pool, thereby triggering GCN2-sensed translational stress [39, 51, 52], this mechanism is unlikely to be the primary driver as identical sequences transcribed with m^1^Ψ – which would deplete the identical tRNAs – do not induce eIF2α phosphorylation.

Stalled and colliding ribosomes have also been shown to activate GCN2 kinase [39, 53, 54]. It is possible that slower translation elongation rates of non-optimal codons could lead to ribosome collisions, which have been linked to several cellular stress response pathways [24, 25, 38, 39, 51]. Furthermore, our data showing that GCN2 inhibition restores protein yields from non-optimal mRNAs supports a translation-dependent mechanism, likely with ribosome stalling or collisions serving as an optimality sensor. Although a previous work did not show a correlation between ribosome collision frequency and codon optimality [36], the discordance with our data likely stems from two key factors: our unique sequence design (Table S2) [36, 37], and our use of *in vitro* synthesized RNAs, contrasting with the plasmid and viral-based assays previously used [36].

Our results are consistent with previous reports showing that RNA circularization also tempers eIF2α phosphorylation [17, 55]. As IRES-driven translation initiation is less efficient than cap-dependent translation, we hypothesize that ribosome collisions on circRNA coding regions are limited since IRES-driven translation constrains ribosome loading [56]. Similar to RNA circularization, we show that m^1^Ψ modification also lowered eIF2α phosphorylation, indicating that platform design can temper stress signaling. Earlier work showed that substituting U with Ψ or m^1^Ψ increases amino acid misincorporation during translation in both prokaryotic and mammalian translation systems [57]. Although ribosome collisions were not assessed, a provocative hypothesis is that, on average, such m^1^Ψ-caused amino acid substitutions shorten ribosome residence time at sub-optimal codons, thus reducing pausing and collision frequency. However, further experiments are needed to determine whether the accumulation of uncharged tRNAs or ribosome stalling/collision initiate the stress response [39].

Notably, both CNOT3 and the remainder of the CCR4-NOT complex are known to mediate non-optimal RNA surveillance and turnover, likely explaining degradation of our linear mRNAs [22, 58, 59]. However, our data showing optimality-dependent changes in protein output from (and the half-lives of) our circRNA reporters is inconsistent with CNOT3-driven deadenylation since circRNAs lack poly(A) tails and thus suggest another mechanism contributes to surveilling non-optimal coding sequences.

Furthermore, we showed that codon optimality can also contribute to the immunogenicity of exogenous RNAs. Specifically, non-optimal mRNAs led to the induction of several ISGs, antiviral response genes, inflammatory cytokines, and innate immune regulators. Notably, substitutions of non-optimal codons by synonymous, optimal codons significantly mitigated innate immune response activation by unmodified linear mRNAs. ISG activation was also fully or partially abrogated by using m^1^Ψ and circularizing the RNAs containing identical coding sequences. This effect might be partly due to the lower uridine content in optimal mRNAs, which impedes recognition by TLRs [7, 9, 10, 60–62] and RIG-I activation [63, 64] and a reduction in double-stranded RNA by-products from m^1^Ψ-containing *in vitro* transcription reactions [65]. This observation also suggests that crosstalk between ribosome pausing (or stalling) caused by non-optimal codons and innate immunity gene activation is possible [26]. One possible explanation is that our linear mRNAs included trace amounts of dsRNA or other impurities that are sometimes produced by non-templated T7 polymerase activity in run-off *in vitro* transcription reactions [66–68]. While we cannot rule out residual impurities entirely, since the transcripts – optimal and non-optimal– all begin and end in identical sequences and are synthesized by identical methods, any residual contaminants would be present in equal concentrations across all samples. Indeed, as judged by eIF2α phosphorylation, our experiments with C16, a PKR inhibitor, show that dsRNA is only a minor (and optimality-independent) contributor to the stress response observed in response to exogenous mRNAs.

While increasing protein output is often a key goal of most synthetic protein coding sequences, we must also note that achieving maximal codon optimality will not always be desirable for every synthetic RNA. Indeed, doing so could be counterproductive as maximizing translation rates could have negative effects [69–72] with the use of less optimal codons allowing for proper nascent polypeptide chain folding [73–77]. The inclusion of non-optimal codons to slow translation rates may be particularly important for the folding of intrinsically disordered regions [72, 78]. As an example, increasing the optimality of the mRNA encoding the CFTR gene has been reported to impair the protein’s ability to fold properly [72, 79, 80]. Similar dynamics have been demonstrated for other proteins associated with human diseases such as KRAS in cancers and factor IX blood coagulation disorders [81, 82].

In conclusion, the codon optimality and the platform technology of the exogenous RNAs both play important roles in evading (or harnessing) cellular stress and innate immune responses. Importantly, mRNAs enriched in poorly optimal codons can trigger cellular stress and alter the expression of unintended mRNAs. Further, since each RNA platform technology possesses intrinsic strengths and weaknesses, sequence engineering approaches can be combined with platform selection to leverage their capabilities to best suit particular biomedical applications. In particular, modified linear mRNAs will likely remain the preferred platform technology to express proteins where a short-term burst of expression lasting 1-3 days is needed. Our data agrees with those from others that show circRNAs could be the best platform technology for extended protein expression but suggest that the optimality of the coding region is a key determinant of circRNA lifespan *in vivo*.

## Materials and Methods

All materials, reagents, their suppliers, and catalog numbers are included in Table S3: Key Resources.

### Plasmid construction

The template plasmids used for *in vitro* transcription of linear and circular mRNAs encoding non-optimal luciferase reporters were constructed using e-Zyvec’s (Polyplus-Sartorius) whole plasmid synthesis service. The design of the degron-containing luciferase reporters was patterned after those reported here [45]. The linear mRNA template consists of a T7 promoter, 5’ UTR, NcoI and XhoI cloning sites for inserting the ORFs, 3’ UTR, 90-nt poly(A) tail, followed by BspQI restriction site. The circRNA template begins with a T7 promoter and ends with a BspQI restriction site. Between the T7 promoter and the restriction site, homology regions, spacers and permuted *Anabaena* pre-tRNA group I intron-exon sequences on the 5’ and 3’ ends flank a CVB3 IRES and the indicated coding region that was adapted from the literature [12, 13]. The optimal reporter gene fragments were synthesized by Twist Bioscience and subcloned into linear mRNA and circRNA vectors by ligation and HiFi DNA assembly, respectively. To generate optimal linear mRNA templates, the linear mRNA backbone and gene fragments were digested by NcoI and XhoI followed by ligation. The circRNA backbone was PCR amplified using primers CS31 and CS32 (see Table S4 for all primer sequences). Gene fragments were PCR amplified using primers CS33 and CS34. The amplified products were gel purified and assembled by NEBuilder HiFi DNA Assembly master mix. All linear mRNA and circRNA vectors were transformed into NEB stable competent *E. coli* and NEB 5-alpha competent *E. coli*, respectively. The plasmids were miniprepped with Monarch Plasmid Miniprep Kit and sequence-verified with whole-plasmid sequencing. For linear mRNA templates, the length of poly(A) tail was verified by Sanger sequencing. All novel plasmids will be provided in a timely manner upon request.

### *In silico* RNA Sequence assessment

The minimum free energy (MFE) was calculated from the predicted RNA secondary structure using RNAFold v2.7.0 at the default settings[83]. The codon adaptation index (CAI) was calculated using CAI calculator on https://www.biologicscorp.com/tools/CAICalculator/ [29].

### mRNA synthesis and purification

The sequences for all mRNAs and circRNAs used in the study, and also the sequence of a representative circRNA precursor prior to circularization are shown in Table S5. Unmodified or modified linear mRNAs were synthesized by *in vitro* transcription from a linearized plasmid DNA template and co-transcriptionally capped using a HiScribe^®^ T7 mRNA Kit with CleanCap Reagent AG Kit according to the manufacturer’s instructions. For modified linear mRNAs, UTP was replaced by *N*^1^-methylpseudo-UTP. circRNA precursors were synthesized by *in vitro* transcription from a linearized plasmid DNA template using a HiScribe T7^®^ High Yield RNA Synthesis Kit according to the manufacturer’s instructions. The *in vitro* transcription reactions were treated with DNase I to remove residual DNA template and RNAs were purified by Monarch Spin RNA Cleanup Kit. We modified an earlier method to make circRNA [12, 84] . Briefly, GTP was added to a final concentration of 2 mM to the purified circRNA precursor (see Table S5) in circularization buffer containing 50 mM Tris-HCl, 10 mM MgCl_2_, 1 mM DTT, pH 7.5. The RNA was then heated at 55°C for 8 min. The circRNA products were further purified using a Monarch Spin RNA Cleanup Kit. To remove leftover linear RNA species, the purified circRNA products were treated with 0.5 U of RNase R per 1 μg of circularized RNA at 37°C for 1 h. The quality and quantity of RNA were evaluated by Denovix DS-11 FX+ Spectrophotometer and FlashGel RNA Cassettes (Figure S1B-D).

### Encapsulation of RNA into lipid nanoparticles (LNP)

LNP containing optimal and non-optimal FLuc RNA were prepared as previously described [85]. Briefly, microfluidic mixing by NanoAssemblr BT Microfluidics Mixing System (Precision NanoSystems, Vancouver, CA) of an ethanolic phase containing 0.8 mMol 1,2-distearoyl-sn-glycero-3-phosphocholine (DPSC), 3.85 mMol cholesterol, 0.15 mMol 1,2-distearoyl-sn-glycero-3-phosphoethanolamine-N-[methoxy(polyethylene glycol)-2000] (ammonium salt) (DSPE-P E G ( 200 0) ) , and 5. 2 m M ol he pt ade c a n - 9 - y l 8 - ((2 - hy dr ox y et hy l ) ( 6 - ox o - 6 - (undecyloxy)hexyl)amino)octanoate (SM-102) with an aqueous phase containing 150 μg RNA (optimal or non-optimal) in 0.1 M Citrate Buffer (pH 5) at 1:3 ratio at a flow rate of 10 mL/min was performed. To remove residual ethanol and unencapsulated RNA, LNP were dialyzed using D-Tube Dialyzer Maxi, 12-14 kDa with 1X PBS Buffer. LNP were then characterized for hydrodynamic diameter, polydispersity index and zeta potential by dynamic light scattering (Malvern Zetasizer, Malvern Instruments). RNA content was assessed by RiboGreen RNA Assay as described by the manufacturer. The LNP had a diameter in the range of 70-110 nm, neutral zeta potential and encapsulation efficiency of >90%. Final working concentrations were diluted in 1X PBS before animal injections.

### *In Vivo* assessment of FLuc RNA expression

All animal experiments were conducted in accordance with protocols approved by the institutional animal care and use committee (IACUC) at the Houston Methodist Research Institute. Female BALB/c mice (4-5-week-old) were purchased from Jackson Laboratories. The mice were housed with controlled temperature (25 °C), 12:12h lighting cycle, and access to standard diet and water.

Mice received flank injections with 5 μg (in 100 μL PBS) of RNA encapsulated in LNP. To reduce intra-animal variability, each mouse (n=5) received an intramuscular injection with LNP containing optimal FLuc RNA LNP (right flank) and non-optimal FLuc RNA LNP (left flank). Mice were imaged daily for 3 days to assess bioluminescent signal following intraperitoneal injection of 150mg/kg luciferin using IVIS System (Perkin Elmer). IVIS bioluminescence imaging was performed for whole animal (dorsal view), left flank, and right flank positions. After imaging at 72h (3 days), the animals were sacrificed according to IACUC-approved protocols.

### Cell culture and transfection

HEK293 and NIH/3T3 cells were cultured in Dulbecco’s modified Eagle’s medium (DMEM) plus 10% FBS; U2OS cells in McCoy’s 5A plus 10% FBS; BJ cells in Eagle’s Minimum Essential Medium (EMEM) plus 10% FBS. All cells were maintained in a 37°C incubator with 5% CO_2_ and passaged approximately every 3 days. mRNAs were transfected with Lipofectamine MessengerMAX Transfection Reagent at equimolar concentrations unless stated otherwise along with an internal control mRNA per the manufacturer’s protocols. The total amount of RNA transfected in each experiment ranged between 20 and 70 ng/well of a 96 well tissue culture dish. The transfection reagent was removed, and fresh medium was added to the cells 6 h after transfection.

### *In vitro* Luciferase assays

The reagents for the dual luciferase assays were obtained from Promega and the assays were performed according to the manufacturer’s instructions. Briefly, cells were lysed with Passive Lysis Buffer at the indicated time points after transfection. For Firefly/*Renilla* assays, luciferase activity was measured using Dual-Glo Luciferase Assay System. For Firefly/NanoLuc assays, luciferase activity was measured using Nano-Glo Dual-Luciferase Reporter Assay System. Luminescence was measured on a white 96-well flat-bottom microplate using a SpectraMax iD5 Microplate Reader (Molecular Devices).

For protein half-life determination, 24 h after transfection, cells were either lysed with the passive lysis buffer and frozen at -20°C (for T0 control) or treated with 100 μg/mL cycloheximide (CHX) for the indicated amounts of time before lysis/freezing. Luciferase assays were performed on thawed samples using the appropriate dual-reporter assay system mentioned above.

For determining proteasome-mediated protein degradation, cells were transfected with degron-tagged reporter mRNAs for 16 h. Subsequently, cells were transfected with the optimality reporter mRNAs and treated with DMSO (for control) or 10 μg/mL MG-132. After 6 h, cells were lysed with passive lysis buffer and luciferase assays were performed using the corresponding dual-reporter assay system.

### Live fluorescence microscopy imaging

An Incucyte S3 Live-Cell Analysis System (Sartorius) was used to measure and quantify the levels of EGFP control mRNA expression. Four independently transfected wells were used to measure each time point and condition. For each well, five images from different positions within each well were acquired at time points indicated after transfection for each signal determination. The signals for each image were quantified using the Incucyte S3 software (Incucyte 2022B Rev2), averaged for each well, and are shown as the mean fluorescence intensity.

### Cell viability assay

HEK293 cells were transfected with unmodified non-optimal or optimal RLuc reporter mRNAs. After 24 h and 48 h, cell viability was assayed using CellTiter-Glo Luminescence Cell Viability Assay according to the manufacturer’s protocol. Briefly, a volume of the reagent equal to the cell culture medium was added to each well. After 10-minute incubation at room temperature, luminescence was measured on a white 96-well flat-bottom microplate using a SpectraMax iD5 Microplate Reader (Molecular Devices).

## Western blot analysis

Sixteen hours after transfection, media was removed, and cells were washed with phosphate-buffered saline (PBS). Cells were collected by scraping with a cell lifter and lysates prepared using NP-40 lysis buffer (50 mM Tris, pH 8.0, 150 mM NaCl, 1% NP-40) with protease and phosphatase inhibitors. Proteins were collected from the supernatant after centrifugation at 16,000 x g for 20 min. Protein concentrations were determined using the Pierce BCA protein assay kit using a bovine serum albumin standard curve. Protein samples were then prepared in Laemmli sample buffer containing 50 mM DTT and heated at 95°C for 5 min. Samples were run on a 12% or 4-20% Mini-PROTEAN TGX Precast Protein Gel with Tris/Glycine/SDS buffer. The protein was transferred to a 0.2 μM PVDF membrane using Trans-blot Turbo Transfer System according to the manufacturer’s instructions. The membrane was blocked with 1x TBST with 5% BSA and sustained rocking at room temperature for 1 h. The following antibodies were used: phosphorylated eIF2α (Ser51) (1:1000 dilution), eIF2α (1:1000 dilution), α-tubulin (1:20000 dilution), β-actin HRP-conjugated (1:5000 dilution), FLAG Tag (1:5000 dilution),HRP-conjugated goat anti-rabbit IgG (1:5000 dilution), HRP-conjugated horse anti-mouse IgG (1:5000 dilution). Primary antibodies were diluted in 1x TBST with 5% BSA, and blots were incubated overnight at 4°C with sustained rocking, except for HRP-conjugated β-actin, which was incubated for 1 h at room temperature. Membranes were washed three times with 1x TBST and incubated with secondary antibodies with rocking at room temperature for 1 h.

For lysates of cells treated with kinase inhibitors, membranes were blocked in Everyblot (Bio-Rad) for 20 min with rocking at room temperature. Primary and secondary antibodies were subsequently diluted in Everyblot blocking buffer as well. Membranes were washed three times with 1x TBST, and protein signal was detected with the Clarity Western ECL Substrate and imaged on the ChemiDoc MP Imaging System (Bio-Rad).

For eIF2α kinase inhibition experiments, cells were pre-treated prior to mRNA transfection with individual kinase inhibitors at concentrations and durations that were demonstrated effective in the literature [42, 43]. For PERK inhibition, cells were pre-treated with AMG-44 at 20 μM for 2.5 h [43]. For GCN2 inhibition, cells were pre-treated with A-92 at 1 μM for 5 h[42]. For PKR inhibition, cells were pre-treated with C16 at 1 μM for 1 h [42].

### RT-qPCR

Total RNA was extracted from cultured cells by adding TRI Reagent to the cells and RNA was extracted by phase separation and isopropanol precipitation per the manufacturer’s instructions. One μg of total cellular RNA was treated with DNase I and reverse transcribed with random hexamer primers and ProtoScript II Reverse Transcriptase. RT-qPCR was performed with SSOAdvanced Universal SYBR Green Supermix on a CFX96 Touch Real-Time PCR Detection System. Each sample was measured in technical triplicates, and three biological replicates were performed per experiment. All primers used are listed in Table S4. Relative gene expression was calculated by the 2^-ΔΔCt^ method using β-actin as an internal control.

## Data Availability

All Supplementary Data are available Online. The underlying replicate data for the experiments performed in this study will be made available upon request.

## CRediT authorship contribution statement

CS: Conceptualization, Methodology, Investigation, Formal Analysis, Writing – original draft, Writing – review & editing, Data curation, Visualization, Supervision; NB: Methodology, Investigation, Formal Analysis, Writing – review & editing, Data curation, Visualization; EC: Methodology, Investigation, Formal Analysis, Writing – review & editing; TN: Investigation, Data curation, Formal Analysis; BG: Methodology, Formal Analysis, Writing – review & editing, Data curation, Visualization, Resources, Funding Acquisition, Supervision; DLK: Conceptualization, Methodology, Investigation, Formal Analysis, Writing – original draft, Writing – review & editing, Visualization, Resources, Funding Acquisition, Supervision.

## Acknowledgements

We would like to thank all the other members of the Kiss RNA Lab for their support and fruitful discussions regarding the project. We would like to thank the staff of the Advanced Cellular and Tissue Microscopy Core Facility at the Houston Methodist Research Institute (HMRI) for technical support. BG, EC and TN acknowledge the support of CPRIT RP200619 and CDMRP MB230026 (to BG). The codon optimality portion of this work was supported by a grant from the National Institute of General Medical Sciences [R35GM137819] and circRNA production was supported by a Career Cornerstone Award and an RNA therapeutics fund from Houston Methodist Hospital Foundation (to DLK).

The content presented here is solely the responsibility of the authors and does not represent the official views of the Cancer Prevention and Research Institute of Texas (CPRIT), the Department of Defense, the HMRI, or the National Institutes of Health.

