## Supplemental Tables for "Codon optimality modulates cellular stress and innate immune responses triggered by exogenous RNAs"

**Supplementary Tables**

**Table S1: Sampling of Human Gene Codon Adaptation Index (CAI) scores**
Genes marked with an * are known to have mutations linked to protein folding. Genes marked with ** have been linked to co-translational protein folding errors.

| Gene Symbol | Refseq ID | # of Amino Acids | CAI Score |
|---|---|---|---|
| ACTB | NM_001101.5 | 375 | 0.90 |
| GAPDH | NM_002046.7 | 335 | 0.83 |
| HBB | NM_000518.5 | 147 | 0.87 |
| TUBB | NM_178014.4 | 444 | 0.83 |
| NUP62 | NM_016553.5 | 522 | 0.83 |
| PDGFRA | NM_006206.6 | 1089 | 0.77 |
| PDGFRB | NM_002609.4 | 1106 | 0.82 |
| TTN | NM_001267550.2 | 35991 | 0.73 |
| DMD** | NM_004006.3 | 3685 | 0.72 |
| CFTR** | NM_000492.4 | 1480 | 0.70 |
| KRAS** | NM_004985.5 | 188 | 0.70 |
| COL1A1* | NM_000088.4 | 1464 | 0.77 |
| COL1A2* | NM_000089.4 | 1366 | 0.72 |
| COL3A1* | NM_000090.4 | 1466 | 0.73 |
| TARDBP | NM_007375.4 | 414 | 0.75 |
| SOD1** | NM_000454.5 | 154 | 0.78 |
| CLN3 | NM_001042432.2 | 438 | 0.80 |
| APP | NM_000484.4 | 770 | 0.79 |
| T53* | NM_000546.6 | 393 | 0.81 |
| EEF2 | NM_001961.4 | 858 | 0.87 |
| INS | NM_000207.3 | 110 | 0.85 |
| SERPINI1* | NM_001122752.2 | 410 | 0.73 |
| SERPINA1* | NM_000295.5 | 418 | 0.83 |
| VCL | NM_014000.3 | 1134 | 0.76 |
| FHIT | NM_002012.4 | 147 | 0.79 |
| RNGTT | NM_003800.5 | 597 | 0.74 |
| RNMT | NM_003799.3 | 476 | 0.73 |
| RPL37 | NM_000997.5 | 97 | 0.74 |
| RPLP0 | NM_001002.4 | 317 | 0.81 |
| RPL27 | NM_000988.5 | 136 | 0.84 |
| RPL31 | NM_000993.5 | 125 | 0.78 |
| RB1* | NM_000321.3 | 928 | 0.70 |
| MYH7* | NM_000257.4 | 1935 | 0.87 |
| LAMA2* | NM_000426.4 | 3122 | 0.75 |
| GBA1 | NM_000157.4 | 536 | 0.79 |
| HK1 | NM_000188.3 | 917 | 0.81 |
| ATM | NM_000051.4 | 3056 | 0.68 |
| FMR1 | NM_002024.6 | 632 | 0.73 |
| BRCA1 | NM_007294.4 | 1863 | 0.72 |
| BRCA2 | NM_000059.4 | 3418 | 0.70 |
| COMT* | NM_000754.4 | 271 | 0.86 |
| NKX2-5 | NM_004387.4 | 324 | 0.78 |

| Gene Symbol | Refseq ID | # of Amino Acids | CAI Score |
|---|---|---|---|
| VCP | NM_007126.5 | 806 | 0.78 |
| SIRT1 | NM_012238.5 | 747 | 0.70 |
| SIRT6 | NM_016539.4 | 328 | 0.84 |
| PTEN | NM_000314.8 | 403 | 0.73 |
| TERT | NM_198253.3 | 1132 | 0.81 |
| FBN1* | NM_000138.5 | 2871 | 0.80 |
| PTBP3 | NM_001163788.4 | 524 | 0.70 |
| RHO | NM_000539.3 | 348 | 0.85 |
| PMP22** | NM_000304.4 | 160 | 0.78 |
| SNCA | NM_000345.4 | 140 | 0.77 |
| HTT* | NM_001388492.1 | 3142 | 0.75 |
| MECP2 | NM_004992.4 | 486 | 0.83 |
| LDLR* | NM_000527.5 | 860 | 0.84 |
| HEXA* | NM_000520.6 | 529 | 0.78 |
| C9orf72 | NM_018325.5 | 481 | 0.70 |
| MAPT* | NM_001377265.1 | 833 | 0.79 |
| PAH* | NM_000277.3 | 452 | 0.77 |
| RET* | NM_020975.6 | 1114 | 0.81 |
| OPA1 | NM_130837.3 | 1015 | 0.69 |
| THRA | NM_199334.5 | 410 | 0.85 |
| THRB | NM_001354712.2 | 461 | 0.79 |
| NF1 | NM_001042492.3 | 2839 | 0.70 |
| EGFR | NM_005228.5 | 1210 | 0.80 |
| VHL | NM_000551.4 | 213 | 0.78 |
| MLH1 | NM_000249.4 | 756 | 0.76 |
| LMNA* | NM_170707.4 | 664 | 0.84 |
| DIS3L | NM_001143688.3 | 1054 | 0.75 |
| HES5 | NM_001010926.4 | 166 | 0.86 |
| S100A16 | NM_080388.3 | 103 | 0.86 |
| CTNNB1* | NM_001904.4 | 781 | 0.70 |
| DDX5 | NM_004396.5 | 614 | 0.73 |
| EML4 | NM_019063.5 | 981 | 0.72 |
| GPX4 | NM_002085.5 | 197 | 0.85 |
| PFKM | NM_000289.6 | 780 | 0.79 |
| HSPA1A | NM_005345.6 | 641 | 0.88 |
| HSP90AA1 | NM_005348.4 | 732 | 0.75 |
| DNAJB1 | NM_006145.3 | 340 | 0.81 |
| HSPB1 | NM_001540.5 | 205 | 0.82 |
| PDIA2 | NM_006849.4 | 525 | 0.81 |
| ATF4 | NM_182810.3 | 351 | 0.78 |
| VIM | NM_003380.5 | 466 | 0.81 |
| MYH2 | NM_017534.6 | 1941 | 0.81 |
| SCAMP1 | NM_004866.6 | 338 | 0.70 |
| TXN | NM_003329.4 | 105 | 0.75 |
| AKT1 | NM_001382430.1 | 480 | 0.87 |
| MTOR | NM_004958.4 | 2549 | 0.79 |
| BUB1 | NM_004336.5 | 1085 | 0.73 |
| CDC42 | NM_001791.4 | 191 | 0.71 |

**Table S2: Codon selection differences between three codon optimality formulas**

The Codon optimality formula employed in this paper is compared to two other sequence designs for Firefly Luciferase protein. Only the most favored codon for each amino acid encoded by multiple tRNAs is shown. Differences in codon selection frequency are shown both as a % change and as a count of how many codons are different between the most optimal design listed in Barrington et al. or as the highest optimality output provided by the iCodon webtool [36, 37].

| Most used Codon (per AA) | Codon use (this paper) | Usage in Barrington et al. | % change | # different v Barrington (of 550) | Usage in iCodon | % change | # different vs iCodon (of 550) |
|---|---|---|---|---|---|---|---|
| Leu/CTG | 96% | 44% | **+52%** | 27 | 45% | **-51%** | 26 |
| Cys/TGC | 100% | 100% | 0% | -- | 50% | **-50%** | 2 |
| Tyr/TAC | 100% | 100% | 0% | -- | 63% | **-37%** | 7 |
| Phe/TTC | 97% | 100% | **-3%** | 1 | 50% | **-47%** | 14 |
| Asp/GAC | 94% | 100% | **-6%** | 2 | 34% | **-60%** | 19 |
| Glu/GAG | 94% | 100% | **-6%** | 2 | 30% | **-64%** | 21 |
| Gln/CAG | 94% | 100% | **-6%** | 1 | 25% | **-69%** | 11 |
| Asn/AAC | 88% | 100% | **-12%** | 2 | 53% | **-35%** | 6 |
| His/CAC | 86% | 100% | **-14%** | 2 | 43% | **-43%** | 6 |
| Ala/GCC | 79% | 44% | **-35%** | 15 | 30% | **-49%** | 21 |
| Ile/ATC | 92% | 49% | **-43%** | 16 | 59% | **-33%** | 13 |
| Val/GTG | 93% | 44% | **-49%** | 22 | 29% | **-64%** | 29 |
| Pro/CCC | 66% | 0% | **-66%** | 19 | 24% | **-42%** | 12 |
| Gly/GGC | 98% | 43% | **-55%** | 25 | 17% | **-81%** | 26 |
| Ser/AGC | 86% | 39% | **-47%** | 13 | 4% | **-82%** | 23 |
| Arg/CGG | 80% | 30% | **-50%** | 10 | 5% | **-75%** | 15 |
| Lys/AAG | 97% | 0% | **-97%** | 38 | 51% | **-46%** | 39 |
| Thr/ACC | 83% | 0% | **-83%** | 24 | 38% | **-45%** | 13 |
| # Different AAs (only 1st choice codons) | | | | 219 | | | 303 |

**Table S3: Key Resources**

| Reagents or resources | Source | Identifier (Cat. No.) |
|---|---|---|
| **Antibodies** | | |
| α-phosphorylated eIF2α (Ser51) | Cell Signaling Technology | 3398S |
| α-eIF2α | Cell Signaling Technology | 5324S |
| α-alpha tubulin | Proteintech | 66031-1-Ig |
| HRP-conjugated goat anti-rabbit IgG | Cell Signaling Technology | 7074S |
| HRP-conjugated horse anti-mouse IgG | Cell Signaling Technology | 7076S |
| α-beta actin (13E5) rabbit mAb (HRP Conjugate) | Cell Signaling Technology | 5125S |
| α-FLAG-tag | Proteintech | 66008-3-Ig |
| **Bacterial strains** | | |
| NEB 5-alpha competent *E. coli* | NEB | C2987U |
| NEB stable competent *E. coli* | NEB | C3040H |
| **Reagents & materials** | | |
| NEBuilder HiFi DNA Assembly master mix | NEB | E2621S |
| Monarch Plasmid Miniprep Kit | NEB | T1010L |
| HiScribe® T7 mRNA Kit with CleanCap Reagent AG Kit | NEB | E2080S |
| HiScribe T7 High Yield RNA Synthesis Kit | NEB | E2040S |
| $N^1$-methylpseudo-UTP | ThermoFisher Scientific | NU505201 |
| DNase I | NEB | M0303S |
| Monarch Spin RNA Cleanup Kit | NEB | T2040L |
| RNase R | Lucigen Corporation | RNR07250 |
| FlashGel RNA Cassettes | Lonza | 57027 |
| 100X Pen/Strep | Gibco | 15140-122 |
| DMEM, high glucose | Gibco | 11965092 |
| FBS | Corning | 35-075-CV |
| McCoy's 5A | Gibco | 16600-082 |
| Phosphate Buffered Saline (PBS) | GenClone | 25-507B |

| | | |
|---|---|---|
| OptiMEM I (1X) Reduced Serum Media | Gibco | 11058-021 |
| Trypsin-EDTA 0.25% | Gibco | 25200-056 |
| DMSO | Sigma | D4540 |
| Lipofectamine MessengerMAX Transfection Reagent | ThermoFisher Scientific | LMRNA015 |
| Passive Lysis Buffer | Promega | E1941 |
| Dual-Glo Luciferase Assay System | Promega | E2920 |
| Nano-Glo Dual-Luciferase Reporter Assay System | Promega | N1620 |
| Cycloheximide (CHX) | Sigma | C1988 |
| MG-132 | Sigma | 474787 |
| Pierce BCA protein assay kit | ThermoFisher Scientific | 23225 |
| Laemmli sample buffer | Bio-Rad | 1610747 |
| 12% Mini-PROTEAN TGX Precast Protein Gel | Bio-Rad | 4561043 |
| 4-20% Mini-PROTEAN TGX Precast Protein Gel | Bio-Rad | 4561094 |
| 10X Tris/Glycine/SDS buffer | Bio-Rad | 1610732 |
| Trans-blot Turbo Transfer System | Bio-Rad | 1704272 |
| Trans-Blot Turbo 5x Transfer Buffer | Bio-Rad | 10026938 |
| EveryBlot Blocking Buffer | Bio-Rad | 12010020 |
| 10X TBST | APEX | 18-235B |
| BSA | ThermoFisher Scientific | BP9706-100 |
| Halt Protease Inhibitor Cocktail (100X) | ThermoFisher Scientific | 78430 |
| Phosphatase Inhibitor Cocktail II | Sigma | P5726 |
| Phosphatase Inhibitor Cocktail III | Sigma | P0044 |
| PMSF 0.1M | Sigma | 93482-50ml-F |
| NP-40 (10%) | ThermoFisher Scientific | 28324 |
| Precision Plus Protein Dual Color Standards | Bio-Rad | 1610374 |
| Clarity Western ECL Substrate | Bio-Rad | 1705061 |
| TRI Reagent | Zymo Research Corporation | R2050-1-200 |
| Chloroform | Sigma | C2432 |
| Random hexamer primers | Promega | C1181 |
| ProtoScript II Reverse Transcriptase | NEB | M0368S |

| | | |
|---|---|---|
| SSOAdvanced Universal SYBR Green Supermix | Bio-Rad | 1725274 |
| Cell Titer-Glo Luminescence Cell Viability Assay | Promega | G7570 |
| Trans-Blot Turbo Mini PVDF Membrane, 0.2 µm | Bio-Rad | 12023954 |
| AMG-44 | Sigma | SML3049-5MG |
| A-92 | Axon Medchem | Axon 2720 |
| C16 | Sigma | I9785-5MG |
| dNTP Solution mix | NEB | N0447S |
| **Cell lines** | | |
| BJ | ATCC | CRL-2522 |
| Flp-In T-Rex HEK293 | Gibco | R78007 |
| NIH/3T3 | ATCC | CRL-1658 |
| U2OS | ATCC | HTB-96 |
| ***In Vivo* studies** | | |
| Female BALB/c mice | The Jackson Laboratories | 000651 |
| D-Luciferin | Gold Biotechnology | LUCK |
| 1,2-Distearoyl-sn-glycero-3-phosphocholine (DPSC) | Avanti Polar Lipids | 850365P |
| Cholesterol | Sigma | C8667 |
| 1,2-Distearoyl-sn-glycero-3-phosphoethanolamine-N-[methoxy(polyethylene glycol)-2000] (ammonium salt) (DSPE-PEG(2000)) | Avanti Polar Lipids | 880120P |
| Heptadecan-9-yl 8-((2-hydroxyethyl)(6-oxo-6-(undecyloxy)hexyl)amino)octanoate (SM-102) | Avanti Polar Lipids | 792885 |
| Citrate buffer (pH 5) | ThermoFisher | J62918.AP |
| D-Tube Dialyzer Maxi, 12-14 kDa | Sigma | 71510-3 |
| RiboGreen RNA Assay | ThermoFisher | R11490 |
| **Software and web server** | | |
| GraphPad Prism 10.1.2 | | |
| ImageLab v.4.1 | | |

| |
|---|
| Incucyte 2022B Rev2 |
| RNAfold 2.7.0 |
| https://www.biologicscorp.com/tools/CAICalculator/ |

**Table S4: Oligonucleotide primers used in this study**

| Primer name | Sequence | Purpose |
|---|---|---|
| CS31 | TATGCGTTACCGGCGAGACGCTAC | PCR cloning |
| CS32 | CCTCTTTCAAGCTAAGTGGTATAAACCC | PCR cloning |
| CS33 | GGGTTTATACCACTTAGCTTGAAAGAGG | PCR cloning |
| CS34 | GTAGCGTCTCGCCGGTAACGCATA | PCR cloning |
| IFI44 Forward | TGGGAGCTGGACCCTGTAAA | qPCR |
| IFI44 Reverse | CCTCCCTTAGATTCCCTATTTGCT | qPCR |
| DDX60 Forward | CCGAGGAAGGAAAATGTCGC | qPCR |
| DDX60 Reverse | CTCACGCAAGGAAACACTGATA | qPCR |
| IFIT2 Forward | GCACTGCAACCATGAGTGAGA | qPCR |
| IFIT2 Reverse | GCCTCGTTTTGCCCTTTGAG | qPCR |
| ISG15 Forward | CCAGGATGCTCAGAGGTTCG | qPCR |
| ISG15 Reverse | GGGACCTGACGGTGAAGATG | qPCR |
| RSAD2 Forward | CTCTGTGGAGGAGCCTGGTC | qPCR |
| RSAD2 Reverse | AAGTTGATCTTCTCCATACCAGCTT | qPCR |
| TNF-α Forward | TCCCCAGGGACCTCTCTCTA | qPCR |
| TNF-α Reverse | AGGGTTTGCTACAACATGGGC | qPCR |
| IL-6 Forward | AGCCACTCACCTCTTCAGAAC | qPCR |
| IL-6 Reverse | GCCTCTTTGCTGCTTTCACAC | qPCR |
| RIG-I Forward | TGTGGGCAA TGTCA TCAAAA | qPCR |
| RIG-I Reverse | GAAGCACTTGCTACCTCTTGC | qPCR |
| MDA5 Forward | GGCACCATGGGAAGTGATT | qPCR |
| MDA5 Reverse | ATTTGGTAAGGCCTGAGCTG | qPCR |
| OAS1 Forward | GCTCCTACCCTGTGTGTGTGT | qPCR |
| OAS1 Reverse | TGGTGAGAGTACTGAGGAAGA | qPCR |
| OASL Forward | AGGGTACAGATGGGACATCG | qPCR |
| OASL Reverse | AAGGGTTCACGATGAGGTTG | qPCR |
| PKR Forward | TCTTCATGTATGTGACACTGC | qPCR |
| PKR Reverse | CACACAGTCAAGGTCCTT | qPCR |
| ACTB Forward | CCCGCGAGCACAGAGCCTCGCCTTTGCCGA | qPCR |
| ACTB Reverse | CCTTCTGACCCATGCCCACCATCACGCCC | qPCR |

**Table S5: RNA sequences used in this study (stand-alone table as excel file)**

The RNA sequences for the different reporter RNAs are shown. Please note that all linear mRNAs begin with an 'A' as the first transcribed nucleotide and were in vitro transcribed using NEB's HiScribe T7 + Clean Cap AG kit. Therefore, mRNAs have an m7G cap1 structure. For all RNAs, coding sequences are in **bold** and 3xFLAG and 3xFLAG+hCL1+hPEST degron sequences are in **purple**. Please note that the RNA sequences for FLAG+/- degron tags are different for optimal/non-optimal RNAs. IRES sequences are underlined, UTR sequences are in *italics*, Stop codons are in **red**, the poly(A) tail and other sequences retained in the final circRNAs are shown in standard font. For consistency, final circRNAs are shown with the start of the CVB3 IRES as position 1, followed by the coding region, and then the other sequences included in the final circRNA. The two bases showing the circularization splice junction are in **blue**. A representative circRNA precursor is shown as a linear RNA with a placeholder for the open reading frame.

| RNA used | Sequence |
|---|---|
| linear nanoluciferase_3xF LAG_non-optimal | *AGGUAGUAUUCUUCUGGUCCCCACAGACUCAGAGAGAACCCGCCACCGCCACC* **AUGGUUUUCACCUUGGAAGAUUUCGUCGGGGAUUG**<br>**GCGUCAAACCGCAGGUUACAACCUCGAUCAGGUUUUGGAACAGGGCGGGGUAAGUUCUCUCUUCCAAAAUCUUGGAGUCUCCGUCAC**<br>**GCCUAUCCAGAGAAUUGUUUUGUCAGGCGAAAAUGGAUUGAAAAUUGAUAUUCAUGUCAUUAUACCUUAUGAGGGUCUGAGCGGGGA**<br>**CCAAAUGGGACAGAUUGAAAAGAUUUUCAAAGUCGUCUACCCAGUCGACGACCAUCAUUUCAAGGUGAUUUUGCAUUAUGGAACACUG**<br>**GUCAUUGAUGGAGUGACCCCUAACAUGAUAGACUACUUCGGCCGCCCCUAUGAGGGGAUAGCGGUGUUCGACGGUAAGAAAAUAACA**<br>**GUCACCGGCACUUUGUGGAAUGGCAAUAAAAUCAUCGAUGAAAGAUUGAUCAAUCCUGAUGGCUCACUGUUGUUUAGGGUUACAAUA**<br>**AACGGGGUAACGGGAUGGCGGCUCUGCGAACGGAUUCUUGCCGACUACAAAGACCAUGACGGUGAUUAUAAAGAUCAUGAUAUCGAU**<br>**UACAAGGAUGACGAUGACAAGUGAUAA***CUCGAGCUGGUACUGCAUGCACGCAAUGCUAGCUGCCCCUUUCCCGUCCUGGUACCCCG*<br>*AGUCUCCCCCGACCUCGGGUCCCAGGUAUGCUCCCACCUCCACCUGCCCCACUCACCACCU* AAAAAAAAAAAAAAAAAAAAAAAAAAAAA<br>AAAAAAAAAAAAAAAAAAAAAAAAAAAAAAAAAAAAAAAAAAAAAAAAAAAAAAAAAAAAAA |
| linear nanoluciferase_3xF LAG_optimal | *AGGUAGUAUUCUUCUGGUCCCCACAGACUCAGAGAGAACCCGCCACCGCCACC* **AUGGUGUUCACCCUGGAGGACUUCGUGGGCGACU**<br>**GGCGGCAGACCGCCGGCUACAACCUGGACCAGGUGCUGGAGCAGGGCGGAGUGAGCUCCCUGUUCCAGAACCUGGGCGUCAGCGUG**<br>**ACCCCCAUCCAGCGGAUCGUGCUGAGCGGCGAGAACGGCCUGAAGAUUGACAUCCACGUGAUCAUCCCAUACGAGGGCCUCAGCGGC**<br>**GACCAGAUGGGCCAGAUCGAGAAGAUCUUCAAGGUGGUCUACCCCGUGGACGACCACCACUUCAAGGUGAUCCUGCACUACGGCACC**<br>**CUGGUGAUCGACGGCGUGACCCCCAACAUGAUCGACUACUUCGGCAGACCCUACGAGGGCAUCGCCGUGUUCGAUGGCAAGAAGAUC**<br>**ACCGUGACCGGCACACUGUGGAACGGCAAUAAGAUCAUCGACGAGCGGCUGAUCAACCCAGACGGCAGCCUGCUGUUCAGAGUGACC**<br>**AUCAACGGCGUGACCGGCUGGCGGCUGUGCGAGCGGAUCCUGGCCGACUACAAGGACCACGACGGCGAUUAUAAGGACCACGACAUC**<br>**GACUACAAGGACGAUGACGACAAGUGAUAA***CUCGAGCUGGUACUGCAUGCACGCAAUGCUAGCUGCCCCUUUCCCGUCCUGGUACC*<br>*CCGAGUCUCCCCCGACCUCGGGUCCCAGGUAUGCUCCCACCUCCACCUGCCCCACUCACCACCU* AAAAAAAAAAAAAAAAAAAAAAAAAA<br>AAAAAAAAAAAAAAAAAAAAAAAAAAAAAAAAAAAAAAAAAAAAAAAAAAAAAAAAAAAAAA |
| linear nanoluciferase_3xF LAG_non-optimal+CL1/PEST | *AGGUAGUAUUCUUCUGGUCCCCACAGACUCAGAGAGAACCCGCCACCGCCACC* **AUGGUUUUCACCUUGGAAGAUUUCGUCGGGGAUUG**<br>**GCGUCAAACCGCAGGUUACAACCUCGAUCAGGUUUUGGAACAGGGCGGGGUAAGUUCUCUCUUCCAAAAUCUUGGAGUCUCCGUCAC**<br>**GCCUAUCCAGAGAAUUGUUUUGUCAGGCGAAAAUGGAUUGAAAAUUGAUAUUCAUGUCAUUAUACCUUAUGAGGGUCUGAGCGGGGA**<br>**CCAAAUGGGACAGAUUGAAAAGAUUUUCAAAGUCGUCUACCCAGUCGACGACCAUCAUUUCAAGGUGAUUUUGCAUUAUGGAACACUG**<br>**GUCAUUGAUGGAGUGACCCCUAACAUGAUAGACUACUUCGGCCGCCCCUAUGAGGGGAUAGCGGUGUUCGACGGUAAGAAAAUAACA**<br>**GUCACCGGCACUUUGUGGAAUGGCAAUAAAAUCAUCGAUGAAAGAUUGAUCAAUCCUGAUGGCUCACUGUUGUUUAGGGUUACAAUA**<br>**AACGGGGUAACGGGAUGGCGGCUCUGCGAACGGAUUCUUGCCGACUACAAAGACCAUGACGGUGAUUAUAAAGAUCAUGAUAUCGAU**<br>**UACAAGGAUGACGAUGACAAGAAUUCUGCUUGCAAGAACUGGUUCAGUAGCUUAAGCCACUUUGUGAUCCACCUUAACAGCCACGGC**<br>**UUUCCGCCUGAGGUUGAGGAACAGGCCGCCGGUACAUUGCCUAUGUCCUGCGCACAAGAAAGCGGUAUGGACCGGCACCCAGCCGCU**<br>**UGUGCUUCAGCUCGCAUCAACGUCUGAUAA***CUCGAGCUGGUACUGCAUGCACGCAAUGCUAGCUGCCCCUUUCCCGUCCUGGUACC*<br>*CCGAGUCUCCCCCGACCUCGGGUCCCAGGUAUGCUCCCACCUCCACCUGCCCCACUCACCACCU* AAAAAAAAAAAAAAAAAAAAAAAAAA<br>AAAAAAAAAAAAAAAAAAAAAAAAAAAAAAAAAAAAAAAAAAAAAAAAAAAAAAAAAAAAAA |

| | |
|---|---|
| linear nanoluciferase_3xFLAG_optimal+CL1/PEST | *AGGUAGUAUUCUUCUGGUCCCCACAGACUCAGAGAGAACCCGCCACCGCCACC* **AUGGUGUUCACCCUGGAGGACUUCGUGGGCGACU**<br>**GGCGGCAGACCGCCGGCUACAACCUGGACCAGGUGCUGGAGCAGGGCGGAGUGAGCUCCCUGUUCCAGAACCUGGGCGUCAGCGUG**<br>**ACCCCCAUCCAGCGGAUCGUGCUGAGCGGCGAGAACGGCCUGAAGAUUGACAUCCACGUGAUCAUCCCAUACGAGGGCCUCAGCGGC**<br>**GACCAGAUGGGCCAGAUCGAGAAGAUCUUCAAGGUGGUCUACCCCGUGGACGACCACCACUUCAAGGUGAUCCUGCACUACGGCACC**<br>**CUGGUGAUCGACGGCGUGACCCCCAACAUGAUCGACUACUUCGGCAGACCCUACGAGGGCAUCGCCGUGUUCGAUGGCAAGAAGAUC**<br>**ACCGUGACCGGCACACUGUGGAACGGCAAUAAGAUCAUCGACGAGCGGCUGAUCAACCCAGACGGCAGCCUGCUGUUCAGAGUGACC**<br>**AUCAACGGCGUGACCGGCUGGCGGCUGUGCGAGCGGAUCCUGGCCGACUACAAGGACCACGACGGCGAUUAUAAGGACCACGACAUC**<br>**GACUACAAGGACGAUGACGACAAGAACAGCGCUUGCAAGAACUGGUUCAGCAGCCUCAGCCACUUCGUGAUCCACCUGAACAGCCAC**<br>**GGCUUCCCCCCCGAGGUGGAGGAGCAGGCCGCCGGAACCCUGCCCAUGUCCUGCGCCCAGGAGAGCGGGAUGGACCGGCACCCAGC**<br>**CGCUUGCGCCAGCGCCCGGAUCAACGUCUGAUAA***CUCGAGCUGGUACUGCAUGCACGCAAUGCUAGCUGCCCCUUUCCCGUCCUGGG*<br>*UACCCCGAGUCUCCCCCGACCUCGGGUCCCAGGUAUGCUCCCACCUCCACCUGCCCCACUCACCACCU* AAAAAAAAAAAAAAAAAAAAAA<br>AAAAAAAAAAAAAAAAAAAAAAAAAAAAAAAAAAAAAAAAAAAAAAAAAAAAAAAAAAAAAAAAAAAAAAAAAAAAAAAAAAAAA |
| circular nanoluciferase_3xFLAG_non-optimal | UUAAAACAGCCUGUGGGUUGAUCCCACCCACAGGCCCAUUGGGCGCUAGCACUCUGGUAUCACGGUACCUUUGUGCGCCUGUUUUAUA<br>CCCCCUCCCCCAACUGUAACUUAGAAGUAACACACACCGAUCAACAGUCAGCGUGGCACACCAGCCACGUUUUGAUCAAGCACUUCUGU<br>UACCCCGGACUGAGUAUCAAUAGACUGCUCACGCGGUUGAAGGAGAAAGCGUUCGUUAUCCGGCCAACUACUUCGAAAAACCUAGUAAC<br>ACCGUGGAAGUUGCAGAGUGUUUCGCUCAGCACUACCCCAGUGUAGAUCAGGUCGAUGAGUCACCGCAUUCCCCACGGGCGACCGUGG<br>CGGUGGCUGCGUUGGCGGCCUGCCCAUGGGGAAACCCAUGGGACGCUCUAAUACAGACAUGGUGCGAAGAGUCUAUUGAGCUAGUUG<br>GUAGUCCUCCGGCCCCUGAAUGCGGCUAAUCCUAACUGCGGAGCACACACCCUCAAGCCAGAGGGCAGUGUGUCGUAACGGGCAACUC<br>UGCAGCGGAACCGACUACUUUGGGUGUCCGUGUUUCAUUUUAUUCCUAUACUGGCUGCUUAUGGUGACAAUUGAGAGAUCGUUACCAU<br>AUAGCUAUUGGAUUGGCCAUCCGGUGACUAAUAGAGCUAUUAUAUAUCCCUUUGUUGGGUUUAUACCACUUAGCUUGAAAGAGGUUAAA<br>ACAUUACAAUUCAUUGUUAAGUUGAAUACAGCAAA**AUGGUUUUCACCUUGGAAGAUUUCGUCGGGGAUUGGCGUCAAACCGCAGGUUA**<br>**CAACCUCGAUCAGGUUUUGGAACAGGGCGGGGUAAGUUCUCUCUUCCAAAAUCUUGGAGUCUCCGUCACGCCUAUCCAGAGAAUUGU**<br>**UUUGUCAGGCGAAAAUGGAUUGAAAAUUGAUAUUCAUGUCAUUAUACCUUAUGAGGGUCUGAGCGGGGACCAAAUGGGACAGAUUGA**<br>**AAAGAUUUUCAAAGUCGUCUACCCAGUCGACGACCAUCAUUUCAAGGUGAUUUUGCAUUAUGGAACACUGGUCAUUGAUGGAGUGAC**<br>**CCCUAACAUGAUAGACUACUUCGGCCGCCCCUAUGAGGGGAUAGCGGUGUUCGACGGUAAGAAAAUAACAGUCACCGGCACUUUGUG**<br>**GAAUGGCAAUAAAAUCAUCGAUGAAAGAUUGAUCAAUCCUGAUGGCUCACUGUUGUUUAGGGUUACAAUAAACGGGGUAACGGGAUG**<br>**GCGGCUCUGCGAACGGAUUCUUGCCGACUACAAAGACCAUGACGGUGAUUAUAAAGAUCAUGAUAUCGAUUACAAGGAUGACGAUGA**<br>**CAAGUGAUAA**CGGUAAAAAAAACAAAAAACAAAACGGCUAUUAUGCGUUACCGGCGAGACGCUACGGACUUAAAUAAU**UG**AAUGAAAAUC<br>CGUUGACCUUAAACGGUCGUGUGGGUUCAAGUCCCUCCACCCCCACGCCGGAAACGCAAUAGCCGAAAAACAAAAAACAAAAAAAACAAA<br>AAAAAAACCAAAAAAACAAAACACA |

| | |
|---|---|
| circular nanoluciferase_3xFLAG_optimal | UUAAAACAGCCUGUGGGUUGAUCCCACCCACAGGCCCAUUGGGCGCUAGCACUCUGGUAUCACGGUACCUUUGUGCGCCUGUUUUAUA<br>CCCCCUCCCCCAACUGUAACUUAGAAGUAACACACACCGAUCAACAGUCAGCGUGGCACACCAGCCACGUUUUGAUCAAGCACUUCUGU<br>UACCCCGGACUGAGUAUCAAUAGACUGCUCACGCGGUUGAAGGAGAAAGCGUUCGUUAUCCGGCCAACUACUUCGAAAAACCUAGUAAC<br>ACCGUGGAAGUUGCAGAGUGUUUCGCUCAGCACUACCCCAGUGUAGAUCAGGUCGAUGAGUCACCGCAUUCCCCACGGGCGACCGUGG<br>CGGUGGCUGCGUUGGCGGCCUGCCCAUGGGGAAACCCAUGGGACGCUCUAAUACAGACAUGGUGCGAAGAGUCUAUUGAGCUAGUUG<br>GUAGUCCUCCGGCCCCUGAAUGCGGCUAAUCCUAACUGCGGAGCACACACCCUCAAGCCAGAGGGCAGUGUGUCGUAACGGGCAACUC<br>UGCAGCGGAACCGACUACUUUGGGUGUCCGUGUUUCAUUUUAUUCCUAUACUGGCUGCUUAUGGUGACAAUUGAGAGAUCGUUACCAU<br>AUAGCUAUUGGAUUGGCCAUCCGGUGACUAAUAGAGCUAUUAUAUAUCCCUUUGUUGGGUUUAUACCACUUAGCUUGAAAGAGGUUAAA<br>ACAUUACAAUUCAUUGUUAAGUUGAAUACAGCAAA**AUGGUGUUCACCCUGGAGGACUUCGUGGGCGACUGGCGGCAGACCGCCGGCUA**<br>**CAACCUGGACCAGGUGCUGGAGCAGGGCGGAGUGAGCUCCCUGUUCCAGAACCUGGGCGUCAGCGUGACCCCCAUCCAGCGGAUCGU**<br>**GCUGAGCGGCGAGAACGGCCUGAAGAUUGACAUCCACGUGAUCAUCCCAUACGAGGGCCUCAGCGGCGACCAGAUGGGCCAGAUCGA**<br>**GAAGAUCUUCAAGGUGGUCUACCCCGUGGACGACCACCACUUCAAGGUGAUCCUGCACUACGGCACCCUGGUGAUCGACGGCGUGAC**<br>**CCCCAACAUGAUCGACUACUUCGGCAGACCCUACGAGGGCAUCGCCGUGUUCGAUGGCAAGAAGAUCACCGUGACCGGCACACUGUG**<br>**GAACGGCAAUAAGAUCAUCGACGAGCGGCUGAUCAACCCAGACGGCAGCCUGCUGUUCAGAGUGACCAUCAACGGCGUGACCGGCUG**<br>**GCGGCUGUGCGAGCGGAUCCUGGCC**GACUACAAGGACCACGACGGCGAUUAUAAGGACCACGACAUCGACUACAAGGACGAUGACGA<br>CAAG**UGAUAA**CGGUAAAAAAAAACAAAAAACAAAACGGCUAUUAUGCGUUACCGGCGAGACGCUACGGACUUAAAUAAUUGAAUGAAAAUC<br>CGUUGACCUUAAACGGUCGUGUGGGUUCAAGUCCCUCCACCCCCACGCCGGAAACGCAAUAGCCGAAAAACAAAAAACAAAAAAAACAAA<br>AAAAAAACCAAAAAAACAAAACACA |
| circular nanoluciferase_3xFLAG_non-optimal+CL1/PEST | UUAAAACAGCCUGUGGGUUGAUCCCACCCACAGGCCCAUUGGGCGCUAGCACUCUGGUAUCACGGUACCUUUGUGCGCCUGUUUUAUA<br>CCCCCUCCCCCAACUGUAACUUAGAAGUAACACACACCGAUCAACAGUCAGCGUGGCACACCAGCCACGUUUUGAUCAAGCACUUCUGU<br>UACCCCGGACUGAGUAUCAAUAGACUGCUCACGCGGUUGAAGGAGAAAGCGUUCGUUAUCCGGCCAACUACUUCGAAAAACCUAGUAAC<br>ACCGUGGAAGUUGCAGAGUGUUUCGCUCAGCACUACCCCAGUGUAGAUCAGGUCGAUGAGUCACCGCAUUCCCCACGGGCGACCGUGG<br>CGGUGGCUGCGUUGGCGGCCUGCCCAUGGGGAAACCCAUGGGACGCUCUAAUACAGACAUGGUGCGAAGAGUCUAUUGAGCUAGUUG<br>GUAGUCCUCCGGCCCCUGAAUGCGGCUAAUCCUAACUGCGGAGCACACACCCUCAAGCCAGAGGGCAGUGUGUCGUAACGGGCAACUC<br>UGCAGCGGAACCGACUACUUUGGGUGUCCGUGUUUCAUUUUAUUCCUAUACUGGCUGCUUAUGGUGACAAUUGAGAGAUCGUUACCAU<br>AUAGCUAUUGGAUUGGCCAUCCGGUGACUAAUAGAGCUAUUAUAUAUCCCUUUGUUGGGUUUAUACCACUUAGCUUGAAAGAGGUUAAA<br>ACAUUACAAUUCAUUGUUAAGUUGAAUACAGCAAA**AUGGUUUUCACCUUGGAAGAUUUCGUCGGGGAUUGGCGUCAAACCGCAGGUUA**<br>**CAACCUCGAUCAGGUUUUGGAACAGGGCGGGGUAAGUUCUCUCUUCCAAAAUCUUGGAGUCUCCGUCACGCCUAUCCAGAGAAUUGU**<br>**UUUGUCAGGCGAAAAUGGAUUGAAAAUUGAUAUUCAUGUCAUUAUACCUUAUGAGGGUCUGAGCGGGGACCAAAUGGGACAGAUUGA**<br>**AAAGAUUUUCAAAGUCGUCUACCCAGUCGACGACCAUCAUUUCAAGGUGAUUUUGCAUUAUGGAACACUGGUCAUUGAUGGAGUGAC**<br>**CCCUAACAUGAUAGACUACUUCGGCCGCCCCUAUGAGGGGAUAGCGGUGUUCGACGGUAAGAAAAUAACAGUCACCGGCACUUUGUG**<br>**GAAUGGCAAUAAAAUCAUCGAUGAAAGAUUGAUCAAUCCUGAUGGCUCACUGUUGUUUAGGGUUACAAUAAACGGGGUAACGGAUG**<br>**GCGGCUCUGCGAACGGAUUCUUGCCGACUACAAAGACCAUGACGGUGAUUAUAAAGAUCAUGAUAUCGAUUACAAGGAUGACGAUGA**<br>**CAAGAAUUCUGCUUGCAAGAACUGGUUCAGUAGCUUAAGCCACUUUGUGAUCCACCUUAACAGCCACGGCUUUCCGCCUGAGGUUGA**<br>**GGAACAGGCCGCCGGUACAUUGCCUAUGUCCUGCGCACAAGAAAGCGGUAUGGACCGGCACCCAGCCGCUUGUGCUUCAGCUCGCAU**<br>**CAACGUCUGAUAA**CGGUAAAAAAAAACAAAAAACAAAACGGCUAUUAUGCGUUACCGGCGAGACGCUACGGACUUAAAUAAUUGAAUGAAA<br>AUCCGUUGACCUUAAACGGUCGUGUGGGUUCAAGUCCCUCCACCCCCACGCCGGAAACGCAAUAGCCGAAAAACAAAAAACAAAAAAAAC<br>AAAAAAAAAACCAAAAAAACAAAACACA |

| | |
|---|---|
| circular nanoluciferase_3xFLAG_optimal+CL1/PEST | UUAAAACAGCCUGUGGGUUGAUCCCACCCACAGGCCCAUUGGGCGCUAGCACUCUGGUAUCACGGUACCUUUGUGCGCCUGUUUUAUA<br>CCCCCUCCCCCAACUGUAACUUAGAAGUAACACACACCGAUCAACAGUCAGCGUGGCACACCAGCCACGUUUUGAUCAAGCACUUCUGU<br>UACCCCGGACUGAGUAUCAAUAGACUGCUCACGCGGUUGAAGGAGAAAGCGUUCGUUAUCCGGCCAACUACUUCGAAAAACCUAGUAAC<br>ACCGUGGAAGUUGCAGAGUGUUUCGCUCAGCACUACCCCAGUGUAGAUCAGGUCGAUGAGUCACCGCAUUCCCCACGGGCGACCGUGG<br>CGGUGGCUGCGUUGGCGGCCUGCCCAUGGGGAAACCCAUGGGACGCUCUAAUACAGACAUGGUGCGAAGAGUCUAUUGAGCUAGUUG<br>GUAGUCCUCCGGCCCCUGAAUGCGGCUAAUCCUAACUGCGGAGCACACACCCUCAAGCCAGAGGGCAGUGUGUCGUAACGGGCAACUC<br>UGCAGCGGAACCGACUACUUUGGGUGUCCGUGUUUCAUUUUAUUCCUAUACUGGCUGCUUAUGGUGACAAUUGAGAGAUCGUUACCAU<br>AUAGCUAUUGGAUUGGCCAUCCGGUGACUAAUAGAGCUAUUAUAUCCCUUUGUUGGGUUUAUACCACUUAGCUUGAAAGAGGUUAAA<br>ACAUUACAAUUCAUUGUUAAGUUGAAUACAGCAAA**AUGAUGGUGUUCACCCUGGAGGACUUCGUGGGCGACUGGCGGCAGACCGCCGG**<br>**CUACAACCUGGACCAGGUGCUGGAGCAGGGCGGAGUGAGCUCCCUGUUCCAGAACCUGGGCGUCAGCGUGACCCCCAUCCAGCGGAU**<br>**CGUGCUGAGCGGCGAGAACGGCCUGAAGAUUGACAUCCACGUGAUCAUCCCAUACGAGGGCCUCAGCGGCGACCAGAUGGGCCAGAU**<br>**CGAGAAGAUCUUCAAGGUGGUCUACCCCGUGGACGACCACCACUUCAAGGUGAUCCUGCACUACGGCACCCUGGUGAUCGACGGCGU**<br>**GACCCCCAACAUGAUCGACUACUUCGGCAGACCCUACGAGGGCAUCGCCGUGUUCGAUGGCAAGAAGAUCACCGUGACCGGCACACU**<br>**GUGGAACGGCAAUAAGAUCAUCGACGAGCGGCUGAUCAACCCAGACGGCAGCCUGCUGUUCAGAGUGACCAUCAACGGCGUGACCGG**<br>**CUGGCGGCUGUGCGAGCGGAUCCUGGCC**GACUACAAGGACCACGACGGCGAUUAUAAGGACCACGACAUCGACUACAAGGACGAUGA<br>CGACAAGAACAGCGCUUGCAAGAACUGGUUCAGCAGCCUCAGCCACUUCGUGAUCCACCUGAACAGCCACGGCUUCCCCCCCGAGGU<br>GGAGGAGCAGGCCGCCGGAACCCUGCCCAUGUCCUGCGCCCAGGAGAGCGGGAUGGACCGGCACCCAGCCGCUUGCGCCAGCGCCC<br>GGAUCAACGUC**UGAUAA**CGGUAAAAAAAAACAAAAAACAAAACGGCUAUUAUGCGUUACCGGCGAGACGCUACGGACUUAAAUAAUUGAAU<br>GAAAAUCCGUUGACCUUAAACGGUCGUGUGGGUUCAAGUCCCUCCACCCCCACGCCGGAAACGCAAUAGCCGAAAAACAAAAAACAAAAA<br>AAACAAAAAAAAAAACCAAAAAAACAAAACACA |
| linear *Renilla* luciferase_non-optimal | *AGGUAGUAUUCUUCUGGUCCCCACAGACUCAGAGAGAACCCGCCACCGCCACC* **AUGGCUUCGAAAGUUUAUGAUCCAGAACAAAGGAA**<br>**ACGGAUGAUAACUGGUCCGCAGUGGUGGGCCAGAUGUAAACAAAUGAAUGUUCUUGAUUCAUUUAUUAAUUAUUAUGAUUCAGAAAAA**<br>**CAUGCAGAAAAUGCUGUUAUUUUUUUACAUGGUAACGCGGCCUCUUCUUAUUUAUGGCGACAUGUUGUGCCACAUAUUGAGCCAGUA**<br>**GCGCGGUGUAUUAUACCAGACCUUAUUGGUAUGGGCAAAUCAGGCAAAUCUGGUAAUGGUUCUUAUAGGUUACUUGAUCAUUACAAA**<br>**UAUCUUACUGCAUGGUUUGAACUUCUUAAUUUACCAAAGAAGAUCAUUUUUGUCGGCCAUGAUUGGGGUGCUUGUUUGGCAUUUCAU**<br>**UAUAGCUAUGAGCAUCAAGAUAAGAUCAAAGCAAUAGUUCACGCUGAAAGUGUAGUAGAUGUGAUUGAAUCAUGGGAUGAAUGGCCU**<br>**GAUAUUGAAGAAGAUAUUGCGUUGAUCAAAUCUGAAGAAGGAGAAAAAAUGGUUUUGGAGAAUAACUUCUUCGUGGAAACCAUGUUG**<br>**CCAUCAAAAAUCAUGAGAAAGUUAGAACCAGAAGAAUUUGCAGCAUAUCUUGAACCAUUCAAAGAGAAAGGUGAAGUUCGUCGUCCAA**<br>**CAUUAUCAUGGCCUCGUGAAAUCCCGUUAGUAAAAGGUGGUAAACCUGACGUUGUACAAAUUGUUAGGAAUUAUAAUGCUUAUCUACG**<br>**UGCAAGUGAUGAUUUACCAAAAAUGUUUAUUGAAUCGGACCCAGGAUUCUUUUCCAAUGCUAUUGUUGAAGGUGCCAAGAAGUUUCC**<br>**UAAUACUGAAUUUGUCAAAGUAAAAGGUCUUCAUUUUUCGCAAGAAGAUGCACCUGAUGAAAUGGGAAAAUAUAUCAAAUCGUUCGUU**<br>**GAGCGAGUUCUCAAAAAUGAACAAUGAUAA***CUCGAGCUGGUACUGCAUGCACGCAAUGCUAGCUGCCCCUUUCCCGUCCUGGGUACC*<br>*CCGAGUCUCCCCCGACCUCGGGUCCCAGGUAUGCUCCCACCUCCACCUGCCCCACUCACCACCU* AAAAAAAAAAAAAAAAAAAAAAAAA<br>AAAAAAAAAAAAAAAAAAAAAAAAAAAAAAAAAAAAAAAAAAAAAAAAAAAAAAAAAAAAA |

| | |
|---|---|
| linear *Renilla* luciferase_optimal | *AGGUAGUAUUCUUCUGGUCCCCACAGACUCAGAGAGAACCCGCCACCGCCACC* **AUGGCCAGCAAGGUGUACGACCCCGAGCAGCGGAAGCGGAUGAUCACCGGCCCCCAGUGGUGGGCCCGGUGCAAGCAGAUGAACGUGCUGGACAGCUUCAUCAACUACUACGACAGCGAGAAGCACGCCGAGAACGCCGUGAUCUUCCUGCACGGCAACGCUGCCUCCAGCUACCUGUGGCGGCACGUGGUGCCACACAUCGAGCCCGUGGCCCGGUGCAUCAUUCCAGACCUGAUCGGCAUGGGCAAGAGCGGAAAGAGCGGCAACGGCAGCUACCGGCUGCUGGACCACUACAAGUACCUGACCGCCUGGUUCGAGCUGCUGAACCUGCCCAAGAAGAUCAUCUUCGUGGGCCACGACUGGGGCGCCUGCCUCGCUUUCCACUACAGCUACGAGCACCAGGACAAGAUCAAGGCCAUCGUGCACGCCGAGAGCGUGGUCGACGUGAUCGAGAGCUGGGACGAGUGGCCCGACAUCGAGGAGGACAUCGCCCUGAUCAAGAGCGAGGAGGGCGAGAAGAUGGUGCUGGAGAAUAACUUCUUCGUGGAGACAAUGCUGCCAAGCAAGAUCAUGAGAAAGCUGGAGCCCGAGGAGUUCGCCGCUUACCUGGAACCCUUCAAGGAGAAGGGCGAGGUGAGACGGCCCACCCUGAGCUGGCCAAGAGAAAUCCCCCUGGUGAAGGGCGGCAAGCCCGACGUGGUCCAGAUCGUGCGGAACUACAACGCCUACCUGCGGGCCAGCGACGACCUGCCAAAGAUGUUCAUCGAGAGCGACCCCGGCUUCUUUAGCAACGCCAUCGUGGAGGGCGCCAAGAAAUUCCCCAACACCGAGUUCGUGAAGGUGAAGGGCCUGCAUUUCUCCCAGGAGGAUGCUCCCGACGAGAUGGGCAAGUACAUUAAGUCCUUCGUGGAGCGGGUGCUGAAGAACGAGCAGUGAUAA***CUCGAGCUGGUACUGCAUGCACGCAAUGCUAGCUGCCCCUUUCCCGUCCUGGGUACCCCGAGUCUCCCCCGACCUCGGGUCCCAGGUAUGCUCCCACCUCCACCUGCCCCACUCACCACCU* AAAAAAAAAAAAAAAAAAAAAAAAAAAAAAAAAAAAAAAAAAAAAAAAAAAAAAAAAAAAAAAAAAAAAAAAAAAAAAAAAAAAAAAAAAA |
| linear *Renilla* luciferase_non-optimal+CL1/PEST | *AGGUAGUAUUCUUCUGGUCCCCACAGACUCAGAGAGAACCCGCCACCGCCACC* **AUGGCUUCGAAAGUUUAUGAUCCAGAACAAAGGAAACGGAUGAUAACUGGUCCGCAGUGGUGGGCCAGAUGUAAACAAAUGAAUGUUCUUGAUUCAUUUAUUAAUUAUUAUGAUUCAGAAAAACAUGCAGAAAAUGCUGUUAUUUUUUUACAUGGUAACGCGGCCUCUUCUUAUUUAUGGCGACAUGUUGUGCCACAUAUUGAGCCAGUAGCGCGGUGUAUUAUACCAGACCUUAUUGGUAUGGGCAAAUCAGGCAAAUCUGGUAAUGGUUCUUAUAGGUUACUUGAUCAUUACAAAUAUCUUACUGCAUGGUUUGAACUUCUUAAUUUACCAAAGAAGAUCAUUUUUGUCGGCCAUGAUUGGGGUGCUUGUUUGGCAUUUCAUUAUAGCUAUGAGCAUCAAGAUAAGAUCAAAGCAAUAGUUCACGCUGAAAGUGUAGUAGAUGUGAUUGAAUCAUGGGAUGAAUGGCCUGAUAUUGAAGAAGAUAUUGCGUUGAUCAAAUCUGAAGAAGGAGAAAAAAUGGUUUUGGAGAAUAACUUCUUCGUGGAAACCAUGUUGCCAUCAAAAAUCAUGAGAAAGUUAGAACCAGAAGAAUUUGCAGCAUAUCUUGAACCAUUCAAAGAGAAAGGUGAAGUUCGUCGUCCAACAUUAUCAUGGCCUCGUGAAAUCCCGUUAGUAAAAGGUGGUAAACCUGACGUUGUACAAAUUGUUAGGAAUUAUAAUGCUUAUCUACGUGCAAGUGAUGAUUUACCAAAAAUGUUUAUUGAAUCGGACCCAGGAUUCUUUUCCAAUGCUAUUGUUGAAGGUGCCAAGAAGUUUCCUAAUACUGAAUUUGUCAAAGUAAAAGGUCUUCAUUUUUCGCAAGAAGAUGCACCUGAUGAAAUGGGAAAAUAUAUCAAAUCGUUCGUUGAGCGAGUUCUCAAAAAUGAACAAAAUUCUGCUUGCAAGAACUGGUUCAGUAGCUUAAGCCACUUUGUGAUCCACCUUAACAGCCACGGCUUUCCGCCUGAGGUUGAGGAACAGGCCGCCGGUACAUUGCCUAUGUCCUGCGCACAAGAAAGCGGUAUGGACCGGCACCCAGCCGCUUGUGCUUCAGCUCGCAUCAACGUCUGAUAA***CUCGAGCUGGUACUGCAUGCACGCAAUGCUAGCUGCCCCUUUCCCGUCCUGGGUACCCCGAGUCUCCCCCGACCUCGGGUCCCAGGUAUGCUCCCACCUCCACCUGCCCCACUCACCACCU* AAAAAAAAAAAAAAAAAAAAAAAAAAAAAAAAAAAAAAAAAAAAAAAAAAAAAAAAAAAAAAAAAAAAAAAAAAAAAAAAAAAAAAAAAAA |

| | |
|---|---|
| linear *Renilla* luciferase_optimal+ CL1/PEST | *AGGUAGUAUUCUUCUGGUCCCCACAGACUCAGAGAGAACCCGCCACCGCCACC* **AUGGCCAGCAAGGUGUACGACCCCGAGCAGCGGAA<br>GCGGAUGAUCACCGGCCCCCAGUGGUGGGCCCGGUGCAAGCAGAUGAACGUGCUGGACAGCUUCAUCAACUACUACGACAGCGAGAA<br>GCACGCCGAGAACGCCGUGAUCUUCCUGCACGGCAACGCUGCCUCCAGCUACCUGUGGCGGCACGUGGUGCCACACAUCGAGCCCGU<br>GGCCCGGUGCAUCAUUCCAGACCUGAUCGGCAUGGGCAAGAGCGGAAAGAGCGGCAACGGCAGCUACCGGCUGCUGGACCACUACAA<br>GUACCUGACCGCCUGGUUCGAGCUGCUGAACCUGCCCAAGAAGAUCAUCUUCGUGGGCCACGACUGGGGCGCCUGCCUCGCUUUCCA<br>CUACAGCUACGAGCACCAGGACAAGAUCAAGGCCAUCGUGCACGCCGAGAGCGUGGUCGACGUGAUCGAGAGCUGGGACGAGUGGCC<br>CGACAUCGAGGAGGACAUCGCCCUGAUCAAGAGCGAGGAGGGCGAGAAGAUGGUGCUGGAGAAUAACUUCUUCGUGGAGACAAUGCU<br>GCCAAGCAAGAUCAUGAGAAAGCUGGAGCCCGAGGAGUUCGCCGCUUACCUGGAACCCUUCAAGGAGAAGGGCGAGGUGAGACGGCC<br>CACCCUGAGCUGGCCAAGAGAAAUCCCCCUGGUGAAGGGCGGCAAGCCCGACGUGGUCCAGAUCGUGCGGAACUACAACGCCUACCU<br>GCGGGCCAGCGACGACCUGCCAAAGAUGUUCAUCGAGAGCGACCCCGGCUUCUUUAGCAACGCCAUCGUGGAGGGCGCCAAGAAAUU<br>CCCCAACACCGAGUUCGUGAAGGUGAAGGGCCUGCAUUUCUCCCAGGAGGAUGCUCCCGACGAGAUGGGCAAGUACAUUAAGUCCUU<br>CGUGGAGCGGGUGCUGAAGAACGAGCAGAACAGCGCUUGCAAGAACUGGUUCAGCAGCCUCAGCCACUUCGUGAUCCACCUGAACAG<br>CCACGGCUUCCCCCCCGAGGUGGAGGAGCAGGCCGCCGGAACCCUGCCCAUGUCCUGCGCCCAGGAGAGCGGGAUGGACCGGCACC<br>CAGCCGCUUGCGCCAGCGCCCGGAUCAACGUCUGAUAA***CUCGAGCUGGUACUGCAUGCACGCAAUGCUAGCUGCCCCUUUCCCGUCC<br>UGGGUACCCCGAGUCUCCCCCGACCUCGGGUCCCAGGUAUGCUCCCACCUCCACCUGCCCCACUCACCACCU* AAAAAAAAAAAAAAAA<br>AAAAAAAAAAAAAAAAAAAAAAAAAAAAAAAAAAAAAAAAAAAAAAAAAAAAAAAAAAAAAAAAAAAAAAAAAAAAAA |

| | |
|---|---|
| circular *Renilla* luciferase_non-optimal | UUAAAACAGCCUGUGGGUUGAUCCCACCCACAGGCCCAUUGGGCGCUAGCACUCUGGUAUCACGGUACCUUUGUGCGCCUGUUUUAUA<br>CCCCCUCCCCCAACUGUAACUUAGAAGUAACACACACCGAUCAACAGUCAGCGUGGCACACCAGCCACGUUUUGAUCAAGCACUUCUGU<br>UACCCCGGACUGAGUAUCAAUAGACUGCUCACGCGGUUGAAGGAGAAAGCGUUCGUUAUCCGGCCAACUACUUCGAAAAACCUAGUAAC<br>ACCGUGGAAGUUGCAGAGUGUUUCGCUCAGCACUACCCCAGUGUAGAUCAGGUCGAUGAGUCACCGCAUUCCCCACGGGCGACCGUGG<br>CGGUGGCUGCGUUGGCGGCCUGCCCAUGGGGAAACCCAUGGGACGCUCUAAUACAGACAUGGUGCGAAGAGUCUAUUGAGCUAGUUG<br>GUAGUCCUCCGGCCCCUGAAUGCGGCUAAUCCUAACUGCGGAGCACACACCCUCAAGCCAGAGGGCAGUGUGUCGUAACGGGCAACUC<br>UGCAGCGGAACCGACUACUUUGGGUGUCCGUGUUUCAUUUUAUUCCUAUACUGGCUGCUUAUGGUGACAAUUGAGAGAUCGUUACCAU<br>AUAGCUAUUGGAUUGGCCAUCCGGUGACUAAUAGAGCUAUUAUAUAUCCCUUUGUUGGGUUUAUACCACUUAGCUUGAAAGAGGUUAAA<br>ACAUUACAAUUCAUUGUUAAGUUGAAUACAGCAAA**AUGGCUUCGAAAGUUUAUGAUCCAGAACAAAGGAAACGGAUGAUAACUGGUCC**<br>**GCAGUGGUGGGCCAGAUGUAAACAAAUGAAUGUUCUUGAUUCAUUUAUUAAUUAUUAUGAUUCAGAAAAACAUGCAGAAAAUGCUGUU**<br>**AUUUUUUUACAUGGUAACGCGGCCUCUUCUUAUUUAUGGCGACAUGUUGUGCCACAUAUUGAGCCAGUAGCGCGGUGUAUUAUACCA**<br>**GACCUUAUUGGUAUGGGCAAAUCAGGCAAAUCUGGUAAUGGUUCUUAUAGGUUACUUGAUCAUUACAAAUAUCUUACUGCAUGGUUU**<br>**GAACUUCUUAAUUUACCAAAGAAGAUCAUUUUUGUCGGCCAUGAUUGGGGUGCUUGUUUGGCAUUUCAUUAUAGCUAUGAGCAUCAA**<br>**GAUAAGAUCAAAGCAAUAGUUCACGCUGAAAGUGUAGUAGAUGUGAUUGAAUCAUGGGAUGAAUGGCCUGAUAUUGAAGAAGAUAUU**<br>**GCGUUGAUCAAAUCUGAAGAAGGAGAAAAAAUGGUUUUGGAGAAUAACUUCUUCGUGGAAACCAUGUUGCCAUCAAAAAUCAUGAGAA**<br>**AGUUAGAACCAGAAGAAUUUGCAGCAUAUCUUGAACCAUUCAAAGAGAAAGGUGAAGUUCGUCGUCCAACAUUAUCAUGGCCUCGUGA**<br>**AAUCCCGUUAGUAAAAGGUGGUAAACCUGACGUUGUACAAAUUGUUAGGAAUUAUAAUGCUUAUCUACGUGCAAGUGAUGAUUUACCA**<br>**AAAAUGUUUAUUGAAUCGGACCCAGGAUUCUUUUCCAAUGCUAUUGUUGAAGGUGCCAAGAAGUUUCCUAAUACUGAAUUUGUCAAAG**<br>**UAAAAGGUCUUCAUUUUUCGCAAGAAGAUGCACCUGAUGAAAUGGGAAAAUAUAUCAAAUCGUUCGUUGAGCGAGUUCUCAAAAAUGA**<br>**ACAAUGAUAA**CGGUAAAAAAAACAAAAAACAAAACGGCUAUUAUGCGUUACCGGCGAGACGCUACGGACUUAAAUAAU**UG**AAUGAAAAUC<br>CGUUGACCUUAAACGGUCGUGUGGGUUCAAGUCCCUCCACCCCCACGCCGGAAACGCAAUAGCCGAAAAACAAAAAACAAAAAAAACAAA<br>AAAAAAACCAAAAAAACAAAACACA |

| | |
|---|---|
| circular *Renilla* luciferase_optimal | UUAAAACAGCCUGUGGGUUGAUCCCACCCACAGGCCCAUUGGGCGCUAGCACUCUGGUAUCACGGUACCUUUGUGCGCCUGUUUUAUA<br>CCCCCUCCCCCAACUGUAACUUAGAAGUAACACACACCGAUCAACAGUCAGCGUGGCACACCAGCCACGUUUUGAUCAAGCACUUCUGU<br>UACCCCGGACUGAGUAUCAAUAGACUGCUCACGCGGUUGAAGGAGAAAGCGUUCGUUAUCCGGCCAACUACUUCGAAAAACCUAGUAAC<br>ACCGUGGAAGUUGCAGAGUGUUUCGCUCAGCACUACCCCAGUGUAGAUCAGGUCGAUGAGUCACCGCAUUCCCCACGGGCGACCGUGG<br>CGGUGGCUGCGUUGGCGGCCUGCCCAUGGGGAAACCCAUGGGACGCUCUAAUACAGACAUGGUGCGAAGAGUCUAUUGAGCUAGUUG<br>GUAGUCCUCCGGCCCCUGAAUGCGGCUAAUCCUAACUGCGGAGCACACACCCUCAAGCCAGAGGGCAGUGUGUCGUAACGGGCAACUC<br>UGCAGCGGAACCGACUACUUUGGGUGUCCGUGUUUCAUUUUAUUCCUAUACUGGCUGCUUAUGGUGACAAUUGAGAGAUCGUUACCAU<br>AUAGCUAUUGGAUUGGCCAUCCGGUGACUAAUAGAGCUAUUAUAUAUCCCUUUGUUGGGUUUAUACCACUUAGCUUGAAAGAGGUUAAA<br>ACAUUACAAUUCAUUGUUAAGUUGAAUACAGCAAA**AUGGCCAGCAAGGUGUACGACCCCGAGCAGCGGAAGCGGAUGAUCACCGGCCC**<br>**CCAGUGGUGGGCCCGGUGCAAGCAGAUGAACGUGCUGGACAGCUUCAUCAACUACUACGACAGCGAGAAGCACGCCGAGAACGCCGU**<br>**GAUCUUCCUGCACGGCAACGCUGCCUCCAGCUACCUGUGGCGGCACGUGGUGCCACACAUCGAGCCCGUGGCCCGGUGCAUCAUUCC**<br>**AGACCUGAUCGGCAUGGGCAAGAGCGGAAAGAGCGGCAACGGCAGCUACCGGCUGCUGGACCACUACAAGUACCUGACCGCCUGGUU**<br>**CGAGCUGCUGAACCUGCCCAAGAAGAUCAUCUUCGUGGGCCACGACUGGGGCGCCUGCCUCGCUUUCCACUACAGCUACGAGCACCA**<br>**GGACAAGAUCAAGGCCAUCGUGCACGCCGAGAGCGUGGUCGACGUGAUCGAGAGCUGGGACGAGUGGCCCGACAUCGAGGAGGACA**<br>**UCGCCCUGAUCAAGAGCGAGGAGGGCGAGAAGAUGGUGCUGGAGAAUAACUUCUUCGUGGAGACAAUGCUGCCAAGCAAGAUCAUGA**<br>**GAAAGCUGGAGCCCGAGGAGUUCGCCGCUUACCUGGAACCCUUCAAGGAGAAGGGCGAGGUGAGACGGCCCACCCUGAGCUGGCCAA**<br>**GAGAAAUCCCCCUGGUGAAGGGCGGCAAGCCCGACGUGGUCCAGAUCGUGCGGAACUACAACGCCUACCUGCGGGCCAGCGACGACC**<br>**UGCCAAAGAUGUUCAUCGAGAGCGACCCCGGCUUCUUUAGCAACGCCAUCGUGGAGGGCGCCAAGAAAUUCCCCAACACCGAGUUCG**<br>**UGAAGGUGAAGGGCCUGCAUUUCUCCCAGGAGGAUGCUCCCGACGAGAUGGGCAAGUACAUUAAGUCCUUCGUGGAGCGGGUGCUG**<br>**AAGAACGAGCAGUGAUAA**CGGUAAAAAAAACAAAAAACAAAACGGCUAUUAUGCGUUACCGGCGAGACGCUACGGACUUAAAUAAUUGA<br>AUGAAAAUCCGUUGACCUUAAACGGUCGUGUGGGUUCAAGUCCCUCCACCCCCACGCCGGAAACGCAAUAGCCGAAAAACAAAAAACAAA<br>AAAAACAAAAAAAAAACCAAAAAAACAAAACACA |

| | |
|---|---|
| circular *Renilla* luciferase_non-optimal+CL1/PEST | UUAAAACAGCCUGUGGGUUGAUCCCACCCACAGGCCCAUUGGGCGCUAGCACUCUGGUAUCACGGUACCUUUGUGCGCCUGUUUUAUA<br>CCCCCUCCCCCAACUGUAACUUAGAAGUAACACACACCGAUCAACAGUCAGCGUGGCACACCAGCCACGUUUUGAUCAAGCACUUCUGU<br>UACCCCGGACUGAGUAUCAAUAGACUGCUCACGCGGUUGAAGGAGAAAGCGUUCGUUAUCCGGCCAACUACUUCGAAAAACCUAGUAAC<br>ACCGUGGAAGUUGCAGAGUGUUUCGCUCAGCACUACCCCAGUGUAGAUCAGGUCGAUGAGUCACCGCAUUCCCCACGGGCGACCGUGG<br>CGGUGGCUGCGUUGGCGGCCUGCCCAUGGGGAAACCCAUGGGACGCUCUAAUACAGACAUGGUGCGAAGAGUCUAUUGAGCUAGUUG<br>GUAGUCCUCCGGCCCCUGAAUGCGGCUAAUCCUAACUGCGGAGCACACACCCUCAAGCCAGAGGGCAGUGUGUCGUAACGGGCAACUC<br>UGCAGCGGAACCGACUACUUUGGGUGUCCGUGUUUCAUUUUAUUCCUAUACUGGCUGCUUAUGGUGACAAUUGAGAGAUCGUUACCAU<br>AUAGCUAUUGGAUUGGCCAUCCGGUGACUAAUAGAGCUAUUAUAUAUCCCUUUGUUGGGUUUAUACCACUUAGCUUGAAAGAGGUUAAA<br>ACAUUACAAUUCAUUGUUAAGUUGAAUACAGCAAA**AUGGCUUCGAAAGUUUAUGAUCCAGAACAAAGGAAACGGAUGAUAACUGGUCC**<br>**GCAGUGGUGGGCCAGAUGUAAACAAAUGAAUGUUCUUGAUUCAUUUAUUAAUUAUUAUGAUUCAGAAAAACAUGCAGAAAAUGCUGUU**<br>**AUUUUUUUACAUGGUAACGCGGCCUCUUCUUAUUUAUGGCGACAUGUUGUGCCACAUAUUGAGCCAGUAGCGCGGUGUAUUAUACCA**<br>**GACCUUAUUGGUAUGGGCAAAUCAGGCAAAUCUGGUAAUGGUUCUUAUAGGUUACUUGAUCAUUACAAAUAUCUUACUGCAUGGUUU**<br>**GAACUUCUUAAUUUACCAAAGAAGAUCAUUUUUGUCGGCCAUGAUUGGGGUGCUUGUUUGGCAUUUCAUUAUAGCUAUGAGCAUCAA**<br>**GAUAAGAUCAAAGCAAUAGUUCACGCUGAAAGUGUAGUAGAUGUGAUUGAAUCAUGGGAUGAAUGGCCUGAUAUUGAAGAAGAUAUU**<br>**GCGUUGAUCAAAUCUGAAGAAGGAGAAAAAAUGGUUUUGGAGAAUAACUUCUUCGUGGAAACCAUGUUGCCAUCAAAAAUCAUGAGAA**<br>**AGUUAGAACCAGAAGAAUUUGCAGCAUAUCUUGAACCAUUCAAAGAGAAAGGUGAAGUUCGUCGUCCAACAUUAUCAUGGCCUCGUGA**<br>**AAUCCCGUUAGUAAAAGGUGGUAAACCUGACGUUGUACAAAUUGUUAGGAAUUAUAAUGCUUAUCUACGUGCAAGUGAUGAUUUACCA**<br>**AAAAUGUUUAUUGAAUCGGACCCAGGAUUCUUUUCCAAUGCUAUUGUUGAAGGUGCCAAGAAGUUUCCUAAUACUGAAUUUGUCAAAG**<br>**UAAAAGGUCUUCAUUUUUCGCAAGAAGAUGCACCUGAUGAAAUGGGAAAAUAUAUCAAAUCGUUCGUUGAGCGAGUUCUCAAAAAUGA**<br>**ACAAAAUUCUGCUUGCAAGAACUGGUUCAGUAGCUUAAGCCACUUUGUGAUCCACCUUAACAGCCACGGCUUUCCGCCUGAGGUUGA**<br>**GGAACAGGCCGCCGGUACAUUGCCUAUGUCCUGCGCACAAGAAAGCGGUAUGGACCGGCACCCAGCCGCUUGUGCUUCAGCUCGCAU**<br>**CAACGUCUGAUAA**CGGUAAAAAAAACAAAAAACAAAACGGCUAUUAUGCGUUACCGGCGAGACGCUACGGACUUAAAUAAUUGAAUGAAA<br>AUCCGUUGACCUUAAACGGUCGUGUGGGUUCAAGUCCCUCCACCCCCACGCCGGAAACGCAAUAGCCGAAAAACAAAAAACAAAAAAAAC<br>AAAAAAAAAACCAAAAAAACAAAACACA |

| | |
|---|---|
| circular *Renilla* luciferase_optimal+ CL1/PEST | UUAAAACAGCCUGUGGGUUGAUCCCACCCACAGGCCCAUUGGGCGCUAGCACUCUGGUAUCACGGUACCUUUGUGCGCCUGUUUUAUA<br>CCCCCUCCCCCAACUGUAACUUAGAAGUAACACACACCGAUCAACAGUCAGCGUGGCACACCAGCCACGUUUUGAUCAAGCACUUCUGU<br>UACCCCGGACUGAGUAUCAAUAGACUGCUCACGCGGUUGAAGGAGAAAGCGUUCGUUAUCCGGCCAACUACUUCGAAAAACCUAGUAAC<br>ACCGUGGAAGUUGCAGAGUGUUUCGCUCAGCACUACCCCAGUGUAGAUCAGGUCGAUGAGUCACCGCAUUCCCCACGGGCGACCGUGG<br>CGGUGGCUGCGUUGGCGGCCUGCCCAUGGGGAAACCCAUGGGACGCUCUAAUACAGACAUGGUGCGAAGAGUCUAUUGAGCUAGUUG<br>GUAGUCCUCCGGCCCCUGAAUGCGGCUAAUCCUAACUGCGGAGCACACACCCUCAAGCCAGAGGGCAGUGUGUCGUAACGGGCAACUC<br>UGCAGCGGAACCGACUACUUUGGGUGUCCGUGUUUCAUUUUAUUCCUAUACUGGCUGCUUAUGGUGACAAUUGAGAGAUCGUUACCAU<br>AUAGCUAUUGGAUUGGCCAUCCGGUGACUAAUAGAGCUAUUAUAUAUCCCUUUGUUGGGUUUAUACCACUUAGCUUGAAAGAGGUUAAA<br>ACAUUACAAUUCAUUGUUAAGUUGAAUACAGCAAA**AUGGCCAGCAAGGUGUACGACCCCGAGCAGCGGAAGCGGAUGAUCACCGGCCC**<br>**CCAGUGGUGGGCCCGGUGCAAGCAGAUGAACGUGCUGGACAGCUUCAUCAACUACUACGACAGCGAGAAGCACGCCGAGAACGCCGU**<br>**GAUCUUCCUGCACGGCAACGCUGCCUCCAGCUACCUGUGGCGGCACGUGGUGCCACACAUCGAGCCCGUGGCCCGGUGCAUCAUUCC**<br>**AGACCUGAUCGGCAUGGGCAAGAGCGGAAAGAGCGGCAACGGCAGCUACCGGCUGCUGGACCACUACAAGUACCUGACCGCCUGGUU**<br>**CGAGCUGCUGAACCUGCCCAAGAAGAUCAUCUUCGUGGGCCACGACUGGGGCGCCUGCCUCGCUUUCCACUACAGCUACGAGCACCA**<br>**GGACAAGAUCAAGGCCAUCGUGCACGCCGAGAGCGUGGUCGACGUGAUCGAGAGCUGGGACGAGUGGCCCGACAUCGAGGAGGACA**<br>**UCGCCCUGAUCAAGAGCGAGGAGGGCGAGAAGAUGGUGCUGGAGAAUAACUUCUUCGUGGAGACAAUGCUGCCAAGCAAGAUCAUGA**<br>**GAAAGCUGGAGCCCGAGGAGUUCGCCGCUUACCUGGAACCCUUCAAGGAGAAGGGCGAGGUGAGACGGCCCACCCUGAGCUGGCCAA**<br>**GAGAAAUCCCCCUGGUGAAGGGCGGCAAGCCCGACGUGGUCCAGAUCGUGCGGAACUACAACGCCUACCUGCGGGCCAGCGACGACC**<br>**UGCCAAAGAUGUUCAUCGAGAGCGACCCCGGCUUCUUUAGCAACGCCAUCGUGGAGGGCGCCAAGAAAUUCCCCAACACCGAGUUCG**<br>**UGAAGGUGAAGGGCCUGCAUUUCUCCCAGGAGGAUGCUCCCGACGAGAUGGGCAAGUACAUUAAGUCCUUCGUGGAGCGGGUGCUG**<br>**AAGAACGAGCAGAACAGCGCUUGCAAGAACUGGUUCAGCAGCCUCAGCCACUUCGUGAUCCACCUGAACAGCCACGGCUUCCCCCCC**<br>**GAGGUGGAGGAGCAGGCCGCCGGAACCCUGCCCAUGUCCUGCGCCCAGGAGAGCGGGAUGGACCGGCACCCAGCCGCUUGCGCCAG**<br>**CGCCCGGAUCAACGUCUGAUAA**CGGUAAAAAAAACAAAAAACAAAACGGCUAUUAUGCGUUACCGGCGAGACGCUACGGACUUAAAUAA<br>U**UG**AAUGAAAAUCCGUUGACCUUAAACGGUCGUGUGGGUUCAAGUCCCUCCACCCCCACGCCGGAAACGCAAUAGCCGAAAAACAAAAA<br>ACAAAAAAAACAAAAAAAAAACCAAAAAAACAAAACACA |

| | |
|---|---|
| linear firefly luciferase_3xFLAG_non-optimal | *AGGUAGUAUUCUUCUGGUCCCCACAGACUCAGAGAGAACCCGCCACCGCCACC* **AUGGAAGACGCCAAAAACAUAAAGAAAGGCCCGGC**<br>**GCCAUUCUAUCCGCUGGAAGAUGGAACCGCUGGAGAGCAACUGCAUAAGGCUAUGAAGAGAUACGCCCUGGUUCCUGGAACAAUUGC**<br>**UUUUACAGAUGCACAUAUCGAGGUGGACAUCACUUACGCUGAGUACUUCGAAAUGUCCGUUCGGUUGGCAGAAGCUAUGAAACGAUA**<br>**UGGGCUGAAUACAAAUCACAGAAUCGUCGUAUGCAGUGAAAACUCUCUUCAAUUCUUUAUGCCGGUGUUGGGCGCGUUAUUUAUCGG**<br>**AGUUGCAGUUGCGCCCGCGAACGACAUUUAUAAUGAACGUGAAUUGCUCAACAGUAUGGGCAUUUCGCAGCCUACCGUGGUGUUCGU**<br>**UUCCAAAAAGGGGUUGCAAAAAAUUUUGAACGUGCAAAAAAAGCUCCCAAUCAUCCAAAAAAUUAUUAUCAUGGAUUCUAAAACGGAU**<br>**UACCAGGGAUUUCAGUCGAUGUACACGUUCGUCACAUCUCAUCUACCUCCCGGUUUUAAUGAAUACGAUUUUGUGCCAGAGUCCUUC**<br>**GAUAGGGACAAGACAAUUGCACUGAUCAUGAACUCCUCUGGAUCUACUGGUCUGCCUAAAGGUGUCGCUCUGCCUCAUAGAACUGCC**<br>**UGCGUGAGAUUCUCGCAUGCCAGAGAUCCUAUUUUUGGCAAUCAAAUCAUUCCGGAUACUGCGAUUUUAAGUGUUGUUCCAUUCCAU**<br>**CACGGUUUUGGAAUGUUUACUACACUCGGAUAUUUGAUAUGUGGAUUUCGAGUCGUCUUAAUGUAUAGAUUUGAAGAAGAGCUGUUU**<br>**CUGAGGAGCCUUCAGGAUUACAAGAUUCAAAGUGCGCUGCUGGUGCCAACCCUAUUCUCCUUCUUCGCCAAAAGCACUCUGAUUGAC**<br>**AAAUACGAUUUAUCUAAUUUACACGAAAUUGCUUCUGGUGGCGCUCCCCUCUCUAAGGAAGUCGGGGAAGCGGUUGCCAAGAGGUUC**<br>**CAUCUGCCAGGUAUCAGGCAAGGAUAUGGGCUCACUGAGACUACAUCAGCUAUUCUGAUUACACCCGAGGGGGAUGAUAAACCGGGC**<br>**GCGGUCGGUAAAGUUGUUCCAUUUUUUGAAGCGAAGGUUGUGGAUCUGGAUACCGGGAAAACGCUGGGCGUUAAUCAAAGAGGCGAA**<br>**CUGUGUGUGAGAGGUCCUAUGAUUAUGUCCGGUUAUGUAAACAAUCCGGAAGCGACCAACGCCUUGAUUGACAAGGAUGGAUGGCUA**<br>**CAUUCUGGAGACAUAGCUUACUGGGACGAAGACGAACACUUCUUCAUCGUUGACCGCCUGAAGUCUCUGAUUAAGUACAAAGGCUAU**<br>**CAGGUGGCUCCCGCUGAAUUGGAAUCCAUCUUGCUCCAACACCCCAACAUCUUCGACGCAGGUGUCGCAGGUCUUCCCGACGAUGAC**<br>**GCCGGUGAACUUCCCGCCGCCGUUGUUGUUUUGGAGCACGGAAAGACGAUGACGGAAAAAGAGAUCGUGGAUUACGUCGCCAGUCAA**<br>**GUAACAACCGCGAAAAAGUUGCGCGGAGGAGUUGUGUUUGUGGACGAAGUACCGAAAGGUCUUACCGGAAAACUCGACGCAAGAAAA**<br>**AUCAGAGAGAUCCUCAUAAAGGCCAAGAAGGGCGGAAAGAUCGCCGUGGACUACAAAGACCAUGACGGUGAUUAUAAAGAUCAUGAU**<br>**AUCGAUUACAAGGAUGACGAUGACAAGUGAUAA***CUCGAGCUGGUACUGCAUGCACGCAAUGCUAGCUGCCCCUUUCCCGUCCUGGGU*<br>*ACCCCGAGUCUCCCCCGACCUCGGGUCCCAGGUAUGCUCCCACCUCCACCUGCCCCACUCACCACCU* AAAAAAAAAAAAAAAAAAAAAAA<br>AAAAAAAAAAAAAAAAAAAAAAAAAAAAAAAAAAAAAAAAAAAAAAAAAAAAAAAAAAAAAAAAAAAA |

| | |
|---|---|
| linear firefly luciferase_3xFLAG_optimal | *AGGUAGUAUUCUUCUGGUCCCCACAGACUCAGAGAGAACCCGCCACCGCCACC* **AUGGAGGACGCCAAGAACAUCAAGAAGGGCCCCGC CCCAUUCUACCCCCUGGAGGACGGCACCGCCGGCGAGCAGCUGCACAAGGCCAUGAAGCGGUACGCCCUGGUGCCCGGCACCAUCGC CUUCACCGACGCUCACAUCGAGGUGGACAUUACCUACGCCGAGUACUUCGAAAUGAGCGUGCGGCUGGCCGAGGCCAUGAAGCGGUA CGGCCUGAACACCAACCAUCGGAUCGUGGUGUGCAGCGAGAACUCCCUGCAGUUCUUCAUGCCAGUGCUGGGCGCCCUGUUCAUCGG CGUGGCCGUGGCACCCGCCAACGACAUCUACAACGAGCGGGAGCUGCUGAACAGCAUGGGCAUCUCCCAGCCCACCGUGGUGUUCGU GAGCAAGAAGGGCCUGCAGAAGAUCCUGAACGUGCAGAAGAAACUGCCCAUCAUCCAGAAGAUCAUCAUCAUGGACAGCAAGACAGA CUACCAAGGCUUCCAGAGCAUGUACACCUUCGUGACCAGCCACCUGCCCCCUGGCUUCAACGAGUACGACUUCGUGCCAGAGAGCUU CGACAGAGAUAAGACCAUCGCUCUGAUCAUGAACAGCUCCGGCAGCACCGGCCUGCCAAAGGGCGUGGCCCUGCCCCACAGAACCGC CUGCGUGCGGUUCAGCCACGCCCGGGACCCAAUCUUCGGCAACCAGAUCAUCCCCGACACCGCUAUCCUGAGCGUGGUGCCAUUCCA UCACGGCUUCGGCAUGUUCACCACACUGGGCUACCUGAUCUGCGGCUUCCGGGUGGUGCUGAUGUACAGAUUCGAAGAGGAGCUGUU CCUGCGGAGCCUGCAGGACUACAAGAUCCAGAGCGCCCUGCUGGUGCCCACCCUGUUCAGCUUCUUUGCCAAGAGCACCCUGAUCGA CAAGUACGACCUGAGCAACCUGCACGAGAUCGCCAGCGGCGGAGCCCCCCUGAGCAAGGAGGUGGGCGAGGCCGUGGCUAAGAGAU UCCACCUGCCCGGCAUCCGGCAGGGCUACGGCCUGACCGAGACAACCAGCGCCAUCCUGAUCACCCCCGAGGGCGACGACAAGCCAG GCGCCGUGGGCAAGGUGGUGCCCUUCUUCGAGGCCAAGGUGGUGGACCUGGACACCGGCAAGACACUGGGCGUGAAUCAGCGGGGC GAGCUGUGCGUGCGGGGCCCCAUGAUCAUGAGCGGCUACGUGAAUAACCCAGAGGCCACCAACGCCCUGAUCGACAAGGACGGCUGG CUGCACAGCGGCGACAUCGCCUACUGGGACGAGGACGAGCACUUCUUCAUUGUGGACCGGCUGAAGUCCCUGAUUAAGUACAAGGGC UACCAGGUCGCUCCCGCCGAGCUGGAGAGCAUCCUGCUGCAGCACCCCAACAUCUUCGACGCCGGCGUGGCCGGCCUGCCCGACGAU GACGCCGGCGAGCUCCCAGCUGCCGUGGUCGUGCUGGAGCACGGCAAGACCAUGACCGAGAAGGAGAUCGUGGACUACGUGGCUAG CCAGGUGACCACAGCCAAGAAGCUCCGGGGCGGCGUGGUCUUCGUGGACGAGGUGCCCAAGGGCCUGACCGGCAAGCUGGACGCCC GGAAGAUCCGGGAGAUCCUGAUCAAGGCCAAGAAGGGCGGCAAGAUCGCUGUGGACUACAAGGACCACGACGGCGAUUAUAAGGACC ACGACAUCGACUACAAGGACGAUGACGACAAGUGAUAA***CUCGAGCUGGUACUGCAUGCACGCAAUGCUAGCUGCCCCUUUCCCGUCC UGGGUACCCCGAGUCUCCCCCGACCUCGGGUCCCAGGUAUGCUCCCACCUCCACCUGCCCCACUCACCACCU* AAAAAAAAAAAAAAAAA AAAAAAAAAAAAAAAAAAAAAAAAAAAAAAAAAAAAAAAAAAAAAAAAAAAAAAAAAAAAAAAAAAAAAAAAAAA |

| | |
|---|---|
| linear firefly luciferase_3xFLAG_non-optimal+CL1/PEST | *AGGUAGUAUUCUUCUGGUCCCCACAGACUCAGAGAGAACCCGCCACCGCCACC* **AUGGAAGACGCCAAAAACAUAAAGAAAGGCCCGGC GCCAUUCUAUCCGCUGGAAGAUGGAACCGCUGGAGAGCAACUGCAUAAGGCUAUGAAGAGAUACGCCCUGGUUCCUGGAACAAUUGC UUUUACAGAUGCACAUAUCGAGGUGGACAUCACUUACGCUGAGUACUUCGAAAUGUCCGUUCGGUUGGCAGAAGCUAUGAAACGAUA UGGGCUGAAUACAAAUCACAGAAUCGUCGUAUGCAGUGAAAACUCUCUUCAAUUCUUUAUGCCGGUGUUGGGCGCGUUAUUUAUCGG AGUUGCAGUUGCGCCCGCGAACGACAUUUAUAAUGAACGUGAAUUGCUCAACAGUAUGGGCAUUUCGCAGCCUACCGUGGUGUUCGU UUCCAAAAAGGGGUUGCAAAAAAUUUUGAACGUGCAAAAAAAGCUCCCAAUCAUCCAAAAAAUUAUUAUCAUGGAUUCUAAAACGGAU UACCAGGGAUUUCAGUCGAUGUACACGUUCGUCACAUCUCAUCUACCUCCCGGUUUUAAUGAAUACGAUUUUGUGCCAGAGUCCUUC GAUAGGGACAAGACAAUUGCACUGAUCAUGAACUCCUCUGGAUCUACUGGUCUGCCUAAAGGUGUCGCUCUGCCUCAUAGAACUGCC UGCGUGAGAUUCUCGCAUGCCAGAGAUCCUAUUUUUGGCAAUCAAAUCAUUCCGGAUACUGCGAUUUUAAGUGUUGUUCCAUUCCAU CACGGUUUUGGAAUGUUUACUACACUCGGAUAUUUGAUAUGUGGAUUUCGAGUCGUCUUAAUGUAUAGAUUUGAAGAAGAGCUGUUU CUGAGGAGCCUUCAGGAUUACAAGAUUCAAAGUGCGCUGCUGGUGCCAACCCUAUUCUCCUUCUUCGCCAAAAGCACUCUGAUUGAC AAAUACGAUUUAUCUAAUUUACACGAAAUUGCUUCUGGUGGCGCUCCCCUCUCUAAGGAAGUCGGGGAAGCGGUUGCCAAGAGGUUC CAUCUGCCAGGUAUCAGGCAAGGAUAUGGGCUCACUGAGACUACAUCAGCUAUUCUGAUUACACCCGAGGGGGAUGAUAAACCGGGC GCGGUCGGUAAAGUUGUUCCAUUUUUUGAAGCGAAGGUUGUGGAUCUGGAUACCGGGAAAACGCUGGGCGUUAAUCAAAGAGGCGAA CUGUGUGUGAGAGGUCCUAUGAUUAUGUCCGGUUAUGUAAACAAUCCGGAAGCGACCAACGCCUUGAUUGACAAGGAUGGAUGGCUA CAUUCUGGAGACAUAGCUUACUGGGACGAAGACGAACACUUCUUCAUCGUUGACCGCCUGAAGUCUCUGAUUAAGUACAAAGGCUAU CAGGUGGCUCCCGCUGAAUUGGAAUCCAUCUUGCUCCAACACCCCAACAUCUUCGACGCAGGUGUCGCAGGUCUUCCCGACGAUGAC GCCGGUGAACUUCCCGCCGCCGUUGUUGUUUUGGAGCACGGAAAGACGAUGACGGAAAAAGAGAUCGUGGAUUACGUCGCCAGUCAA GUAACAACCGCGAAAAAGUUGCGCGGAGGAGUUGUGUUUGUGGACGAAGUACCGAAAGGUCUUACCGGAAAACUCGACGCAAGAAAA AUCAGAGAGAUCCUCAUAAAGGCCAAGAAGGGCGGAAAGAUCGCCGUGGACUACAAAGACCAUGACGGUGAUUAUAAAGAUCAUGAU AUCGAUUACAAGGAUGACGAUGACAAGAAUUCUGCUUGCAAGAACUGGUUCAGUAGCUUAAGCCACUUUGUGAUCCACCUUAACAGCC ACGGCUUUCCGCCUGAGGUUGAGGAACAGGCCGCCGGUACAUUGCCUAUGUCCUGCGCACAAGAAAGCGGUAUGGACCGGCACCCAG CCGCUUGUGCUUCAGCUCGCAUCAACGUCUGAUAA** *CUCGAGCUGGUACUGCAUGCACGCAAUGCUAGCUGCCCCUUUCCCGUCCUGG GUACCCCGAGUCUCCCCCGACCUCGGGUCCCAGGUAUGCUCCCACCUCCACCUGCCCCACUCACCACCU* AAAAAAAAAAAAAAAAAAAAA AAAAAAAAAAAAAAAAAAAAAAAAAAAAAAAAAAAAAAAAAAAAAAAAAAAAAAAAAAAAAAAAAAAAAA |

| | |
|---|---|
| linear firefly luciferase_3xFLAG_optimal+CL1/PEST | *AGGUAGUAUUCUUCUGGUCCCCACAGACUCAGAGAGAACCCGCCACCGCCACC* **AUGGAGGACGCCAAGAACAUCAAGAAGGGCCCCGC<br>CCCAUUCUACCCCCUGGAGGACGGCACCGCCGGCGAGCAGCUGCACAAGGCCAUGAAGCGGUACGCCCUGGUGCCCGGCACCAUCGC<br>CUUCACCGACGCUCACAUCGAGGUGGACAUUACCUACGCCGAGUACUUCGAAAUGAGCGUGCGGCUGGCCGAGGCCAUGAAGCGGUA<br>CGGCCUGAACACCAACCAUCGGAUCGUGGUGUGCAGCGAGAACUCCCUGCAGUUCUUCAUGCCAGUGCUGGGCGCCCUGUUCAUCGG<br>CGUGGCCGUGGCACCCGCCAACGACAUCUACAACGAGCGGGAGCUGCUGAACAGCAUGGGCAUCUCCCAGCCCACCGUGGUGUUCGU<br>GAGCAAGAAGGGCCUGCAGAAGAUCCUGAACGUGCAGAAGAAACUGCCCAUCAUCCAGAAGAUCAUCAUCAUGGACAGCAAGACAGA<br>CUACCAAGGCUUCCAGAGCAUGUACACCUUCGUGACCAGCCACCUGCCCCCUGGCUUCAACGAGUACGACUUCGUGCCAGAGAGCUU<br>CGACAGAGAUAAGACCAUCGCUCUGAUCAUGAACAGCUCCGGCAGCACCGGCCUGCCAAAGGGCGUGGCCCUGCCCCACAGAACCGC<br>CUGCGUGCGGUUCAGCCACGCCCGGGACCCAAUCUUCGGCAACCAGAUCAUCCCCGACACCGCUAUCCUGAGCGUGGUGCCAUUCCA<br>UCACGGCUUCGGCAUGUUCACCACACUGGGCUACCUGAUCUGCGGCUUCCGGGUGGUGCUGAUGUACAGAUUCGAAGAGGAGCUGUU<br>CCUGCGGAGCCUGCAGGACUACAAGAUCCAGAGCGCCCUGCUGGUGCCCACCCUGUUCAGCUUCUUUGCCAAGAGCACCCUGAUCGA<br>CAAGUACGACCUGAGCAACCUGCACGAGAUCGCCAGCGGCGGAGCCCCCCUGAGCAAGGAGGUGGGCGAGGCCGUGGCUAAGAGAU<br>UCCACCUGCCCGGCAUCCGGCAGGGCUACGGCCUGACCGAGACAACCAGCGCCAUCCUGAUCACCCCCGAGGGCGACGACAAGCCAG<br>GCGCCGUGGGCAAGGUGGUGCCCUUCUUCGAGGCCAAGGUGGUGGACCUGGACACCGGCAAGACACUGGGCGUGAAUCAGCGGGGC<br>GAGCUGUGCGUGCGGGGCCCCAUGAUCAUGAGCGGCUACGUGAAUAACCCAGAGGCCACCAACGCCCUGAUCGACAAGGACGGCUGG<br>CUGCACAGCGGCGACAUCGCCUACUGGGACGAGGACGAGCACUUCUUCAUUGUGGACCGGCUGAAGUCCCUGAUUAAGUACAAGGGC<br>UACCAGGUCGCUCCCGCCGAGCUGGAGAGCAUCCUGCUGCAGCACCCCAACAUCUUCGACGCCGGCGUGGCCGGCCUGCCCGACGAU<br>GACGCCGGCGAGCUCCCAGCUGCCGUGGUCGUGCUGGAGCACGGCAAGACCAUGACCGAGAAGGAGAUCGUGGACUACGUGGCUAG<br>CCAGGUGACCACAGCCAAGAAGCUCCGGGGCGGCGUGGUCUUCGUGGACGAGGUGCCCAAGGGCCUGACCGGCAAGCUGGACGCCC<br>GGAAGAUCCGGGAGAUCCUGAUCAAGGCCAAGAAGGGCGGCAAGAUCGCUGUGGACUACAAGGACCACGACGGCGAUUAUAAGGACC<br>ACGACAUCGACUACAAGGACGAUGACGACAAGAACAGCGCUUGCAAGAACUGGUUCAGCAGCCUCAGCCACUUCGUGAUCCACCUGA<br>ACAGCCACGGCUUCCCCCCCGAGGUGGAGGAGCAGGCCGCCGGAACCCUGCCCAUGUCCUGCGCCCAGGAGAGCGGGAUGGACCGG<br>CACCCAGCCGCUUGCGCCAGCGCCCGGAUCAACGUCUGAUAA** *CUCGAGCUGGUACUGCAUGCACGCAAUGCUAGCUGCCCCUUUCCC<br>GUCCUGGGUACCCCGAGUCUCCCCCGACCUCGGGUCCCAGGUAUGCUCCCACCUCCACCUGCCCCACUCACCACCU* AAAAAAAAAAAAA<br>AAAAAAAAAAAAAAAAAAAAAAAAAAAAAAAAAAAAAAAAAAAAAAAAAAAAAAAAAAAAAAAAAAAAAAAAAAAAAAAAAAAAAAAAAAAAAAAAA |

| | |
|---|---|
| circular firefly<br>luciferase_3xFLAG<br>_non-optimal | UUAAAACAGCCUGUGGGUUGAUCCCACCCACAGGCCCAUUGGGCGCUAGCACUCUGGUAUCACGGUACCUUUGUGCGCCUGUUUUAUA<br>CCCCCUCCCCCAACUGUAACUUAGAAGUAACACACACCGAUCAACAGUCAGCGUGGCACACCAGCCACGUUUUGAUCAAGCACUUCUGU<br>UACCCCGGACUGAGUAUCAAUAGACUGCUCACGCGGUUGAAGGAGAAAGCGUUCGUUAUCCGGCCAACUACUUCGAAAAACCUAGUAAC<br>ACCGUGGAAGUUGCAGAGUGUUUCGCUCAGCACUACCCCAGUGUAGAUCAGGUCGAUGAGUCACCGCAUUCCCCACGGGCGACCGUGG<br>CGGUGGCUGCGUUGGCGGCCUGCCCAUGGGGAAACCCAUGGGACGCUCUAAUACAGACAUGGUGCGAAGAGUCUAUUGAGCUAGUUG<br>GUAGUCCUCCGGCCCCUGAAUGCGGCUAAUCCUAACUGCGGAGCACACACCCUCAAGCCAGAGGGCAGUGUGUCGUAACGGGCAACUC<br>UGCAGCGGAACCGACUACUUUGGGUGUCCGUGUUUCAUUUUAUUCCUAUACUGGCUGCUUAUGGUGACAAUUGAGAGAUCGUUACCAU<br>AUAGCUAUUGGAUUGGCCAUCCGGUGACUAAUAGAGCUAUUAUAUAUCCCUUUGUUGGGUUUAUACCACUUAGCUUGAAAGAGGUUAAA<br>ACAUUACAAUUCAUUGUUAAGUUGAAUACAGCAAA**AUGGAAGACGCCAAAAACAUAAAGAAAGGCCCGGCGCCAUUCUAUCCGCUGGAA**<br>**GAUGGAACCGCUGGAGAGCAACUGCAUAAGGCUAUGAAGAGAUACGCCCUGGUUCCUGGAACAAUUGCUUUUACAGAUGCACAUAUC**<br>**GAGGUGGACAUCACUUACGCUGAGUACUUCGAAAUGUCCGUUCGGUUGGCAGAAGCUAUGAAACGAUAUGGGCUGAAUACAAAUCAC**<br>**AGAAUCGUCGUAUGCAGUGAAAACUCUCUUCAAUUCUUUAUGCCGGUGUUGGGCGCGUUAUUUAUCGGAGUUGCAGUUGCGCCCGCG**<br>**AACGACAUUUAUAAUGAACGUGAAUUGCUCAACAGUAUGGGCAUUUCGCAGCCUACCGUGGUGUUCGUUUCCAAAAAGGGGUUGCAA**<br>**AAAAUUUUGAACGUGCAAAAAAAGCUCCCAAUCAUCCAAAAAAUUAUUAUCAUGGAUUCUAAAACGGAUUACCAGGGAUUUCAGUCGA**<br>**UGUACACGUUCGUCACAUCUCAUCUACCUCCCGGUUUUAAUGAAUACGAUUUUGUGCCAGAGUCCUUCGAUAGGGACAAGACAAUUGC**<br>**ACUGAUCAUGAACUCCUCUGGAUCUACUGGUCUGCCUAAAGGUGUCGCUCUGCCUCAUAGAACUGCCUGCGUGAGAUUCUCGCAUGC**<br>**CAGAGAUCCUAUUUUUGGCAAUCAAAUCAUUCCGGAUACUGCGAUUUUAAGUGUUGUUCCAUUCCAUCACGGUUUUGGAAUGUUUACU**<br>**ACACUCGGAUAUUUGAUAUGUGGAUUUCGAGUCGUCUUAAUGUAUAGAUUUGAAGAAGAGCUGUUUCUGAGGAGCCUUCAGGAUUAC**<br>**AAGAUUCAAAGUGCGCUGCUGGUGCCAACCCUAUUCUCCUUCUUCGCCAAAAGCACUCUGAUUGACAAAUACGAUUUAUCUAAUUUAC**<br>**ACGAAAUUGCUUCUGGUGGCGCUCCCCUCUCUAAGGAAGUCGGGGAAGCGGUUGCCAAGAGGUUCCAUCUGCCAGGUAUCAGGCAAG**<br>**GAUAUGGGCUCACUGAGACUACAUCAGCUAUUCUGAUUACACCCGAGGGGGAUGAUAAACCGGGCGCGGUCGGUAAAGUUGUUCCAU**<br>**UUUUUGAAGCGAAGGUUGUGGAUCUGGAUACCGGGAAAACGCUGGGCGUUAAUCAAAGAGGCGAACUGUGUGUGAGAGGUCCUAUGA**<br>**UUAUGUCCGGUUAUGUAAACAAUCCGGAAGCGACCAACGCCUUGAUUGACAAGGAUGGAUGGCUACAUUCUGGAGACAUAGCUUACU**<br>**GGGACGAAGACGAACACUUCUUCAUCGUUGACCGCCUGAAGUCUCUGAUUAAGUACAAAGGCUAUCAGGUGGCUCCCGCUGAAUUGG**<br>**AAUCCAUCUUGCUCCAACACCCCAACAUCUUCGACGCAGGUGUCGCAGGUCUUCCCGACGAUGACGCCGGUGAACUUCCCGCCGCCG**<br>**UUGUUGUUUUGGAGCACGGAAAGACGAUGACGGAAAAAGAGAUCGUGGAUUACGUCGCCAGUCAAGUAACAACCGCGAAAAAGUUGC**<br>**GCGGAGGAGUUGUGUUUGUGGACGAAGUACCGAAAGGUCUUACCGGAAAACUCGACGCAAGAAAAAUCAGAGAGAUCCUCAUAAAGG**<br>**CCAAGAAGGGCGGAAAGAUCGCCGUGGACUACAAAGACCAUGACGGUGAUUAUAAAGAUCAUGAUAUCGAUUACAAGGAUGACGAUG**<br>**ACAAGUGAUAA**CGGUAAAAAAAACAAAAAACAAAACGGCUAUUAUGCGUUACCGGCGAGACGCUACGGACUUAAAUAAU**UG**AAUGAAAAU<br>CCGUUGACCUUAAACGGUCGUGUGGGUUCAAGUCCCUCCACCCCCACGCCGGAAACGCAAUAGCCGAAAAACAAAAAACAAAAAAAACAA<br>AAAAAAAACCAAAAAAACAAAACACA |

| | |
|---|---|
| circular firefly<br>luciferase_3xFLAG<br>_optimal | UUAAAACAGCCUGUGGGUUGAUCCCACCCACAGGCCCAUUGGGCGCUAGCACUCUGGUAUCACGGUACCUUUGUGCGCCUGUUUUAUA<br>CCCCCUCCCCCAACUGUAACUUAGAAGUAACACACACCGAUCAACAGUCAGCGUGGCACACCAGCCACGUUUUGAUCAAGCACUUCUGU<br>UACCCCGGACUGAGUAUCAAUAGACUGCUCACGCGGUUGAAGGAGAAAGCGUUCGUUAUCCGGCCAACUACUUCGAAAAACCUAGUAAC<br>ACCGUGGAAGUUGCAGAGUGUUUCGCUCAGCACUACCCCAGUGUAGAUCAGGUCGAUGAGUCACCGCAUUCCCCACGGGCGACCGUGG<br>CGGUGGCUGCGUUGGCGGCCUGCCCAUGGGGAAACCCAUGGGACGCUCUAAUACAGACAUGGUGCGAAGAGUCUAUUGAGCUAGUUG<br>GUAGUCCUCCGGCCCCUGAAUGCGGCUAAUCCUAACUGCGGAGCACACACCCUCAAGCCAGAGGGCAGUGUGUCGUAACGGGCAACUC<br>UGCAGCGGAACCGACUACUUUGGGUGUCCGUGUUUCAUUUUAUUCCUAUACUGGCUGCUUAUGGUGACAAUUGAGAGAUCGUUACCAU<br>AUAGCUAUUGGAUUGGCCAUCCGGUGACUAAUAGAGCUAUUAUAUCCCUUUGUUGGGUUUAUACCACUUAGCUUGAAAGAGGUUAAA<br>ACAUUACAAUUCAUUGUUAAGUUGAAUACAGCAAA**AUGGAGGACGCCAAGAACAUCAAGAAGGGCCCCGCCCCAUUCUACCCCCUGGA**<br>**GGACGGCACCGCCGGCGAGCAGCUGCACAAGGCCAUGAAGCGGUACGCCCUGGUGCCCGGCACCAUCGCCUUCACCGACGCUCACAU**<br>**CGAGGUGGACAUUACCUACGCCGAGUACUUCGAAAUGAGCGUGCGGCUGGCCGAGGCCAUGAAGCGGUACGGCCUGAACACCAACCA**<br>**UCGGAUCGUGGUGUGCAGCGAGAACUCCCUGCAGUUCUUCAUGCCAGUGCUGGGCGCCCUGUUCAUCGGCGUGGCCGUGGCACCCG**<br>**CCAACGACAUCUACAACGAGCGGGAGCUGCUGAACAGCAUGGGCAUCUCCCAGCCCACCGUGGUGUUCGUGAGCAAGAAGGGCCUGC**<br>**AGAAGAUCCUGAACGUGCAGAAGAAACUGCCCAUCAUCCAGAAGAUCAUCAUCAUGGACAGCAAGACAGACUACCAAGGCUUCCAGAG**<br>**CAUGUACACCUUCGUGACCAGCCACCUGCCCCCUGGCUUCAACGAGUACGACUUCGUGCCAGAGAGCUUCGACAGAGAUAAGACCAU**<br>**CGCUCUGAUCAUGAACAGCUCCGGCAGCACCGGCCUGCCAAAGGGCGUGGCCCUGCCCCACAGAACCGCCUGCGUGCGGUUCAGCCA**<br>**CGCCCGGGACCCAAUCUUCGGCAACCAGAUCAUCCCCGACACCGCUAUCCUGAGCGUGGUGCCAUUCCAUCACGGCUUCGGCAUGUU**<br>**CACCACACUGGGCUACCUGAUCUGCGGCUUCCGGGUGGUGCUGAUGUACAGAUUCGAAGAGGAGCUGUUCCUGCGGAGCCUGCAGGA**<br>**CUACAAGAUCCAGAGCGCCCUGCUGGUGCCCACCCUGUUCAGCUUCUUUGCCAAGAGCACCCUGAUCGACAAGUACGACCUGAGCAA**<br>**CCUGCACGAGAUCGCCAGCGGCGGAGCCCCCCUGAGCAAGGAGGUGGGCGAGGCCGUGGCUAAGAGAUUCCACCUGCCCGGCAUCC**<br>**GGCAGGGCUACGGCCUGACCGAGACAACCAGCGCCAUCCUGAUCACCCCCGAGGGCGACGACAAGCCAGGCGCCGUGGGCAAGGUG**<br>**GUGCCCUUCUUCGAGGCCAAGGUGGUGGACCUGGACACCGGCAAGACACUGGGCGUGAAUCAGCGGGGCGAGCUGUGCGUGCGGGG**<br>**CCCCAUGAUCAUGAGCGGCUACGUGAAUAACCCAGAGGCCACCAACGCCCUGAUCGACAAGGACGGCUGGCUGCACAGCGGCGACAU**<br>**CGCCUACUGGGACGAGGACGAGCACUUCUUCAUUGUGGACCGGCUGAAGUCCCUGAUUAAGUACAAGGGCUACCAGGUCGCUCCCGC**<br>**CGAGCUGGAGAGCAUCCUGCUGCAGCACCCCAACAUCUUCGACGCCGGCGUGGCCGGCCUGCCCGACGAUGACGCCGGCGAGCUCCC**<br>**AGCUGCCGUGGUCGUGCUGGAGCACGGCAAGACCAUGACCGAGAAGGAGAUCGUGGACUACGUGGCUAGCCAGGUGACCACAGCCAA**<br>**GAAGCUCCGGGGCGGCGUGGUCUUCGUGGACGAGGUGCCCAAGGGCCUGACCGGCAAGCUGGACGCCCGGAAGAUCCGGGAGAUCC**<br>**UGAUCAAGGCCAAGAAGGGCGGCAAGAUCGCUGUGGACUACAAGGACCACGACGGCGAUUAUAAGGACCACGACAUCGACUACAAGG**<br>**ACGAUGACGACAAGUGAUAA**CGGUAAAAAAAAACAAAAAAACAAAACGGCUAUUAUGCGUUACCGGCGAGACGCUACGGACUUAAAUAAU**U**<br>**G**AAUGAAAAUCCGUUGACCUUAAACGGUCGUGUGGGUUCAAGUCCCUCCACCCCCACGCCGGAAACGCAAUAGCCGAAAAACAAAAAAC<br>AAAAAAAACAAAAAAAAAAACCAAAAAAAACAAAACACA |

| | |
|---|---|
| circular firefly luciferase_3xFLAG _non-optimal+CL1/PEST | UUAAAACAGCCUGUGGGUUGAUCCCACCCACAGGCCCAUUGGGCGCUAGCACUCUGGUAUCACGGUACCUUUGUGCGCCUGUUUUAUA<br>CCCCCUCCCCCAACUGUAACUUAGAAGUAACACACACCGAUCAACAGUCAGCGUGGCACACCAGCCACGUUUUGAUCAAGCACUUCUGU<br>UACCCCGGACUGAGUAUCAAUAGACUGCUCACGCGGUUGAAGGAGAAAGCGUUCGUUAUCCGGCCAACUACUUCGAAAAACCUAGUAAC<br>ACCGUGGAAGUUGCAGAGUGUUUCGCUCAGCACUACCCCAGUGUAGAUCAGGUCGAUGAGUCACCGCAUUCCCCACGGGCGACCGUGG<br>CGGUGGCUGCGUUGGCGGCCUGCCCAUGGGGAAACCCAUGGGACGCUCUAAUACAGACAUGGUGCGAAGAGUCUAUUGAGCUAGUUG<br>GUAGUCCUCCGGCCCCUGAAUGCGGCUAAUCCUAACUGCGGAGCACACACCCUCAAGCCAGAGGGCAGUGUGUCGUAACGGGCAACUC<br>UGCAGCGGAACCGACUACUUUGGGUGUCCGUGUUUCAUUUUAUUCCUAUACUGGCUGCUUAUGGUGACAAUUGAGAGAUCGUUACCAU<br>AUAGCUAUUGGAUUGGCCAUCCGGUGACUAAUAGAGCUAUUAUAUCCCUUUGUUGGGUUUAUACCACUUAGCUUGAAAGAGGUUAAA<br>ACAUUACAAUUCAUUGUUAAGUUGAAUACAGCAAA**AUGGAAGACGCCAAAAACAUAAAGAAAGGCCCGGCGCCAUUCUAUCCGCUGGAA**<br>**GAUGGAACCGCUGGAGAGCAACUGCAUAAGGCUAUGAAGAGAUACGCCCUGGUUCCUGGAACAAUUGCUUUUACAGAUGCACAUAUC**<br>**GAGGUGGACAUCACUUACGCUGAGUACUUCGAAAUGUCCGUUCGGUUGGCAGAAGCUAUGAAACGAUAUGGGCUGAAUACAAAUCAC**<br>**AGAAUCGUCGUAUGCAGUGAAAACUCUCUUCAAUUCUUUAUGCCGGUGUUGGGCGCGUUAUUUAUCGGAGUUGCAGUUGCGCCCGCG**<br>**AACGACAUUUAUAAUGAACGUGAAUUGCUCAACAGUAUGGGCAUUUCGCAGCCUACCGUGGUGUUCGUUUCCAAAAAGGGGUUGCAA**<br>**AAAAUUUUGAACGUGCAAAAAAAGCUCCCAAUCAUCCAAAAAAUUAUUAUCAUGGAUUCUAAAACGGAUUACCAGGGAUUUCAGUCGA**<br>**UGUACACGUUCGUCACAUCUCAUCUACCUCCCGGUUUUAAUGAAUACGAUUUUGUGCCAGAGUCCUUCGAUAGGGACAAGACAAUUGC**<br>**ACUGAUCAUGAACUCCUCUGGAUCUACUGGUCUGCCUAAAGGUGUCGCUCUGCCUCAUAGAACUGCCUGCGUGAGAUUCUCGCAUGC**<br>**CAGAGAUCCUAUUUUUGGCAAUCAAAUCAUUCCGGAUACUGCGAUUUUAAGUGUUGUUCCAUUCCAUCACGGUUUUGGAAUGUUUACU**<br>**ACACUCGGAUAUUUGAUAUGUGGAUUUCGAGUCGUCUUAAUGUAUAGAUUUGAAGAAGAGCUGUUUCUGAGGAGCCUUCAGGAUUAC**<br>**AAGAUUCAAAGUGCGCUGCUGGUGCCAACCCUAUUCUCCUUCUUCGCCAAAAGCACUCUGAUUGACAAAUACGAUUUAUCUAAUUUAC**<br>**ACGAAAUUGCUUCUGGUGGCGCUCCCCUCUCUAAGGAAGUCGGGGAAGCGGUUGCCAAGAGGUUCCAUCUGCCAGGUAUCAGGCAAG**<br>**GAUAUGGGCUCACUGAGACUACAUCAGCUAUUCUGAUUACACCCGAGGGGGAUGAUAAACCGGGCGCGGUCGGUAAAGUUGUUCCAU**<br>**UUUUUGAAGCGAAGGUUGUGGAUCUGGAUACCGGGAAAACGCUGGGCGUUAAUCAAAGAGGCGAACUGUGUGUGAGAGGUCCUAUGA**<br>**UUAUGUCCGGUUAUGUAAACAAUCCGGAAGCGACCAACGCCUUGAUUGACAAGGAUGGAUGGCUACAUUCUGGAGACAUAGCUUACU**<br>**GGGACGAAGACGAACACUUCUUCAUCGUUGACCGCCUGAAGUCUCUGAUUAAGUACAAAGGCUAUCAGGUGGCUCCCGCUGAAUUGG**<br>**AAUCCAUCUUGCUCCAACACCCCAACAUCUUCGACGCAGGUGUCGCAGGUCUUCCCGACGAUGACGCCGGUGAACUUCCCGCCGCCG**<br>**UUGUUGUUUUGGAGCACGGAAAGACGAUGACGGAAAAAGAGAUCGUGGAUUACGUCGCCAGUCAAGUAACAACCGCGAAAAAGUUGC**<br>**GCGGAGGAGUUGUGUUUGUGGACGAAGUACCGAAAGGUCUUACCGGAAAACUCGACGCAAGAAAAAUCAGAGAGAUCCUCAUAAAGG**<br>**CCAAGAAGGGCGGAAAGAUCGCCGUGGACUACAAAGACCAUGACGGUGAUUAUAAAGAUCAUGAUAUCGAUUACAAGGAUGACGAUG**<br>**ACAAGAAUUCUGCUUGCAAGAACUGGUUCAGUAGCUUAAGCCACUUUGUGAUCCACCUUAACAGCCACGGCUUUCCGCCUGAGGUUG**<br>**AGGAACAGGCCGCCGGUACAUUGCCUAUGUCCUGCGCACAAGAAAGCGGUAUGGACCGGCACCCAGCCGCUUGUGCUUCAGCUCGCA**<br>**UCAACGUCUGAUAA**CGGUAAAAAAAACAAAAAACAAAACGGCUAUUAUGCGUUACCGGCGAGACGCUACGGACUUAAAUAAU**UG**AAUGAA<br>AAUCCGUUGACCUUAAACGGUCGUGUGGGUUCAAGUCCCUCCACCCCCACGCCGGAAACGCAAUAGCCGAAAAACAAAAAACAAAAAAAA<br>CAAAAAAAAAACCAAAAAAACAAAACACA |

| | |
|---|---|
| circular firefly luciferase_3xFLAG_optimal+CL1/PEST | UUAAAACAGCCUGUGGGUUGAUCCCACCCACAGGCCCAUUGGGCGCUAGCACUCUGGUAUCACGGUACCUUUGUGCGCCUGUUUUAUA<br>CCCCCUCCCCCAACUGUAACUUAGAAGUAACACACACCGAUCAACAGUCAGCGUGGCACACCAGCCACGUUUUGAUCAAGCACUUCUGU<br>UACCCCGGACUGAGUAUCAAUAGACUGCUCACGCGGUUGAAGGAGAAAGCGUUCGUUAUCCGGCCAACUACUUCGAAAAACCUAGUAAC<br>ACCGUGGAAGUUGCAGAGUGUUUCGCUCAGCACUACCCCAGUGUAGAUCAGGUCGAUGAGUCACCGCAUUCCCCACGGGCGACCGUGG<br>CGGUGGCUGCGUUGGCGGCCUGCCCAUGGGGAAACCCAUGGGACGCUCUAAUACAGACAUGGUGCGAAGAGUCUAUUGAGCUAGUUG<br>GUAGUCCUCCGGCCCCUGAAUGCGGCUAAUCCUAACUGCGGAGCACACACCCUCAAGCCAGAGGGCAGUGUGUCGUAACGGGCAACUC<br>UGCAGCGGAACCGACUACUUUGGGUGUCCGUGUUUCAUUUUAUUCCUAUACUGGCUGCUUAUGGUGACAAUUGAGAGAUCGUUACCAU<br>AUAGCUAUUGGAUUGGCCAUCCGGUGACUAAUAGAGCUAUUAUAUAUCCCUUUGUUGGGUUUAUACCACUUAGCUUGAAAGAGGUUAAA<br>ACAUUACAAUUCAUUGUUAAGUUGAAUACAGCAAA**AUGGAGGACGCCAAGAACAUCAAGAAGGGCCCCGCCCCAUUCUACCCCCUGGA**<br>**GGACGGCACCGCCGGCGAGCAGCUGCACAAGGCCAUGAAGCGGUACGCCCUGGUGCCCGGCACCAUCGCCUUCACCGACGCUCACAU**<br>**CGAGGUGGACAUUACCUACGCCGAGUACUUCGAAAUGAGCGUGCGGCUGGCCGAGGCCAUGAAGCGGUACGGCCUGAACACCAACCA**<br>**UCGGAUCGUGGUGUGCAGCGAGAACUCCCUGCAGUUCUUCAUGCCAGUGCUGGGCGCCCUGUUCAUCGGCGUGGCCGUGGCACCCG**<br>**CCAACGACAUCUACAACGAGCGGGAGCUGCUGAACAGCAUGGGCAUCUCCCAGCCCACCGUGGUGUUCGUGAGCAAGAAGGGCCUGC**<br>**AGAAGAUCCUGAACGUGCAGAAGAAACUGCCCAUCAUCCAGAAGAUCAUCAUCAUGGACAGCAAGACAGACUACCAAGGCUUCCAGAG**<br>**CAUGUACACCUUCGUGACCAGCCACCUGCCCCCUGGCUUCAACGAGUACGACUUCGUGCCAGAGAGCUUCGACAGAGAUAAGACCAU**<br>**CGCUCUGAUCAUGAACAGCUCCGGCAGCACCGGCCUGCCAAAGGGCGUGGCCCUGCCCCACAGAACCGCCUGCGUGCGGUUCAGCCA**<br>**CGCCCGGGACCCAAUCUUCGGCAACCAGAUCAUCCCCGACACCGCUAUCCUGAGCGUGGUGCCAUUCCAUCACGGCUUCGGCAUGUU**<br>**CACCACACUGGGCUACCUGAUCUGCGGCUUCCGGGUGGUGCUGAUGUACAGAUUCGAAGAGGAGCUGUUCCUGCGGAGCCUGCAGGA**<br>**CUACAAGAUCCAGAGCGCCCUGCUGGUGCCCACCCUGUUCAGCUUCUUUGCCAAGAGCACCCUGAUCGACAAGUACGACCUGAGCAA**<br>**CCUGCACGAGAUCGCCAGCGGCGGAGCCCCCCUGAGCAAGGAGGUGGGCGAGGCCGUGGCUAAGAGAUUCCACCUGCCCGGCAUCC**<br>**GGCAGGGCUACGGCCUGACCGAGACAACCAGCGCCAUCCUGAUCACCCCCGAGGGCGACGACAAGCCAGGCGCCGUGGGCAAGGUG**<br>**GUGCCCUUCUUCGAGGCCAAGGUGGUGGACCUGGACACCGGCAAGACACUGGGCGUGAAUCAGCGGGGCGAGCUGUGCGUGCGGGG**<br>**CCCCAUGAUCAUGAGCGGCUACGUGAAUAACCCAGAGGCCACCAACGCCCUGAUCGACAAGGACGGCUGGCUGCACAGCGGCGACAU**<br>**CGCCUACUGGGACGAGGACGAGCACUUCUUCAUUGUGGACCGGCUGAAGUCCCUGAUUAAGUACAAGGGCUACCAGGUCGCUCCCGC**<br>**CGAGCUGGAGAGCAUCCUGCUGCAGCACCCCAACAUCUUCGACGCCGGCGUGGCCGGCCUGCCCGACGAUGACGCCGGCGAGCUCCC**<br>**AGCUGCCGUGGUCGUGCUGGAGCACGGCAAGACCAUGACCGAGAAGGAGAUCGUGGACUACGUGGCUAGCCAGGUGACCACAGCCAA**<br>**GAAGCUCCGGGGCGGCGUGGUCUUCGUGGACGAGGUGCCCAAGGGCCUGACCGGCAAGCUGGACGCCCGGAAGAUCCGGGAGAUCC**<br>**UGAUCAAGGCCAAGAAGGGCGGCAAGAUCGCUGUGGACUACAAGGACCACGACGGCGAUUAUAAGGACCACGACAUCGACUACAAGG**<br>**ACGAUGACGACAAGAACAGCGCUUGCAAGAACUGGUUCAGCAGCCUCAGCCACUUCGUGAUCCACCUGAACAGCCACGGCUUCCCCC**<br>**CCGAGGUGGAGGAGCAGGCCGCCGGAACCCUGCCCAUGUCCUGCGCCCAGGAGAGCGGGAUGGACCGGCACCCAGCCGCUUGCGCC**<br>**AGCGCCCGGAUCAACGUCUGAUAA**CGGUAAAAAAAACAAAAAACAAAACGGCUAUUAUGCGUUACCGGCGAGACGCUACGGACUUAAAU<br>AAUUGAAUGAAAAUCCGUUGACCUUAAACGGUCGUGUGGGUUCAAGUCCCUCCACCCCCACGCCGGAAACGCAAUAGCCGAAAAACAAA<br>AAACAAAAAAAACAAAAAAAAAACCAAAAAAACAAAACACA |

| | |
|---|---|
| linear EGFP_3xFLAG mRNA | *AGGUAGUAUUCUUCUGGUCCCCACAGACUCAGAGAGAACCCGCCACCGCCACC* **AUGGUGAGCAAGGGCGAGGAGCUGUUCACCGGGG**<br>**UGGUGCCCAUCCUGGUCGAGCUGGACGGCGACGUAAACGGCCACAAGUUCAGCGUGUCCGGCGAGGGCGAGGGCGAUGCCACCUAC**<br>**GGCAAGCUGACCCUGAAGUUCAUCUGCACCACCGGCAAGCUGCCCGUGCCCUGGCCCACCCUCGUGACCACCCUGACCUACGGCGUG**<br>**CAGUGCUUCAGCCGCUACCCCGACCACAUGAAGCAGCACGACUUCUUCAAGUCCGCCAUGCCCGAAGGCUACGUCCAGGAGCGCACC**<br>**AUCUUCUUCAAGGACGACGGCAACUACAAGACCCGCGCCGAGGUGAAGUUCGAGGGCGACACCCUGGUGAACCGCAUCGAGCUGAAG**<br>**GGCAUCGACUUCAAGGAGGACGGCAACAUCCUGGGGCACAAGCUGGAGUACAACUACAACAGCCACAACGUCUAUAUCAUGGCCGAC**<br>**AAGCAGAAGAACGGCAUCAAGGUGAACUUCAAGAUCCGCCACAACAUCGAGGACGGCAGCGUGCAGCUCGCCGACCACUACCAGCAG**<br>**AACACCCCCAUUGGCGACGGCCCCGUGCUGCUGCCCGACAACCACUACCUGAGCACCCAGUCCGCCCUGAGCAAAGACCCCAACGAG**<br>**AAGCGCGAUCACAUGGUCCUGCUGGAGUUCGUGACCGCCGCCGGGAUCACUCUCGGCAUGGACGAGCUGUACAAGGACUACAAAGAC**<br>**CAUGACGGUGAUUAUAAAGAUCAUGAUAUCGAUUACAAGGAUGACGAUGACAAGUGAUAA***CUCGAGCUGGUACUGCAUGCACGCAAU*<br>*GCUAGCUGCCCCUUUCCCGUCCUGGGUACCCCGAGUCUCCCCCGACCUCGGGUCCCAGGUAUGCUCCCACCUCCACCUGCCCCACUCA*<br>*CCACCU* AAAAAAAAAAAAAAAAAAAAAAAAAAAAAAAAAAAAAAAAAAAAAAAAAAAAAAAAAAAAAAAAAAAAAAAAAAAAAAAAAAAAAAAAAAAAAA<br>AA |
| representative linear precursor for circRNAs | GGGAGAGGGAGACCCUCGACCGUCGAUUGUCCACUGGUCAACAAUAGAUGACUUACAACUAAUCGGAAGGUGCAGAGACUCGACGGGA<br>GCUACCCUAACGUCAAGACGAGGGUAAAGAGAGAGUCCAAUUCUCAAAGCCAAUAGGCAGUAGCGAAAGCUGCAAGA**G**AAUGAAAAUCC<br>GUUGACCUUAAACGGUCGUGUGGGUUCAAGUCCCUCCACCCCCACGCCGGAAACGCAAUAGCCGAAAAACAAAAAACAAAAAAAACAAAA<br>AAAAAACCAAAAAAACAAAACACAUUAAAACAGCCUGUGGGUUGAUCCCACCCACAGGCCCAUUGGGCGCUAGCACUCUGGUAUCACGGU<br>ACCUUUGUGCGCCUGUUUUAUACCCCCUCCCCCAACUGUAACUUAGAAGUAACACACACCGAUCAACAGUCAGCGUGGCACACCAGCCA<br>CGUUUUGAUCAAGCACUUCUGUUACCCCGGACUGAGUAUCAAUAGACUGCUCACGCGGUUGAAGGAGAAAGCGUUCGUUAUCCGGCCA<br>ACUACUUCGAAAAACCUAGUAACACCGUGGAAGUUGCAGAGUGUUUCGCUCAGCACUACCCCAGUGUAGAUCAGGUCGAUGAGUCACCG<br>CAUUCCCCACGGGCGACCGUGGCGGUGGCUGCGUUGGCGGCCUGCCCAUGGGGAAACCCAUGGGACGCUCUAAUACAGACAUGGUGC<br>GAAGAGUCUAUUGAGCUAGUUGGUAGUCCUCCGGCCCCUGAAUGCGGCUAAUCCUAACUGCGGAGCACACACCCUCAAGCCAGAGGGC<br>AGUGUGUCGUAACGGGCAACUCUGCAGCGGAACCGACUACUUUGGGUGUCCGUGUUUCAUUUUAUUCCUAUACUGGCUGCUUAUGGUG<br>ACAAUUGAGAGAUCGUUACCAUAUAGCUAUUGGAUUGGCCAUCCGGUGACUAAUAGAGCUAUUAUAUAUCCCUUUGUUGGGUUUAUACC<br>ACUUAGCUUGAAAGAGGUUAAAACAUUACAAUUCAUUGUUAAGUUGAAUACAGCAAA **[insert open reading frame beginning with start and codon and ending with stop codon here]**<br>UAAAAAAAACAAAAAACAAAACGGCUAUUAUGCGUUACCGGCGAGACGCUACGGACUUAAAUAAU**U**GAGCCUUAAAGAAGAAAUUCUUUA<br>AGUGGAUGCUCUCAAACUCAGGGAAACCUAAAUCUAGUUAUAGACAAGGCAAUCCUGAGCCAAGCCGAAGUAGUAAUUAGUAAGACCAG<br>UGGACAAUCGACGGAUAACAGCAUAUCUAG |

**Table S5: RNA sequences used in this study.** The RNA sequences for the different reporter RNAs are shown. Please note that all linear mRNAs begin with an 'A' as the first transcribed nucleotide and were in vitro transcribed using NEB's HiScribe T7 + Clean Cap AG kit. Therefore, mRNAs have an m7G cap1 structure. For all RNAs, coding sequences are in **bold** and 3xFLAG and 3xFLAG+hCL1+hPEST degron sequences are in **purple.** Please note that the RNA sequences for FLAG+/- degron tags are different for optimal/non-optimal RNAs. IRES sequences are underlined, UTR sequences are in *italics* , Stop codons are in **red**, the poly(A) tail and other sequences retained in the final circRNAs are shown in standard font. For consistency, final circRNAs are shown with the start of the CVB3 IRES as poition 1, followed by the coding region, and then the other sequences included in the final circRNA. The two bases showing the circularization splice junction are in **blue**. A representative circRNA precursor is shown as a linear RNA with a placeholder for the open reading frame.
