## Supplemental Figures for "Codon optimality modulates cellular stress and innate immune responses triggered by exogenous RNAs"

### Supplementary Figures and legends

**A**

| | | NLuc | RLuc | FLuc |
|---|---|---|---|---|
| **CDS length (nt)** | | 582 | 936 | 1719 |
| **Codon adaptation index (CAI)** | Non-optimal | 0.73 | 0.66 | 0.73 |
| | Optimal | 0.98 | 0.97 | 0.97 |
| **GC content (%)** | Non-optimal | 45 | 37 | 46 |
| | Optimal | 61 | 60 | 62 |
| **U content (%)** | Non-optimal | 26 | 31 | 26 |
| | Optimal | 16 | 15 | 15 |
| **Minimum free energy (MFE, kcal/mol)** | Non-optimal | -246.6 | -295.7 | -563.2 |
| | Optimal | -282.7 | -416.8 | -741.5 |

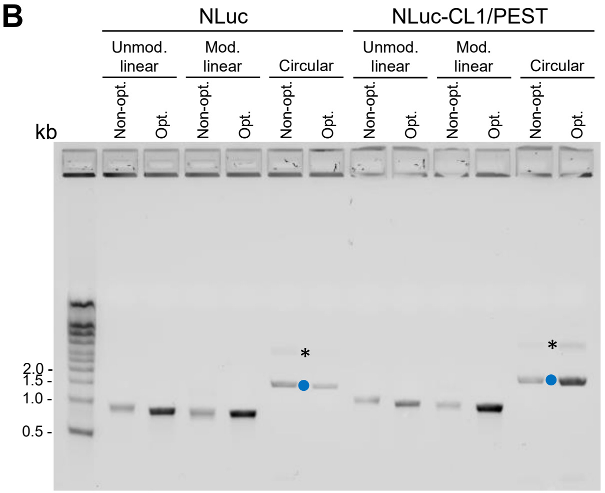


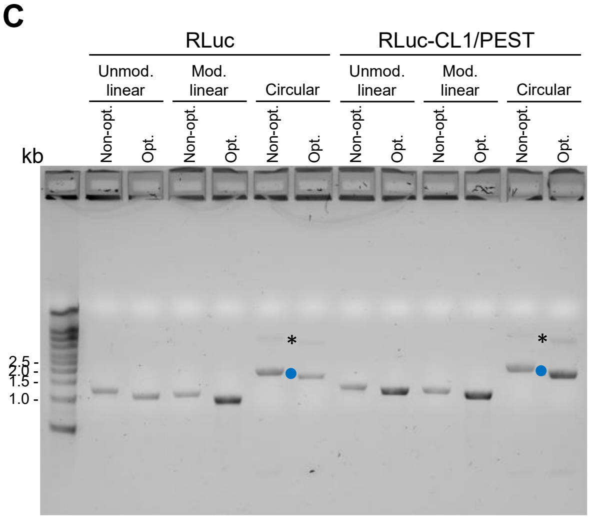


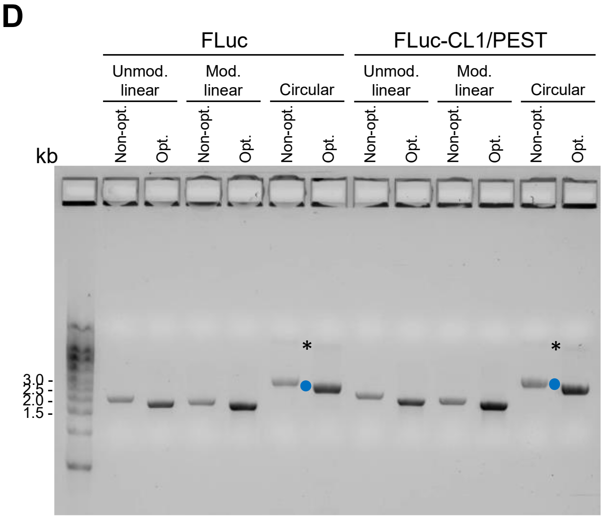


**Figure S1. Coding region comparisons and agarose gels showing the RNA constructs used in this study. (A)** Characteristics of non-optimal and optimal reporter coding sequences used in this study. Codon adaptation index (CAI) scores and minimum free energy (MFE) were calculated as described in methods. **(B-D)** Linear RNAs were synthesized by *in vitro* transcription from a linearized plasmid DNA template and co-transcriptionally capped using a HiScribe® T7 mRNA Kit with CleanCap Reagent AG Kit. For modified linear mRNAs, UTP was replaced by N1-methylpseudo-UTP. circRNA precursors were synthesized by *in vitro* transcription from a linearized plasmid DNA template using a HiScribe® T7 High Yield RNA Synthesis Kit, subjected to the circularization reaction, and then treated with RNase R. Blue circles mark circRNAs and asterisks mark concatenated circRNAs where two copies of the circRNA precursor were joined into a single circRNA.

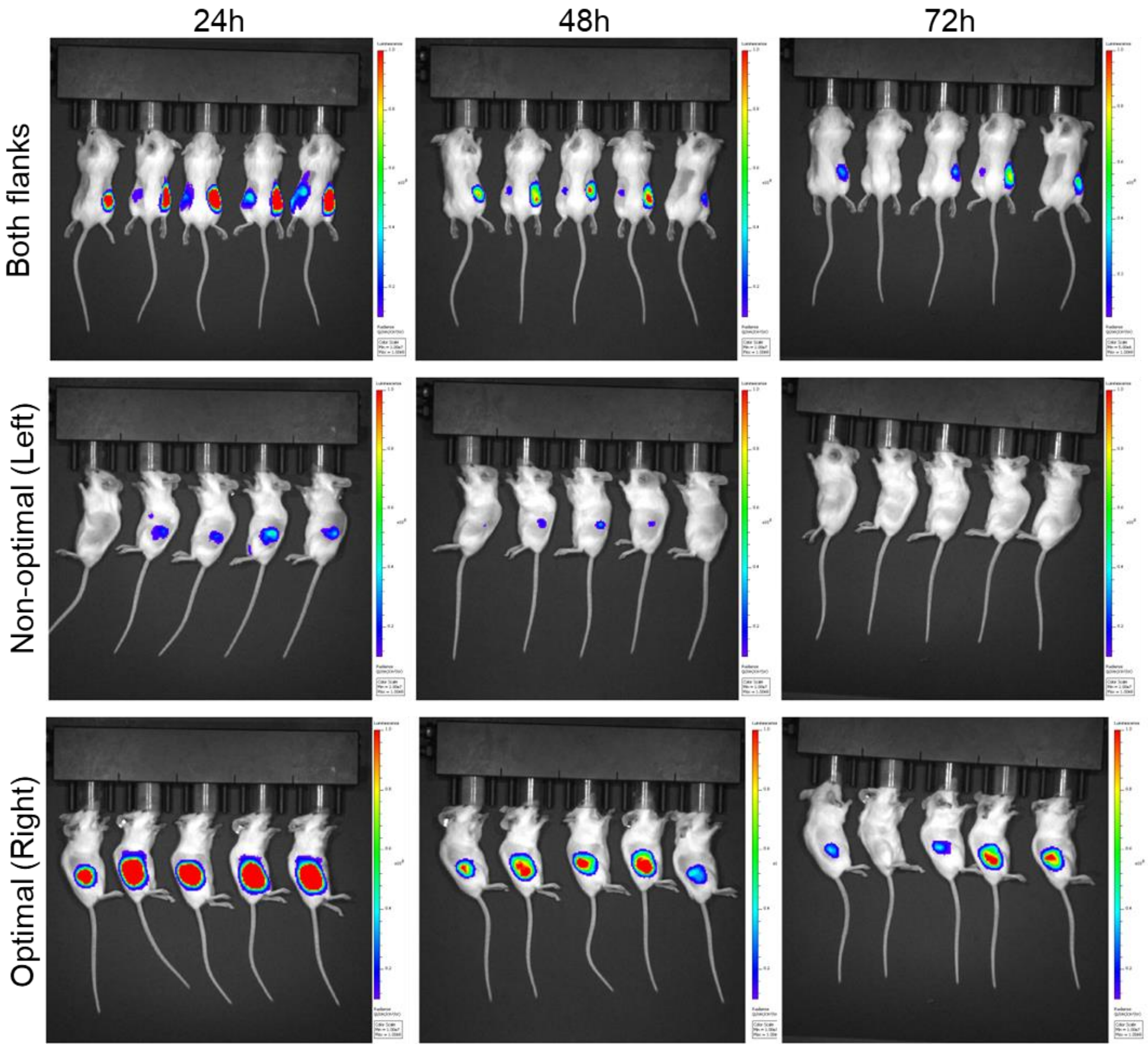


**Figure S2. Highly optimal firefly luciferase (FLuc) mRNAs outperform non-optimal sequences *in vivo*.** Five mice were injected intramuscularly with 5 µg of LNP-encapsulated unmodified FLuc mRNA with a non-optimal coding region (left flank) or our optimal design (right flank). FLuc bioluminescence was imaged daily for 72 h using IVIS. Three views of the mice are shown with (top) an overhead view showing both flanks or views showing only the (middle) left (non-optimal mRNA) side (bottom) or right (optimal mRNA) side of the animal. The scales for all the images range from a minimum value of $10^7$ to a maximum reading of $10^8$. For the continuity of the data set, the top left image (24h overhead image) is the same as in Figure 1A.

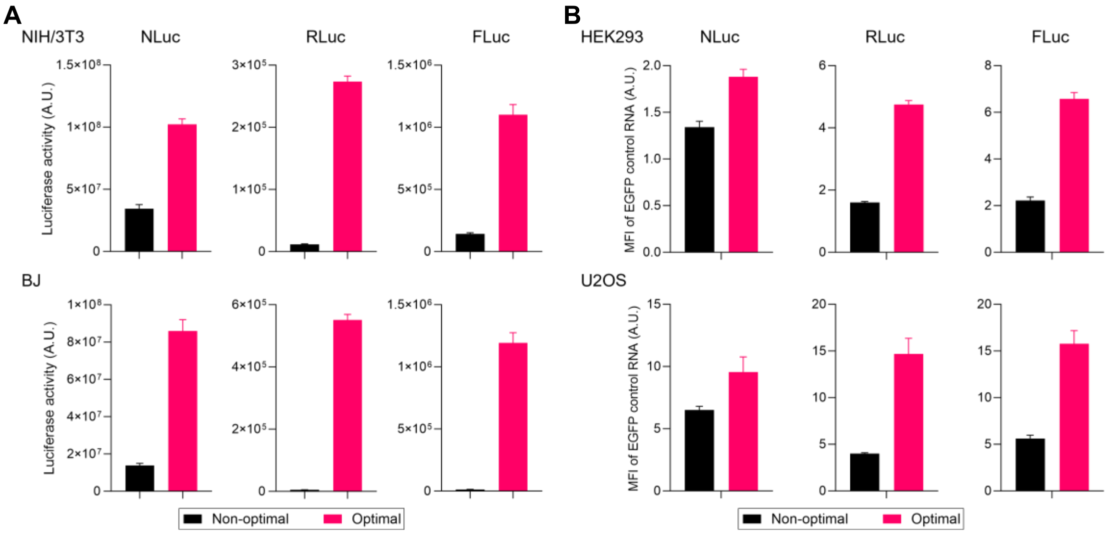


**Figure S3. Codon optimality of test mRNAs affects expression of both test and control mRNAs. (A)** Dual-luciferase assays using lysates from NIH/3T3 or BJ cells harvested 24 h after transfection with mRNAs encoding either optimal (pink) or non-optimal (black) forms of Nanoluciferase (NLuc), *Renilla* luciferase (RLuc), or firefly luciferase (FLuc) reporters along with non-optimal luciferase transfection control RNA. Luciferase activities of the indicated reporter proteins are shown as the mean +Std Dev of four independent experiments. **(B)** mRNAs encoding luciferase proteins with different codon optimality formulas show differences in protein levels of transfection control mRNAs encoding EGFP. Mean green fluorescence intensity of HEK293 or U2OS cells 24 h after co-transfection with mRNAs encoding either non-optimal (black) or optimal (pink) luciferase reporter and an enhanced green fluorescent protein (EGFP) transfection control. Quantitative fluorescence images were taken by an Incucyte S3 and data are shown as the mean signal +Std Dev of four independent replicates with five images taken per replicate.

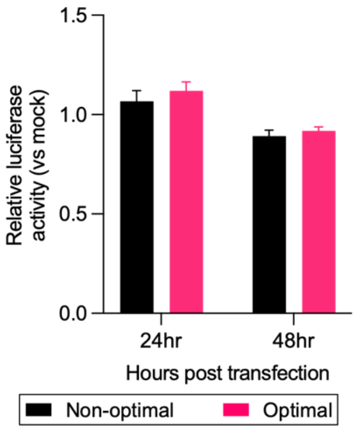


**Figure S4. Codon optimality does not affect cell viability.** Cell viability was determined using CellTiter-Glo® Luminescent Cell Viability Assay. HEK293 cells were transfected with unmodified linear mRNAs expressing non-optimal (black) or optimal (pink) *Renilla* luciferase reporters. Cell viability was assayed at 24 and 48 h after transfection with relative fold change normalized to the viability of control (mock) transfection. Data are shown as the mean of five independent replicates where error bars represent Std Dev.

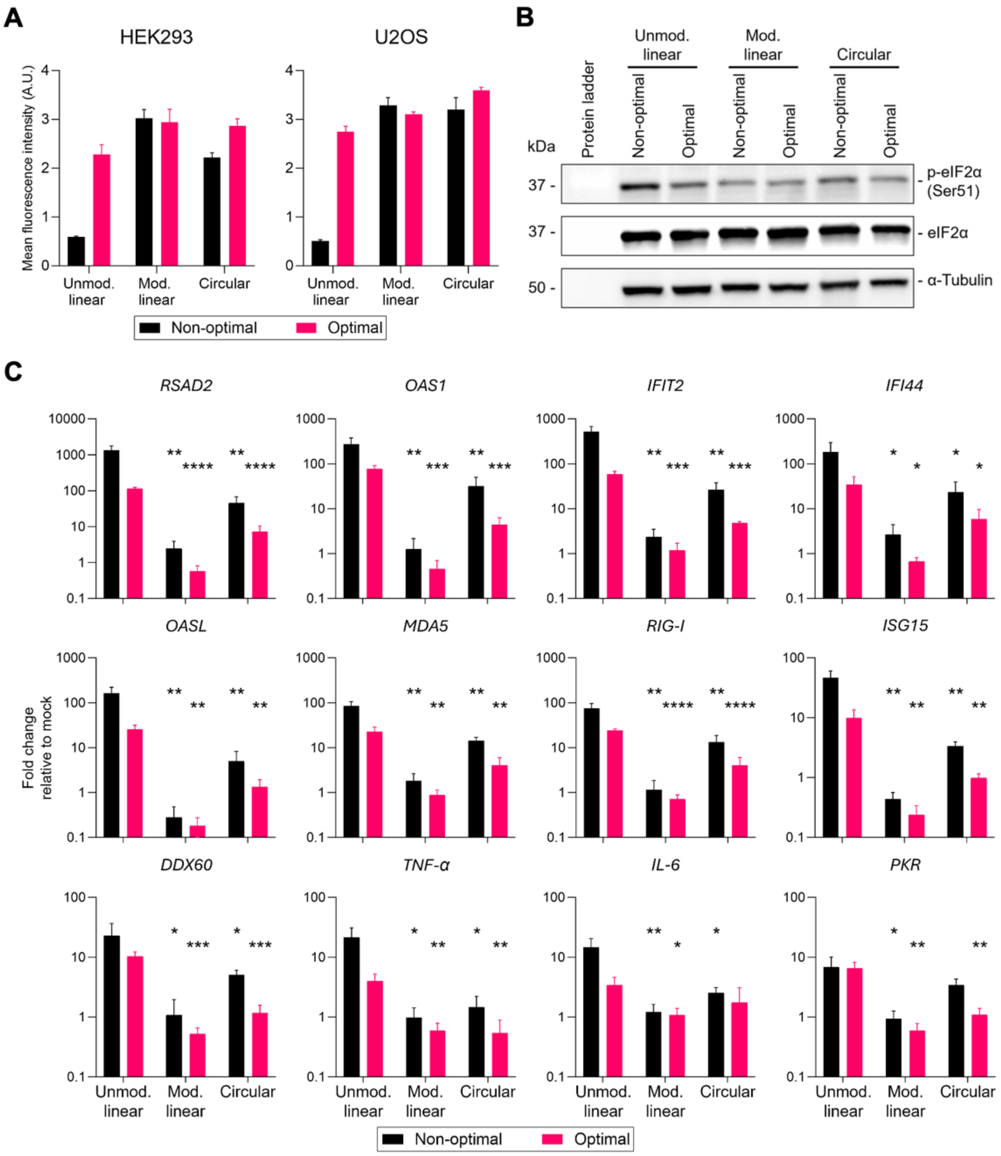


**Figure S5. Modified nucleosides and RNA circularization reduce eIF2 phosphorylation and innate immune gene activation. (A)** Mean green fluorescence intensity of HEK293 or U2OS cells 24 h after transfection with unmodified linear, modified linear, or circular mRNAs expressing non-optimal (black) or optimal (pink) *Renilla* luciferase reporter along with a green fluorescent protein transfection control mRNA. Fluorescence images were taken by an Incucyte S3. Error bars represent Std Dev of four independent replicates with five images taken per replicate. **(B)** Immunoblots were used to assay the phosphorylation of eIF2α in HEK293 cells 16 h after transfection with unmodified linear, modified linear, or circular mRNAs expressing non-optimal or optimal *Renilla* luciferase reporter. Total eIF2α and α-tubulin are used as loading controls. **(C)** U2OS cells were individually transfected with non-optimal or optimal *Renilla* luciferase reporter-encoding unmodified linear, modified linear, or circular RNAs for 16 h. Relative expression of innate immunity genes are measured by RT-qPCR, with relative fold change normalized to expression of control (mock) transfection. Data are shown as the mean of three independent replicates where error bars represent Std Dev. $*p < 0.05$, $**p < 0.01$, $***p < 0.001$, $****p < 0.0001$ compared to unmodified linear with identical coding sequence when using Student's t-test.

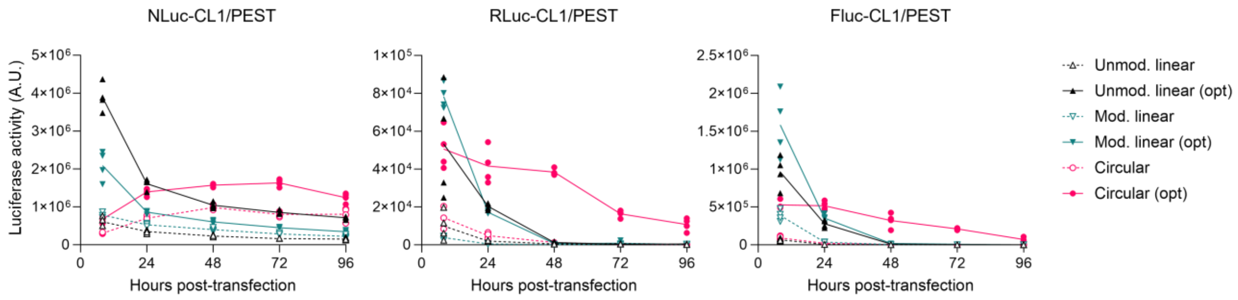


**Figure S6. RNA circularization increases the lifespan of optimal reporter transcripts in HEK293 cells.** HEK293 cells were co-transfected with a transfection control and either unmodified linear, modified linear, or circular mRNAs encoding non-optimal or optimal CL1/PEST degron-tagged luciferase reporters. Protein expression was measured by the corresponding luciferase assay at 8, 24, 48, 72, and 96 h after transfection. The data are presented as a time-course of luciferase activity from 8 h to 96 h of four independent replicates.

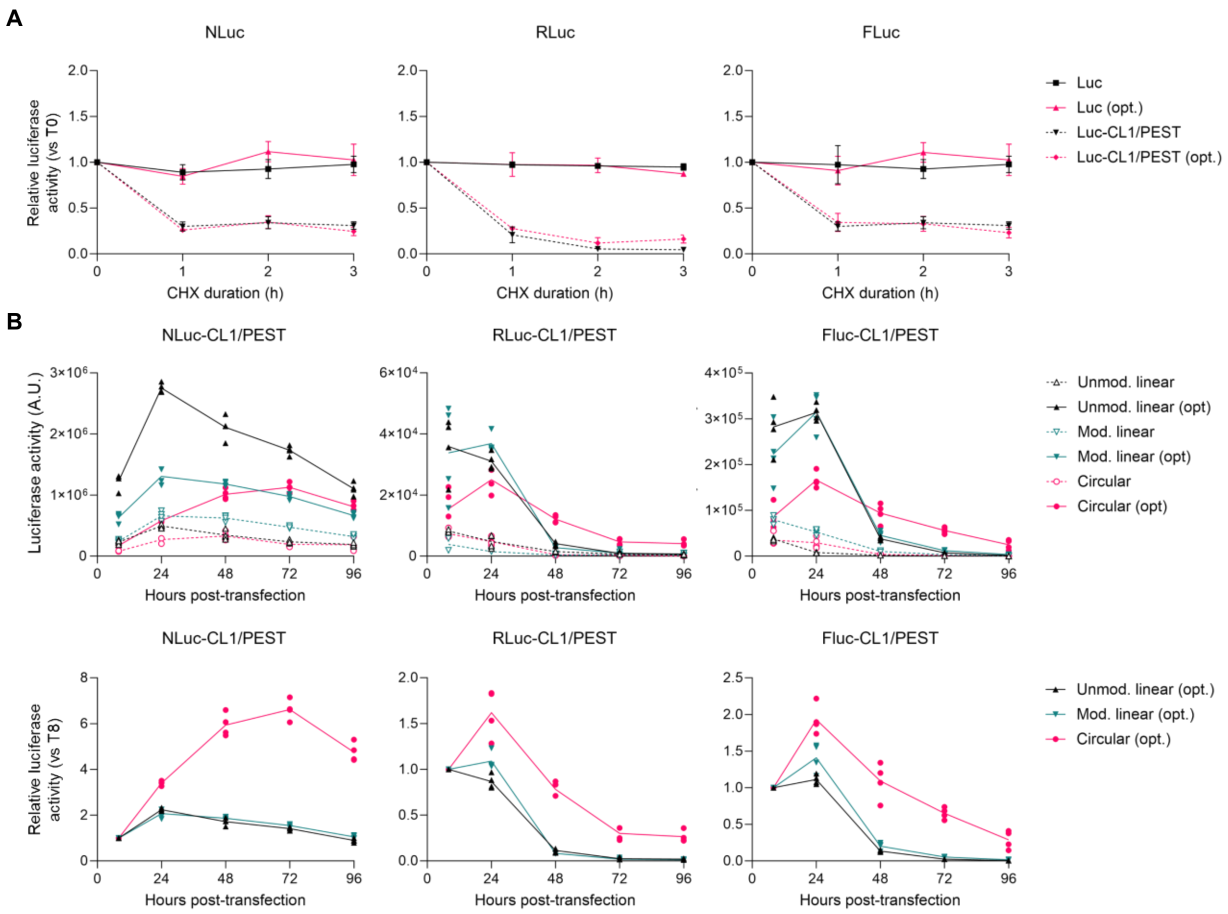


**Figure S7. RNA circularization increases the lifespan of optimal reporter transcripts in U2OS cells. (A)** U2OS cells were co-transfected with linear mRNAs encoding a transfection control and the indicated reporters for 16 h. Luminescence signals were measured and normalized to the co-transfected control at the indicated time points after addition of cycloheximide (CHX, 100 µg/mL). The data shown for each reporter are presented as the mean ±Std Dev of four independent replicates and were compared to the time point when CHX was added (0 h). **(B)** U2OS cells were co-transfected with a transfection control and either unmodified linear, modified linear, or circular RNAs encoding non-optimal or optimal CL1/PEST degron-tagged luciferase reporters. Protein expression was measured by the corresponding luciferase assay at 8, 24, 48, 72, and 96 h after transfection. The data are presented as a time-course of luciferase activity from 8 h to 96 h (top) and as a relative luciferase activity (RLU) (bottom) of four independent replicates and were compared to the signal observed 8 h after transfection.

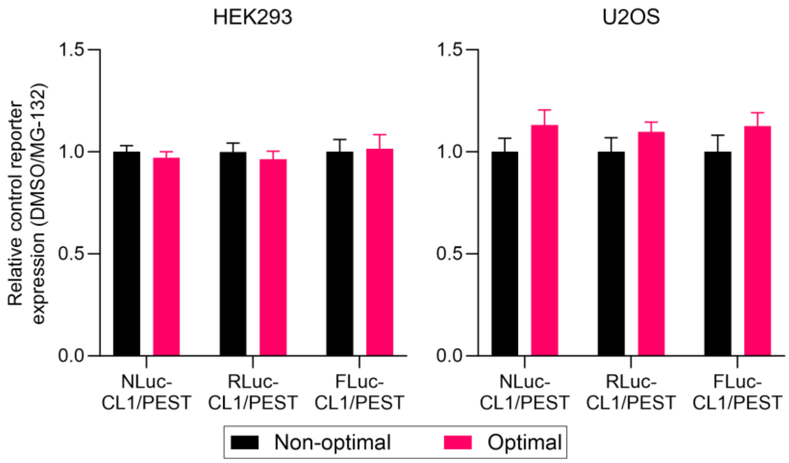


**Figure S8. Codon optimality of test mRNAs does not affect proteasome-mediated degradation.** HEK293 or U2OS cells were transfected with unmodified linear mRNAs encoding control reporters. After 16 h, cells were subsequently transfected with either non-optimal or optimal CL1/PEST degron-tagged reporters. Cells were then treated with DMSO or 10 µg/mL MG-132 to prevent proteasome-mediated degradation, and luminescence signals were measured 6 h later. The luminescence signals of DMSO-treated cells were normalized to those of MG-132 treated cells. The data shown for each reporter were set relative to the non-optimal reporter and are shown as mean ± Std Dev of four independent replicates.
